# A multiscale analysis of early flower development in Arabidopsis provides an integrated view of molecular regulation and growth control

**DOI:** 10.1101/2020.09.25.313312

**Authors:** Yassin Refahi, Argyris Zardilis, Gaël Michelin, Raymond Wightman, Bruno Leggio, Jonathan Legrand, Emmanuel Faure, Laetitia Vachez, Alessia Armezzani, Anne-Evodie Risson, Feng Zhao, Pradeep Das, Nathanaël Prunet, Elliot Meyerowitz, Christophe Godin, Grégoire Malandain, Henrik Jönsson, Jan Traas

**Affiliations:** The Sainsbury Laboratory, University of Cambridge, Bateman Street, Cambridge, CB2 1LR, UK; Laboratoire RDP, Université de Lyon 1, ENS-Lyon, INRAE, CNRS, UCBL, 69364 Lyon, France; Université Côte d’Azur, Inria, CNRS, I3S, France; Division of Biology, California Institute of Technology, Pasadena, CA 91125, USA; Howard Hughes Medical Institute and Division of Biology and Biological Engineering 156-29, California Institute of Technology, Pasadena, CA 91125, USA; Computational Biology and Biological Physics, Lund University, Sölvegatan 14A, 223 62 Lund, Sweden; Department of Applied Mathematics and Theoretical Physics (DAMTP), University of Cambridge, Cambridge, UK; Université de Reims Champagne Ardenne, INRAE, FARE, UMR A 614, 51097 Reims, France

## Abstract

The link between gene regulation and morphogenesis of multicellular organisms is a fundamental problem in biology. We address this question in the floral meristem of *Arabidopsis*, which generates new tissues and organs through complex changes in growth patterns. Starting from high-resolution time-lapse images, we generated a comprehensive 4-D atlas of early flower development including cell lineage, cellular growth rates and the expression patterns of 28 regulatory genes. This information was introduced in MorphoNet, a web-based open-access platform.

The application of mechanistic computational models indicated that the molecular network based on the literature only explained a minority of the expression patterns. This was substantially improved by adding single regulatory hypotheses for individual genes. We next used the integrated information to correlate growth with the combinatorial expression of multiple genes. This led us to propose a set of hypotheses for the action of individual genes in morphogenesis, not visible by simply correlating gene expression and growth. This identified the central transcription factor *LEAFY* as a potential regulator of heterogeneous growth, which was supported by quantifying growth patterns in a *leafy* mutant. By providing an integrated, multiscale view of flower development, this atlas should represent a fundamental step towards mechanistic multiscale-scale models of flower development.

## Introduction

The loss of function of many regulatory genes causes important perturbations in the growth patterns of multicellular organisms, which means that they directly or indirectly affect local growth parameters via the expression of other genes or physical cell properties. The regulatory networks and their dynamics have been extensively studied in a range of model species (e.g. (Briggs et al., 2018) (Chen et al., 2018) (Wagner et al., 2018)). This has been an active field of research, in particular with the advent of single cell sequencing methods, which can now be combined with molecular cell lineage tracking approaches (e.g. (Cotterell et al., 2020; Frieda et al., 2017)). However, there is often only a partial view of how growth is coordinated and as a result gene function is usually expressed in general terms such as organ identity or polarity, referring to their main mutant phenotype in a relatively abstract and qualitative manner. In addition there are still many open questions regarding the regulatory network structures and it is often impossible to test their coherence. An important first step towards addressing these problems is to integrate the existing information on gene expression, and to quantitatively correlate regulatory inputs, for example in the form of gene expression patterns, with the final output, i.e. shape changes during development (Coen et al., 2004; Whitewoods and Coen, 2017). This should then provide a solid basis for more mechanistic studies, involving also the regulation of biochemical interactions and biophysical aspects (Abad et al., 2017; Diaz de la Loza and Thompson, 2017; Pasakarnis et al., 2016; Thompson, 1917; Zhu and Roeder, 2020).

We address this question in the floral meristem (FM) of the model plant *Arabidopsis*, which generates four whorls of floral organs and is one of the best-characterized morphogenetic systems available (Blázquez et al., 2006; Bowman et al., 2012; Chen et al., 2018). The function of a range of key genes together with their domains of expression has mainly been studied on a one by one basis and their individual function, spatial expression and dynamics have been characterized (Fig. 1, supplementary table 1). Like the vegetative and inflorescence meristems, the floral meristem contains a population of stem cells, which are kept in an undifferentiated state by regulatory genes like *SHOOTMERISTEMLESS* (*STM*), *WUSCHEL* (*WUS*) and *CLAVATA* (*CLV*) *1-3* ((Long and Barton, 2000); (Lenhard and Laux, 2003); (Mayer et al., 1998)). Other genes, like *AINTEGUMENTA* and *MONOPTEROS* have been more specifically associated with organ outgrowth ((Krizek, 2009); (Nole-Wilson and Krizek, 2006); (Yamaguchi et al., 2013)). During flower formation, yet another set of regulators, including *PHAVOLUTA, PHABULOSA, ASYMMETRIC LEAVES1* and *2, FILAMENTOUS FLOWER* and *ETTIN*, determines the abaxial/adaxial (dorso-ventral) polarity of the organs, i.e. the identity of the cells next to and further away from the shoot meristem ((Emery et al., 2003); (Iwakawa et al., 2007); (Machida et al., 2015; Sawa et al., 1999a); (McConnell et al., 2001); (Sawa et al., 1999b); (Sessions et al., 1997)). The floral organs are separated by boundary domains characterized by the expression of notably the *CUP SHAPED COTYLEDON* (*CUC*) *1-3* genes ((Aida et al., 1997); (Hibara et al., 2006)). The spatial and temporal regulation also involves several hormones, including cytokinin and auxin ((Besnard et al., 2014a) (Reinhardt et al., 2003)). Cytokinin has been mainly associated with meristematic activity, whereas auxin is required for organ positioning and outgrowth. While the previous regulators can also be found in vegetative meristems, a major subnetwork, including the transcription factors L*EAFY* (*LFY*), *APETALA* (*AP*) *1-3, PISTILLATA* (PI), *AGAMOUS* (AG) and *SEPALLATA* (*SEP*) *1-4*, is involved in defining the type of organs to be produced ((Blázquez et al., 2006; Krizek and Fletcher, 2005); (Goto and Meyerowitz, 1994); (Kaufmann et al., 2009); (Ó’Maoiléidigh et al., 2014); (Parcy et al., 1998); (Pelaz et al., 2000); (Wuest et al., 2012); (Thomson and Wellmer, 2019)).

**Figure 1.**
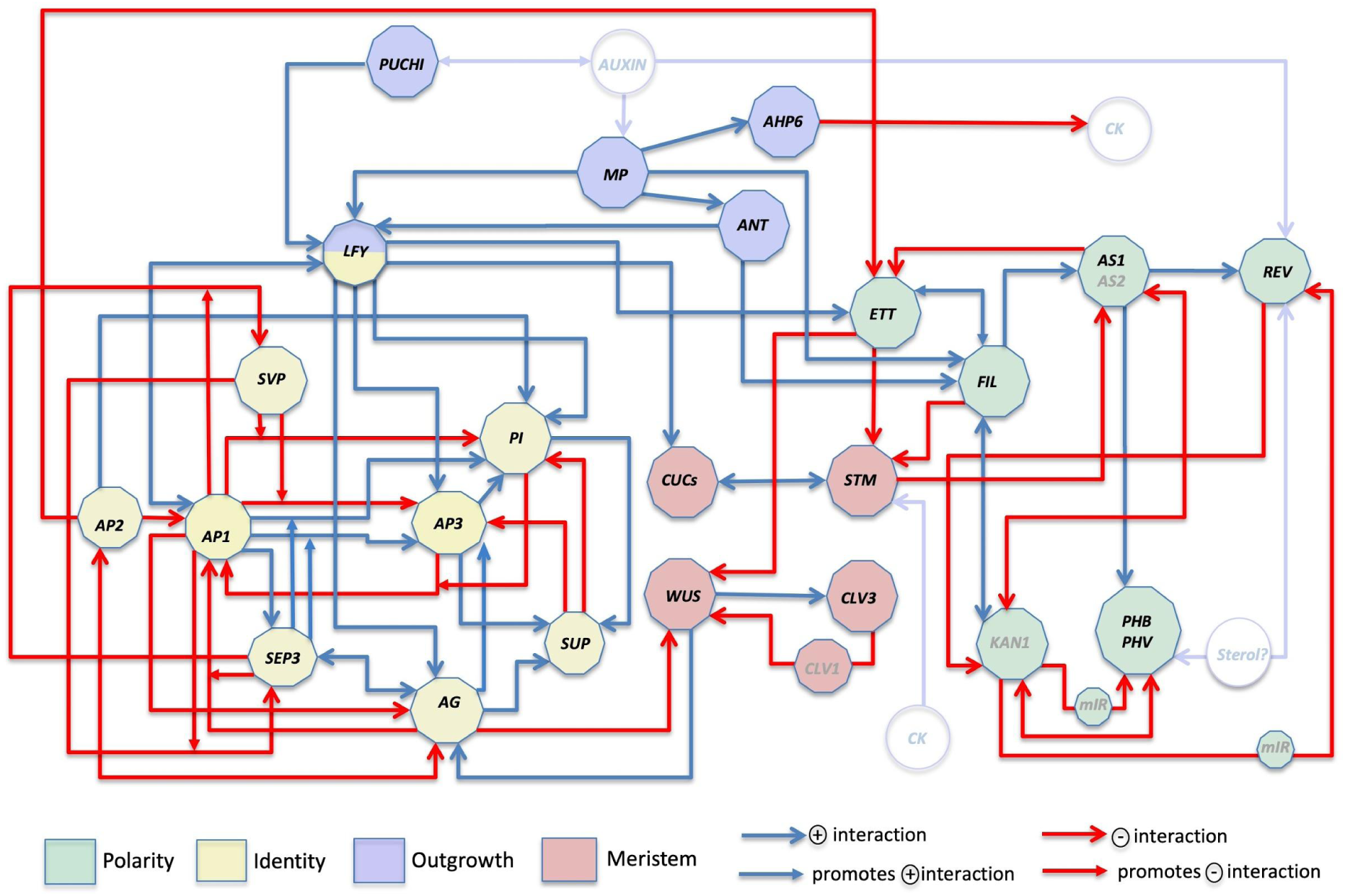
Gene regulatory network proposed for flower patterning and morphogenesis. Red and blue connections represent negative and positive regulations respectively. Colour code indicates function in floral meristem development, floral organ identity (sepals, petals, stamens and carpels) and abaxial/adaxial organ polarity as described in the litterature. The regulators in light blue characters are not recorded in the current version of the Atlas because there is not sufficient information on their expression patterns.

The general architecture of this network and parts thereof have been studied and models for molecular regulation have been proposed (e.g.: (Sánchez-Corrales et al., 2010); (La Rota et al., 2011); (Chen et al., 2018). However, in spite of this extensive body of knowledge, our understanding of how the network orchestrates flower morphogenesis remains fragmentary. The coherence of the existing data needs to be tested, while a more integrated, multiscale view at the level of the whole system is missing. This not only implies a need for a better knowledge of how the network is behaving in time and space at cellular resolution. In addition the network dynamics need to be correlated with growth patterns.

We used high-resolution time-lapse images to generate a comprehensive 4-D atlas of early flower development, including the expression patterns of 28 regulatory genes. This integrated view allowed us to test the coherence of the published data on molecular regulation. A quantitative correlation analysis between gene expression patterns and growth patterns then led us to propose a set of hypotheses for the combinatorial action of regulatory genes in patterning and morphogenesis. Hypotheses concerning the central regulator *LEAFY* were tested experimentally, which supported a role in growth control both during sepal initiation and organ boundary formation.

The results were made available in the form of an interactive web-based atlas using a dedicated online tool called ‘Morphonet’ (http://morphonet.org, (Leggio et al., 2019)) that can be accessed and further developed by the entire scientific community.

## Results

### High-resolution live imaging of flower development reveals consistency in shape and size

We used confocal microscopy to live image flower primordia from initiation to stage 4 when the sepals start to overlie the flower meristem and all four whorls have been specified (Smyth et al., 1990). This was done using a yellow fluorescent protein (YFP) targeted to the plasma membrane or using the membrane specific dye FM 4-64 (see Methods, (Fernandez et al., 2010; Willis et al., 2016)).

The development of six meristems (FM1-6) was recorded in a total of 50 3-D image-stacks, followed by cell segmentation and lineage tracking (Methods, Fig 2, Fig S1). In flower meristems, cell division patterns are not fixed, in contrast to e.g. roots or hypocotyls (Montenegro-Johnson et al., 2015). It is, therefore, not straightforward to compute an average time course, including cell lineage information which is essential to study the correlation between cellular dynamics and gene expression. For this reason we aimed to select a representative series for further analysis. To this end, we compared the shape of all 6 meristems during development. Since there is no obvious way to synchronise flower development and the geometrical shape of individual time points of one series does not exactly correspond to the time points of other series. To circumvent these problems, we applied a registration method to align and compare different acquisition sets using the surface of flower primordia, represented by a point cloud, as the overall shape mesure (Methods, (Michelin et al., 2016)) The quantitative assessment of the variability in shape and size of the flower primordia captured in sequences FM1-4 and FM6 illustrated that they go through similar developmental stages with consistent shapes and sizes (Fig. S2 and S3), while it was not possible to compare the shape of FM5 reliably with that of the other meristems (Fig. S3). This motivated the choice of FM1, which had the highest temporal resolution and spanned floral development from initiation to stage 4, as a representative reference. To facilitate further analysis, this time series was added to the web-based browser MorphoNet (http://morphonet.org/, Methods, (Leggio et al., 2019)).

**Figure 2.**
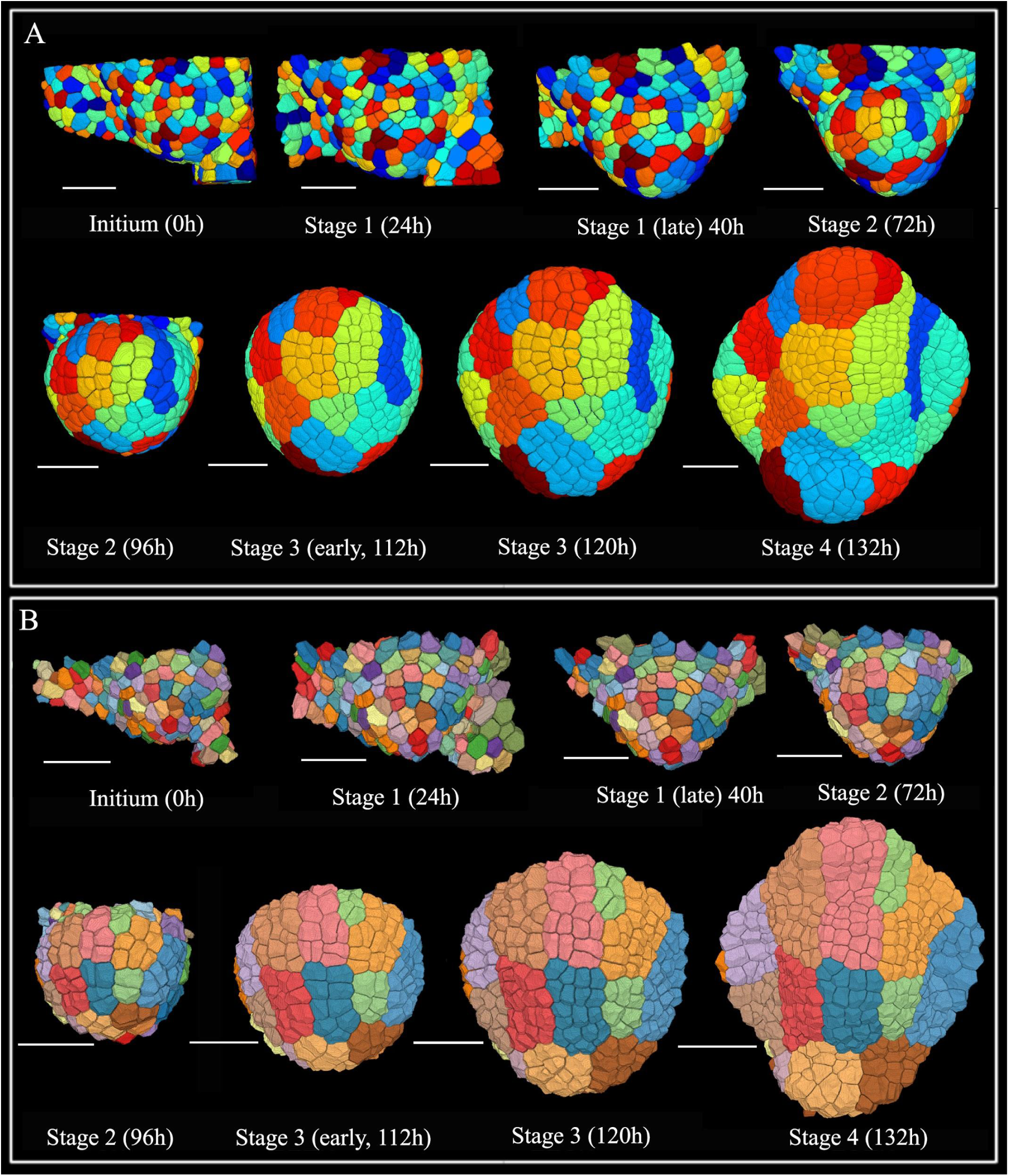
Cell lineage in reference series. 8 out of 18 timepoints of FM1 are shown. Times after first acquisition are indicated between brackets. (**A**) Surface rendering of segmented images of time course 1 showing L1 layer cells. (**B**) Surface rendering of segmented images showing L2 cells (L1 cells are removed. Cells are colored according to computed lineages. Scale bars in A and B 20 um

### An integrated view of gene expression patterns provides a high-resolution spatiotemporal differentiation map of flower development

We next included the expression patterns of 28 important genes involved in floral meristem function, organ identity, organ outgrowth and organ polarity in the 4-D template (Figs. 1, 3A,B). For 21 genes, the often partial published information was complemented by our own results coming from RNA in situ hybridization or confocal live-imaging (Methods). The collected patterns were integrated into the FM1 time course by manually annotating individual cells using the tools available via MorphoNet (Methods, Figs S4-S6, Supplemental file for justification of individual genes). For this purpose 5 distinct stages of development were chosen (as defined by (Smyth et al., 1990)): the initium stage (called here stage 0) and stage 1 to stage 4 (Fig 3B). We chose to perform a binary labelling, i.e. to indicate only the presence or absence of gene expression, given the predominant qualitative nature of the available expression data. The single gene expression patterns (***‘gene patterns’*)** were then combined. We could thus identify cell groups that expressed unique gene combinations and corresponded to specific differentiation states, termed ***‘cell states’***. 28 cell states were present in both L1 and L2 (not taking into account the L1 marker ATML1) and 3 L2-specific states were found (Fig. 3). We next conducted an exploratory data analysis using unsupervised hierarchical clustering, (Methods), to generate a cell state similarity map using Hamming distances, i.e. the number of gene expressions which differ between cell states, as the measure of similarity (Fig. 3C-D). The dendrogram revealed clusters of states, which formed different functional groups of cells through flower development (Fig. 3D, Supplementary Table 2). These included, for example, meristematic cells, boundary cells, as well as cells expressing genes defining polarity or organ primordia. Similarities in expression between alternative cell fates were identified at high resolution, e.g. connecting the boundary domain with the expression domains where petals and anthers are initiated (Fig 3D: boundary cluster, containing states 4, 20, 30, 19, 25 and 29). To investigate the temporal evolution or cell-differentiation paths of the cell states in the outer cell layer, we used the computed cell lineages and built a weighted directed cell state **‘transition graph’** where the nodes are the cell states (Fig. 4, Methods). The graph reveals a core of ‘stem cells’ at the adaxial domain of the bud at stage 1 (State 7), providing all cell types of the flower at stage 4. This is similar to typical tree-like differentiation paths often described for mammalian development (Enver et al., 2009). Similarly, at stage 2, State 7 has split up in sepal ‘precursors’ and the central meristematic domain (State 10), which will give rise to all other states. However, the plasticity of plant cells is clearly represented where several cell types contribute to future stages. For example, sepal tip cells come from all cell states present at stages 0 and 1. Descendants of both bract (State 5) and SAM boundary cells (State 4) contribute to the same cell state at later stages, although being quite different in terms of their original expression patterns.

**Figure 3.**
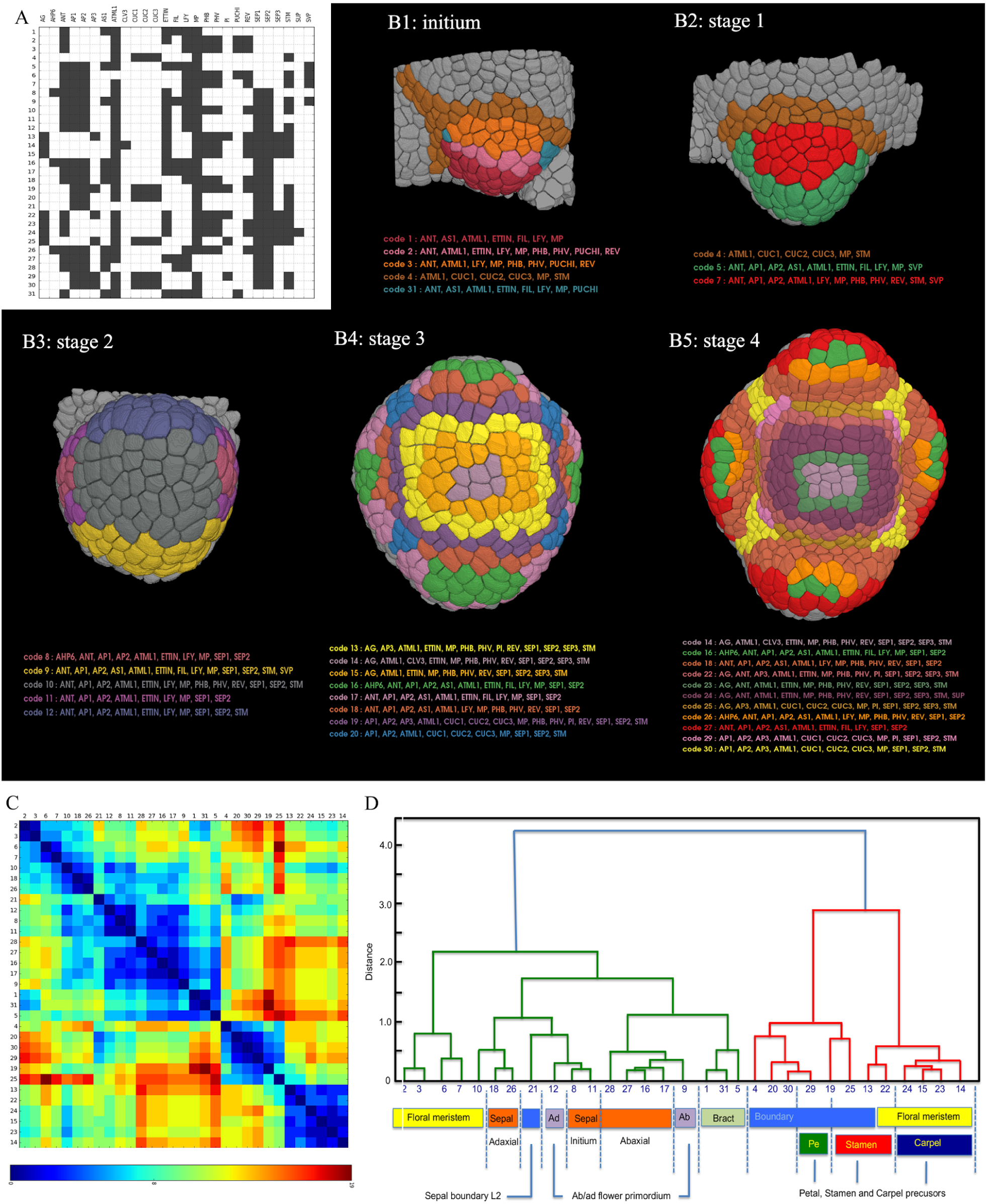
(**A**) Matrix representing 32 combinatorial, binary expression patterns (referred to as ‘states’) of 28 genes. *WUS*, which is only expressed in internal tissues, is not represented. Each row corresponds to a particular state (state numbers are given on the left) and contains black (gene active) and white (gene inactive) squares. Names of the genes are given on top of each column. (**B**) Rendering of gene expression patterns in the L1 layer. Each of the patterns are colored by a unique color and their corresponding codes are also given. B.1., B.2., B.3., B.4., and B.5 correspond respectively to intium, stage 1, stage 2, stage 3, and stage 4 time points. Grey cells have not been annotated (no expression). (**C**) Hierarchical clustering of cell states using Hamming distances as the measure of similarity. The heat map (similarity matrix) corresponds to the Hamming distances and the columns and rows are the patterns. D) Individual clusters or combinations of clusters correspond to specific differentiation domains (organ identity for example) in the growing flower (color coded). Alternatively, the clusters can be assigned to more general ‘functional’ domains not specific for the flower (meristem, boundary domains).

**Figure 4.**
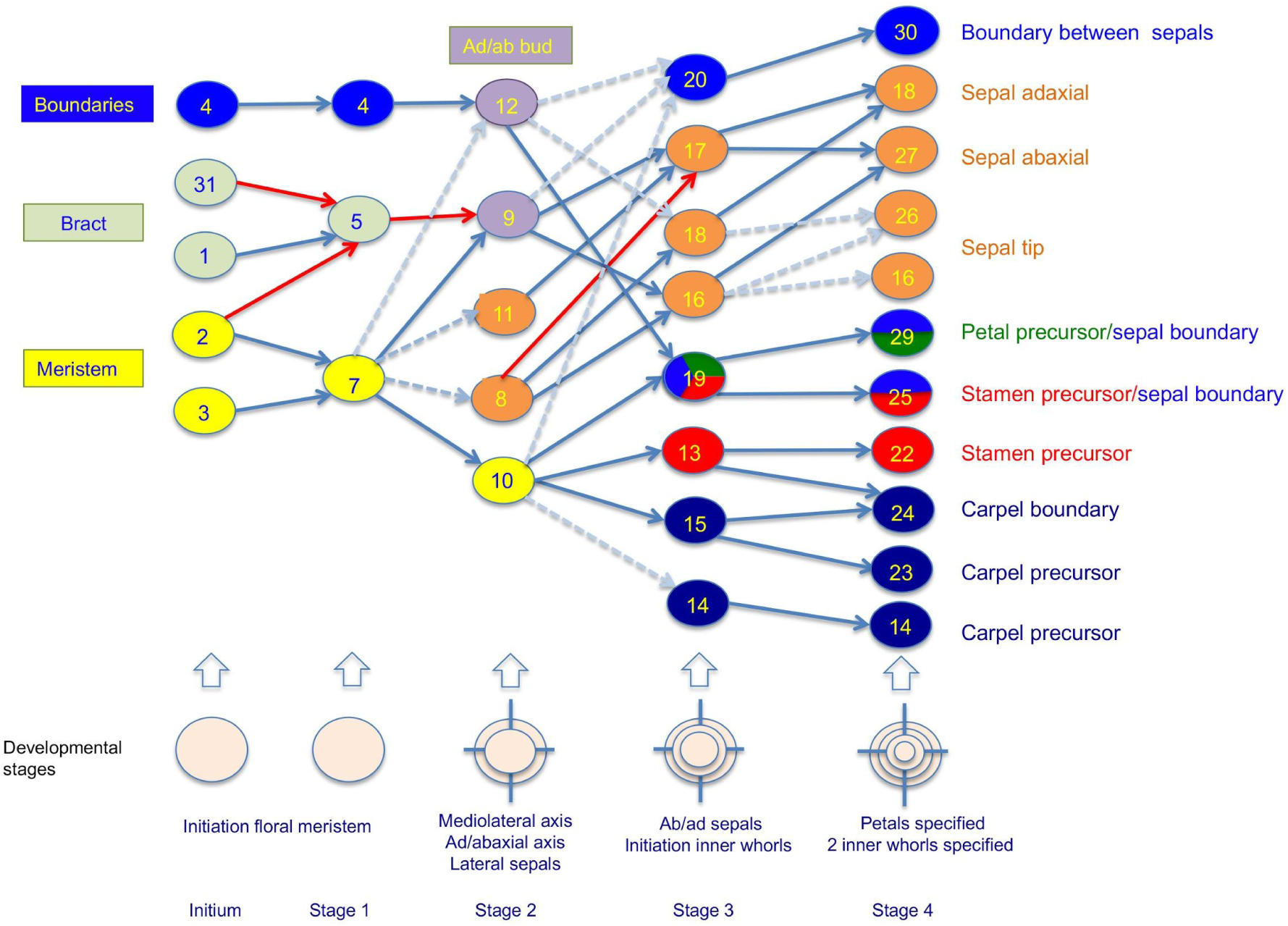
Temporal evolution of clusters and cell states. The graph combines ‘forward’ and ‘backward’ links. Forward links connect specific states at one time point to their descendant patterns in the next time with weights corresponding to the fractions of the daughter cells in each of the descendant patterns. Similarly ‘backward’ links identify states at a previous time point from a current one. Arcs with blue arrows indicate the presence of corresponding arcs both in forward and reverse pattern transition graphs, dashed arrows indicate the presence of corresponding arcs in reverse pattern transition graph only, red arrows indicate the presence of corresponding arc forward patterns transition graph only (see also supplementary figures). The links whose weights were below a threshold of 20% were pruned.

### Adding single regulatory hypotheses for individual genes substantially improves gene pattern predictions

We next examined alternative hypotheses to quantitatively explain these gene expression patterns (Fig. 5). We first analyzed the possibility of the patterns being driven mainly by the lineage, i.e. whether the gene expression at an earlier time point can be used to predict the expression at a later time point. This led to relatively good predictions at early and late stages, but was less successful for transitions between intermediate stages (Fig. 5B-C), as measured by a *Balanced Accuracy* score (BAcc), combining the normalised false positive and false negative rates (Methods). This indicates that most regulatory interactions have an effect during the end of stage 2 and stage 3. For example, for over 75% of the genes we found a BAcc score larger than 0.75 (closer to a perfect pattern than to a random pattern) when following lineages from stage 0 to 1 or from 3 to 4, while this was only true for less than 20% of the genes when cells were followed from stage 2 to 3 (Fig. 5B-C). The result was highly variable between genes and also between individual time points for single genes (Fig. 5B).

**Figure 5:**
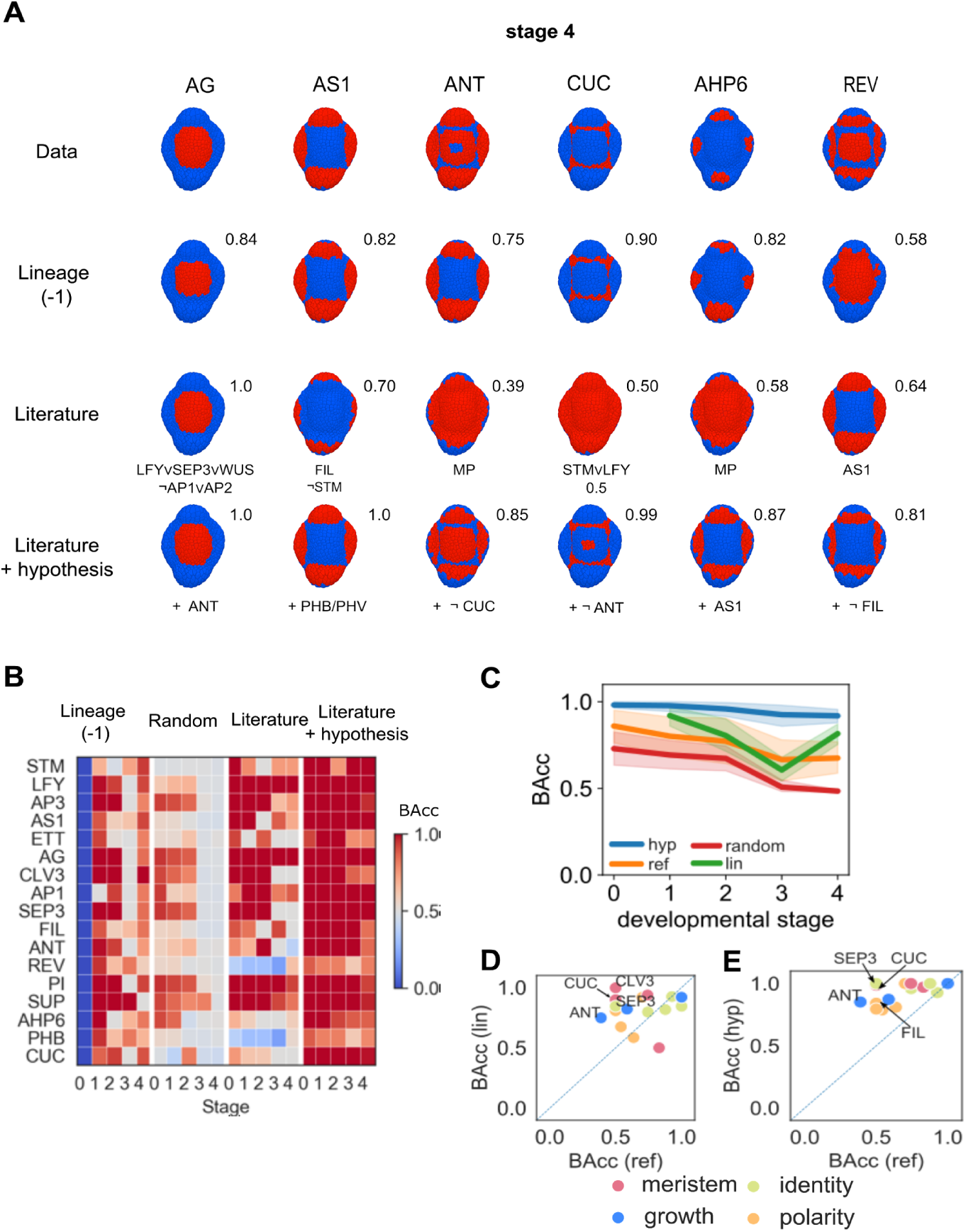
Expected patterns vs predicted patterns from various models. **(A)** Example gene expression patterns at stage 4 (132h). Data from the template (top row) is compared with predicted patterns coming from: Following cell lineages from stage 3 (second row), boolean regulatory model from literature network (Fig. 1, third row), and boolean regulatory model where an additional regulatory arrow has been added as hypothesis (Table S4, fourth row). Numbers above the illustrations indicate the BAcc score. Below the illustrations either the selected combination of inputs (in the case of the literature based boolean model) or the selected best hypothesis (in the case of the augmented literature boolean model) is shown. **(B)** BAcc score for all genes using the methods of lineage, literature network with randomised input (see Methods), literature network regulatory model (Fig. 1), and model with added hypotheses. Result of best hypothesis given for each Stage of flower development **(C)** Average (and standard deviation) of BAcc score over all time points and genes for the same conditions as described in (B). **(D)** Gene by gene comparison of BAcc scores between literature network (ref) and lineage (lin) at stage 4. **(E)** Gene by gene comparison of BAcc scores between literature derived network (Fig 1) and after adding regulatory hypothesis (hyp) at stage 4.

We next tested whether the literature-derived network (Fig. 1) could account for the expression patterns of single genes. To do this, we modelled the gene regulatory network using a set of boolean rules combining activating and repressing inputs suggested by the literature, and combined them in all possible ways using logical ‘AND’ and ‘OR’ rules (Methods). This produced a ranked list of alternative logical combinations for each gene regulation and the combination(s) with the best similarity with the gene expression pattern at hand were selected (Methods, Supplemental Table S3). For comparison, we also used the input regulatory arrows from the literature network but selected random genes as inputs (Fig. 5B-C, Methods). The literature-based regulatory network improved the ability to explain a number of gene expression patterns, in particular during the last two stages (BAcc increase of 0.22 on average for stage 3 and 4) compared to the randomized networks (Fig. 5B-C). Several genes, such as LFY and AG at all time points, AS1 at stages 1, 3-4, and FIL at stage 3 all map perfectly or almost perfectly on the expression patterns extracted from the literature, indicating that the regulation presented in the literature matches the patterns well for a subset of genes at certain time points (Fig. 5A-B). However, this is only true for a minority of genes in the network. While the literature network was, in average, always performing better than the randomized networks, it was only significant at the later stages 3 and 4 (p-values of 0.09, 0.19, 0.23 for stages 0, 1, 2, respectively compared to 0.04 and 0.008 at stages 3 and 4, respectively).

When comparing the literature based regulatory predictions with lineage predictions, the literature-based regulation did not lead to significant improvements on average at any time point, and lineage even performed better at stage 4 (Fig. 5D, p-values 0.11, 0.44, 0.46, 0.02 at stages 1, 2, 3, 4, respectively). At stage 4 for example, the literature-based network only improved the predictions for a small number of genes such as LFY, AG, and SUP, while the patterns of ANT, the CUC genes, and SEP3 were better explained by lineage (Fig. 5A-D).

For a large subset of the genes the expression could not be reproduced using the regulatory interactions provided in the literature nor using lineages, indicating that the regulatory network is not complete. We therefore investigated how the addition or removal of single regulatory inputs changed the ability to predict the spatial pattern of each individual gene. This approach identifies the most plausible extra inputs required to generate the pattern. After testing all additional genes as activators or repressors and all possible logical combinations with the literature-suggested regulation, we identified the regulations leading to the best BAcc score for each gene (Methods, Table S4). This increased the predictability of the gene patterns significantly as compared to using the literature-proposed regulation (p-values 0.012, 0.003, 0.004, <10^−3^, <10^−3^ for timepoints at stages 0, 1, 2, 3 and 4 respectively; Fig. 5A-C, E), and improved the pattern for all but two genes (Fig. 5E).

In summary, our results show that the gene regulatory network provided by the scientific community only significantly improved the predictability of gene expression patterns compared to random interactions at late time points. Compared to cell lineage, the published network improved predictability for a small subset of the genes at specific stages of development only. Significant improvements in predictability were achieved for gene patterns by adding novel single interactions. When combined, the added hypotheses represent a plausible coherent mechanistic description of a gene regulatory network that can explain the gene patterns for early flower development.

### Exploring the genetic control of growth patterns

We next investigated the control of growth during development. We therefore first computed the cellular properties. The distribution of cell sizes in this meristem was in line with previous studies on Arabidopsis meristems (Fernandez et al., 2010; Gibson et al., 2006, 2011; Jackson et al., 2019; Willis et al., 2016) (Fig S7), and the number of cell neighbors within the L1 (epidermal) and L2 (subepidermal) layers was in line with previous studies on plants and animals, (Lewis, 1926) (Gibson et al., 2006, 2011; Jackson et al., 2019; Willis et al., 2016) (Fig S8).

Using cell lineage information, we computed growth rates and growth anisotropy at cellular resolution (Fig. 6, Methods). Since absolute expansion rates can fluctuate considerably between individual flowers and throughout development (Figs. S1 and S2), we focused on relative growth rates and directions between consecutive time points (Figs. 6A). Relative differences in growth rate were particularly striking at stage 3 and 4, when the sepals start to grow out (Fig. 6A, (Smyth et al., 1990)). The analysis of growth directions showed that cells start to grow anisotropically when the sepals are initiated. This was particularly evident on the abaxial side at stage 4, when these organs begin to cover the flower meristem (Fig 6B).

**Figure 6.**
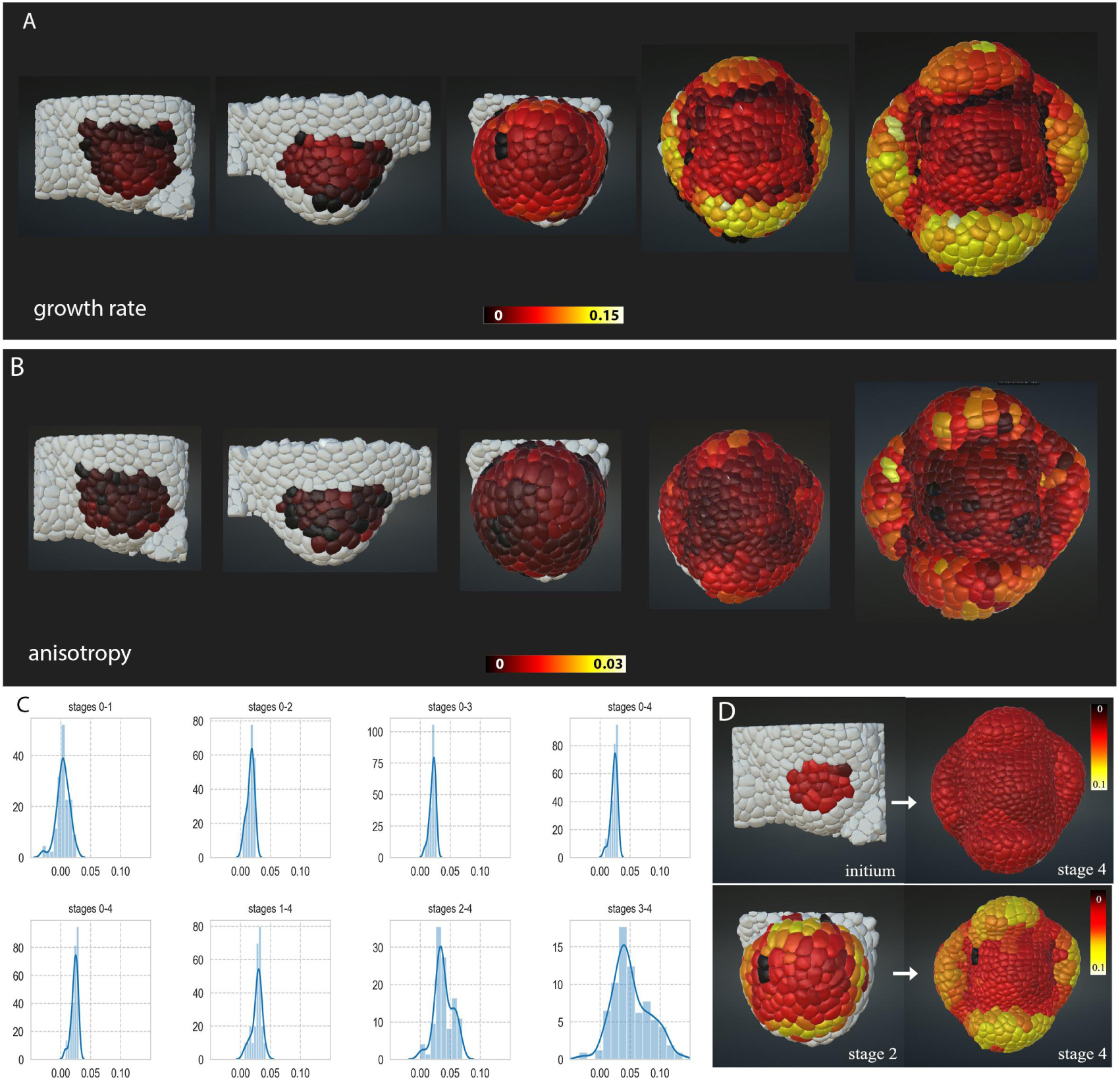
Relative growth rates for single cell clones. **(A)** Relative volumetric growth rates per hour of L1 cells, indicating how much the cells have grown. Color scale bar from nongrowing (black) to more rapidly growing cells (yellow/white) in um3/h. L1 cells with positive, relative growth rates are displayed. Light grey cells are not taken into account. **(B)** Growth anisotropy of L1 cells. Color scale indicates degree of anisotropy per hour (see methods). Light grey cells are not taken into account. (**C**) Distribution of growth rates (um3/h vs number of cells) between different time points. Note that the number of cells followed between timepoints can differ, depending on the available, tracked lineage **(D)** Relative volumetric growth rates per hour of L1 cells between 10-132h (upper panel) and 96 and 132h (lower panel). Color code on initium and stage 2 indicates how much the cells will grow. Color code on stage 4 how much the cells have grown. At 96h the cell lineage and increased growth rates of the sepals are already largely fixed. Two cells have not grown between stage 2 and 4, so their contribution must have been taken over by neighboring cells. **Note: growth rates and degree of anisotropy are taking into account both forward and backward rates (i**.**e. how much the cells have grown and how much they will grow). This is not the case for the first and last point, where resp only forward and backward growth is presented**.

Although all of the genes considered here are involved in growth regulation at some level, it is unknown where, when and to what extent they regulate growth rates and directions. To obtain further information on their roles, we next correlated the growth patterns with gene expression. To quantitatively compare the growth rates of different cell populations we introduced a **‘*relative growth difference*’ (RGD**) defined as (g_1_-g_2_)/(g_1_+g_2_), where g_1_ and g_2_ are the median growth rates within the two populations (Methods). A limited number of individual genes can be consistently connected to low or high relative growth rates (Fig. 7A). The gene expressed in cells with the highest median growth rate is AHP6 (RGD >0.07 and p-value < 0.001 when compared with all other genes). This gene has been linked to organ initiation (Besnard et al., 2014b), and is never connected to slowly growing cells. Conversely, CUC1-3 expressing cells are correlated with slow growth (RGD < 0.4, p-value < 0.003). By contrast, most expression domains show very broad distributions of growth rates indicating no instructive information and several genes have a double-peak distribution.

**Figure 7.**
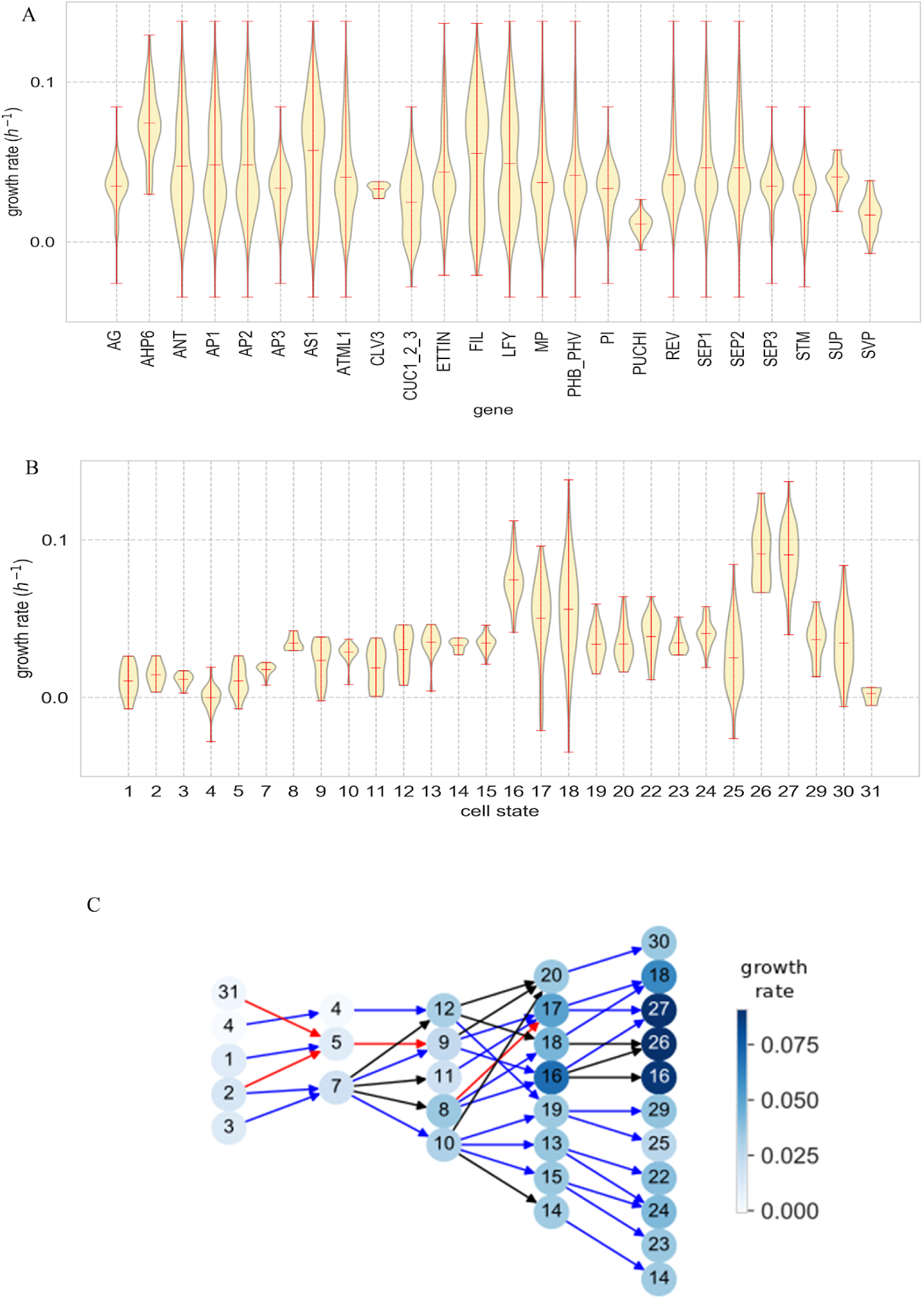
Growth rates correlated with gene expression and cell states. **(A)** Relative growth rates per hour in expression domains of individual genes, (B) Growth rates of combinatorial patterns or cell states, numbered as in Fig 4. Note that the growth rates were calculated from one point to the next point (forward), only the values of the last time point are calculated backwards (marked with *). (C) Growth on the pattern transition graph calculated as the average of the backward and forward rates, except initium stage, only forward and stage 4 only backward (arrows colored as in Fig. 5).

Cell ***states*** are better correlated with growth patterns with more spread between the growth distributions (Fig 7B, cf. Fig 7A). In particular, cell states 26 and 27 (RGD > 0.22, p-values <0.001 when compared with all other states) and their precursor states 16 and 17 (RGD > 0.1, p-values<0.003 when compared to states except 26 and 27) are fast-growing, identifying the central and abaxial side of the sepal (Fig. 3A, Figs. 6A, 7B-C). Whereas growth regulation is captured when using the cell state information, it is not trivial to determine the precise gene combination that provides the regulatory motif, in particular since the analysis of individual gene expression domains did not reveal any strong correlation. We therefore investigated growth correlations starting from pairwise comparisons of genes that had partial overlapping expression patterns (Fig. 8). For a pair of genes A and B, the idea was to see if the cells in the states expressing gene A were growing more slowly or more rapidly in the sub-set of states where they were co-expressed with gene B. This would identify gene B as having potentially a growth-promoting or inhibiting activity within the states where A is expressed. At floral stage 4, for example, STM identifies the slow-growing states 14, 22-24 within the ETTIN domain from the fast-growing states 16 and 27 (RGD = 0.36, Fig. 8A). Similarly, LFY identifies fast growing states within the AP1 domain (RGD = 0.36, Fig. 8A). This analysis was carried out for every gene combination and for every time point, which relates each individual gene expression pattern to all cell states where it has a differential expression (Fig. 8B, Figs. S12-S14). The genes fell into three broad classes: those that were potentially growth-stimulating, those that were potentially growth-inhibiting and a set that apparently had mixed effects (Fig. 8B). This confirmed AHP6 and CUC1-3 as potentially growth-promoting and inhibiting, respectively (cf. Fig. 7A). In addition, genes with very wide growth rate distributions, such as ANT and LFY came up as potential growth-promoting regulators (Fig. 8B).

**Fig 8.**
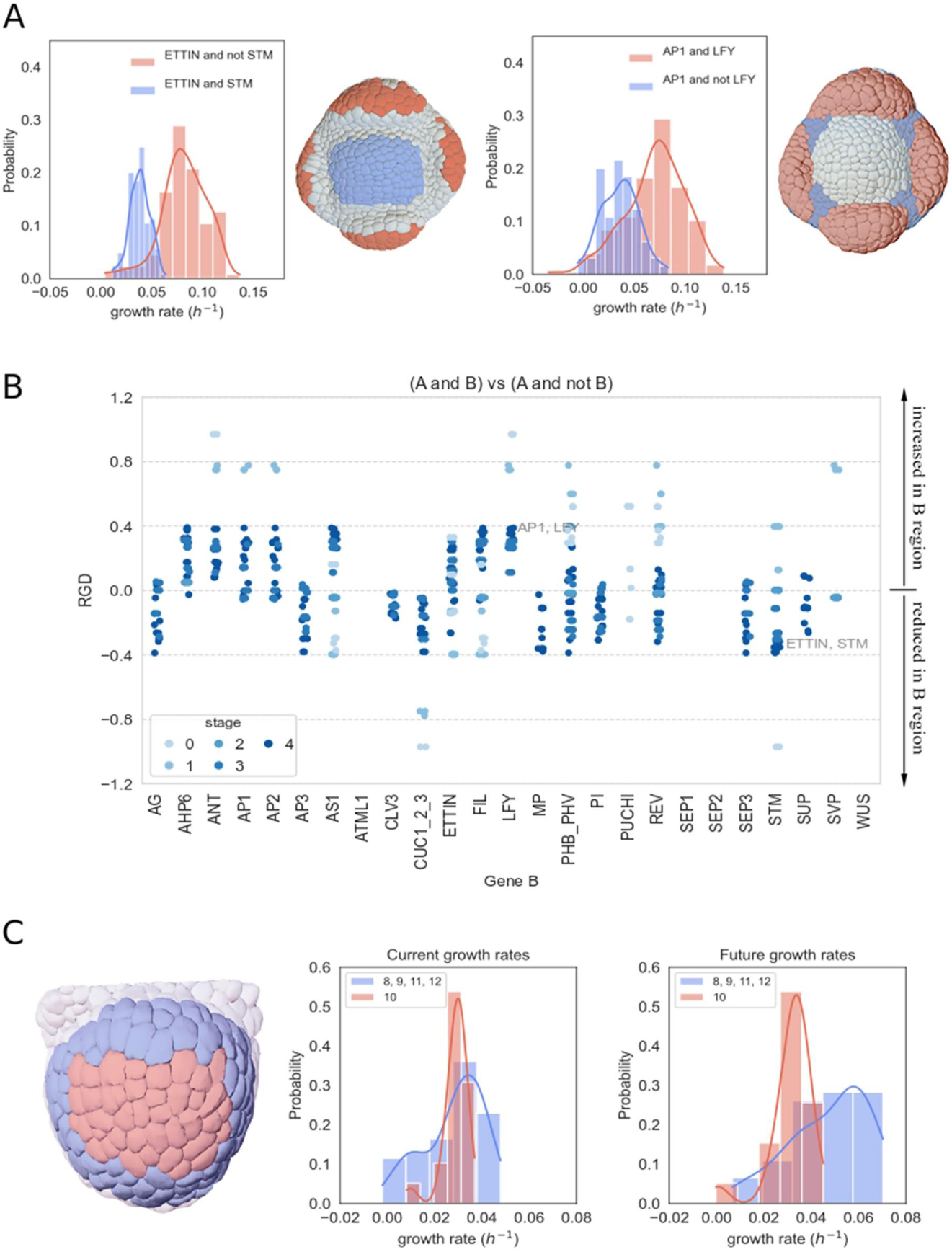
Analysis of growth rates in cell states and gene expression domains. (**A) and (B**.) Correlating growth with gene function. The growth of cells expressing gene A+ B is compared with the growth of those only expressing A. Two examples are given in **(A)** Within the *ETTIN* (gene A) expressing domain, the cells expressing STM (gene B) as well are growing more slowly; within the *AP1* domain, the *LFY* cells are growing more quickly. (**B)**. The results for all gene combinations. Values correspond to the RGD calculated using the median values of the growth distributions. Certain genes are mostly expressed in the more rapidly growing subdomains (e.g. *LFY, AHP6, ANT*), others mostly in the slowly growing subdomains (e.g. *SEP3, AG, CUC1-3*). The darker blue the spots correspond to later time points. **C**. Current and future growth rates of different domains at stage 2 of flower development. Current growth rates are relatively homogeneous in state 10 (orange zone) vs the other states (8, 9, 11, 12, blue zone). However the states in the blue zone will grow much more quickly afterwards, resulting in the outgrowth of the sepals.

In summary, apart from *AHP6* and *CUC1-3*, individual genes could not consistently be connected to relative growth rates and do not seem to act as dominant growth regulators by themselves. However, correlating the pairwise expression patterns of all 28 genes with growth patterns, we were able to propose growth promoting and/or inhibiting activities for a majority of them. Given that we also identify the combination of genes active in these regions (Figs. S12-S14), gene motifs for growth regulation are identified that can be included in mechanistic models.

With regard to the control of growth *directions*, the correlations with the individual gene expression patterns were not particularly informative. CLV3, PUCHI, SUP, SEP3 and SVP which showed low degrees of anisotropic growth (Fig. 9A), but many other genes were expressed in domains with relatively wide distributions. Like for growth rates, the cell states defined much more distinct behaviors. The cell states of the forming sepal are growing most anisotropically as identified in the transition graph (Figs. 6B and 9C), where the abaxial cells of the developing sepal (cell state 27) were also growing slightly more anisotropically than the adaxial side (cell state 16, Fig. 9B-C). This puts the polarity genes, in particular FIL, forward as potential regulators of anisotropic growth. Early time points have relatively low anisotropies, and there is a transition at stage 2, where the whole lineage coming from cell state 10 has relatively more isotropic growth compared to the sepal structures (Fig. 9C).

**Figure 9.**
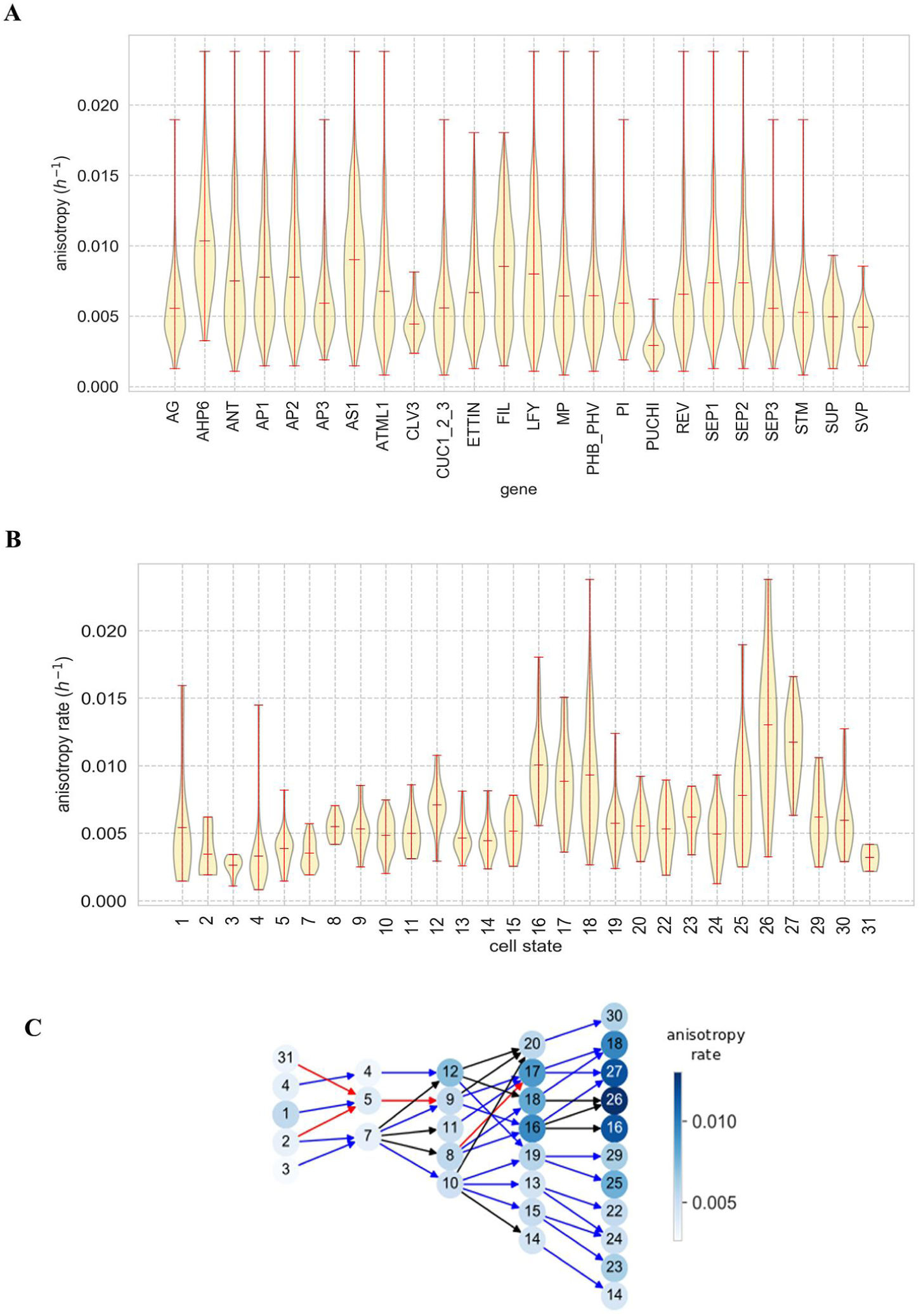
Growth anisotropy correlated with gene expression and cell states. **(A)** 3D growth anisotropy rate of expression domains of individual genes. **(B)** Growth anisotropy/hour in combinatorial patterns or cell states (numbering as in fig 4). Note that the values were calculated from one point to the next point (forward), only the values of the cell states of the last time point are calculated backwards (marked with *). **(C)** Growth anisotropy on the pattern transition graph calculated as the average of the backward and forward anisotropy rate (except initium stage, only forward and stage 4 only backward).

### Quantitative analysis identifies predefined growth patterns at stage 2

Growth analysis identified an increase in heterogeneous and anisotropic growth at later stages (Figs 6A,B 8). To understand when this switch can be first identified, we investigated growth over longer time scales. Considering the growth of cells from early stages (stages 0 and 1) onwards, no obvious spatial pattern of heterogeneity in the contribution to the final flower was found (Fig. 7D, Fig. S10A). This shows that at these early stages, the cells will produce a population of descendants of similar size at stage 4 or at least with no clear spatial correlation (Fig. 6D). However, at stage 2, although the flower bud is still close to a symmetric hemisphere (Figs. 2-3), cells have been committed to become fast or slow-growing at later stages (Fig. 6C, D), even if the growth rate at this time is quite homogeneous (Fig. 6A). The same observation was made for two other meristems with longer time series, FM2 and FM6 (Fig. S10 B,C).

This correlates also with the division of the state transition tree at stage 2, where the central state 10 is defining the central parts of the flower and are highly disconnected from the sepal differentiation lineages (Fig. 3). At stage 2 cell states 8, 9, 11 and 12 grow at approximately the same rates as the neighboring domain (cell state 10) (RGD <0.13, p-values > 0.753, Fig. 8C). Still, when comparing the growth up until stage 4, the descendants of these four cell states grow much faster (RGD>0.19, p-value <0.001, Fig. 9C) and hence identify growth precursor states (Figs. 7C, 8C, cf. Fig 6D). By contrast, cell state 10 has descendant states that are all relatively slow-growing (Fig. 7C).

In conclusion, we identified a transition from a patterning phase with homogeneous growth at stage 2 to a growth phase, where in particular the sepal lineages increase their growth rates and anisotropy.

### Quantitative growth analysis confirms a role of *LFY* in the coordination of growth between specific domains

Cells expressing *LFY* had one of the broadest growth distributions among all genes (Fig. 7A). Still, the more detailed analysis of the growth patterns described above, pointed at a prominent role in stimulating growth during early flower development. Whereas it is well known that *LFY* is required for the specification of floral organ identity, its precise role in organ outgrowth remains to be established and mainly a general role in auxin signalling and patterning has been proposed (Li et al., 2013; Parcy et al., 1998; Yamaguchi et al., 2014). To test whether LFY has a role in growth during early flower development, we live-imaged a number of flowers of the strong *lfy-12* loss of function line (Fig 10A, Fig S16, (Maizel and Weigel, 2004)) during the formation of the four sepal-like organs, i.e. comparable to the stage 2 to 4 transitions in the wild type. In all three acquisitions, the organs in the medio-lateral position grew out first, while the adaxial and abaxial primordia followed later and the boundary regions between sepals were less pronounced. The last sepal-like organ to grow out was slightly misplaced in one of the series, suggesting the beginning of a spiralled phyllotaxis. To analyse how these morphological phenotypes relate to growth in cells where *LFY* is normally expressed, the relative growth rates in the different domains where *LFY* was identified as a potential growth promoter were subsequently compared in one of the time-series for stage 3 and 4 (Fig. 10B, C; cf. Fig. 8, Figs. S13 and S14). In the mutant, these were defined based on morphology (e.g. negative curvature for the boundary) and lineage. The differences in growth rates between these zones were reduced in the mutant compared to wild type at the equivalent of stage 4 (Fig 10B, C) supporting that *LFY* is positively contributing to local growth in early flower development. This was true for all zones identified (Fig. 10C), and the cells in regions normally expressing *LFY* had consistently a higher reduction in growth rate compared to the regions where *LFY* is not expressed in wild type (Fig. 10D). This was partially due to the delay in outgrowth of sepal-like organs in the adaxial and abaxial positions in *lfy*. Whereas the difference between the boundary zone and sepals was most clearly reduced (30% reduction in median RGD), the difference between the meristem center and the sepals was less affected (14% reduction in median RGD).

**Fig 10.**
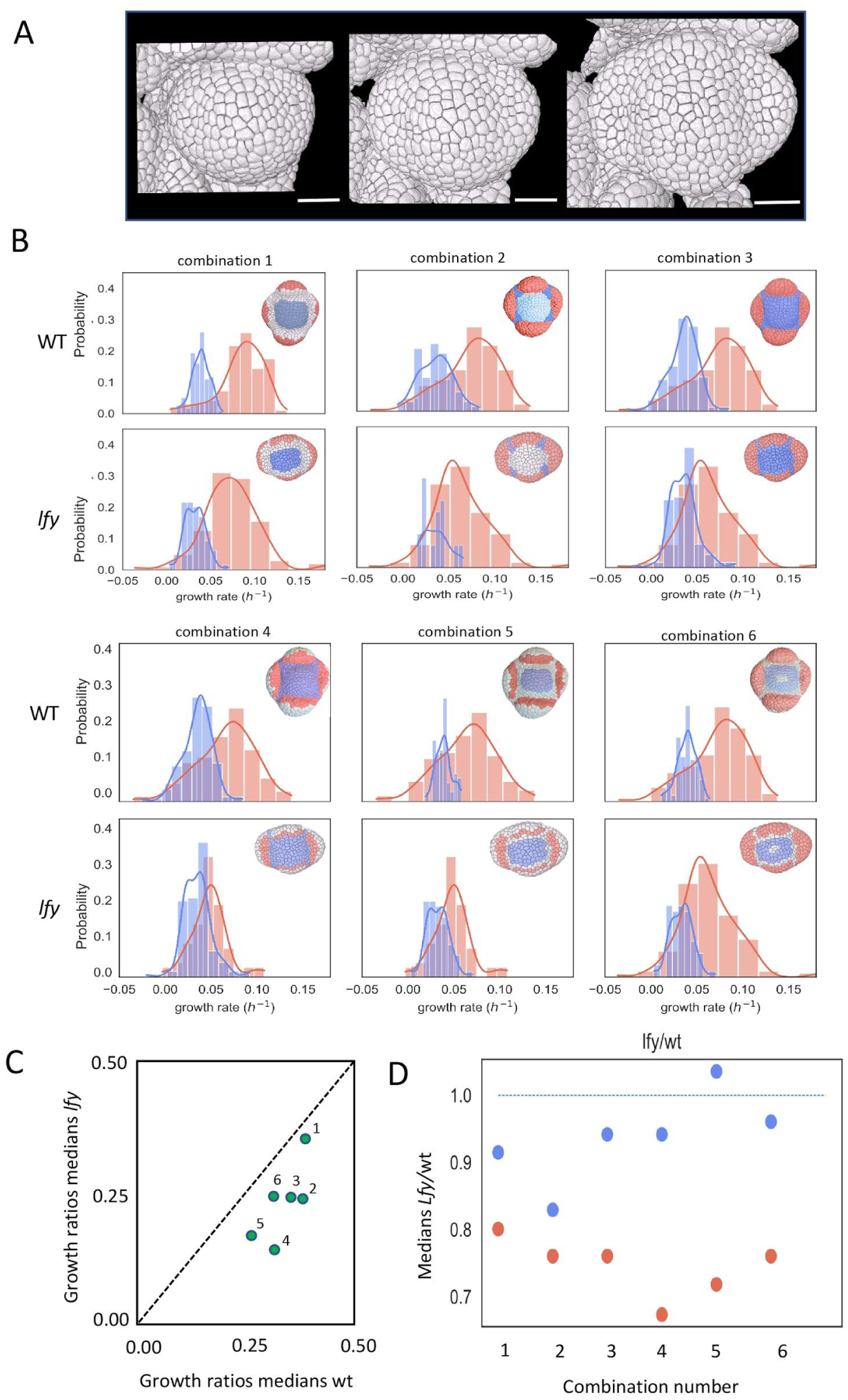
Comparison growth patterns in wildtype (wt) and *lfy* mutant (*lfy*). (**A**) part of the time series of the *lfy-12* mutant (see also Fig S16), equivalent to stage 2, 3 and 4 in WT. (**B**) Comparisons of growth rates between different equivalent regions in WT and *lfy* mutant at stage 4 (cf Fig 8 and S13-14). Blue and red histograms correspond to red and blue zones indicated in images. Combinations of regions are numbered 1-6. (**C**) Plot integrating relative growth differences (RGD: median blue graph - median red graph)/(median blue graph + median red graph) in WT and *lfy* mutant combinations 1-3 at stage 3 and 4. (**D**) Plot showing medians l*fy*/wt (blue median/blue median, red median/red median). In *lfy*, the medians of the blue and red histograms are systematically closer to each other than in wild type, hence the growth differential is reduced. Scale bars in A: 20 um.

## Discussion

We present here a detailed descriptive and quantitative model of early flower development. Integrating information at multiple scales, we have established a number of correlations which led us to propose an important set of testable hypotheses regarding the molecular regulatory network and its link to growth control. These hypotheses could not have been generated easily using other approaches.

### Molecular network dynamics

Based on an extensive analysis of the expression patterns of 28 genes, we propose the existence of at least 31 cell states in the L1 and L2 layers, which mounts up to 60 by taking into account ATML1, expressed in the epidermal layer.

Even considering a simple, binary on/off regulation of gene expression, the patterns are so complex that the analysis of spatial gene regulation becomes impossible based on visual inspection only. We therefore developed a set of tools to analyze the structural dynamics of the molecular network in space and its capacity to predict the observed expression patterns. This indicated that the published interaction network is not complete. Importantly, the addition of a limited set of single regulatory hypotheses not identified during our literature search significantly improved the predictive power of the network. With these extra hypotheses, the coarse network structure and composition are in principle sufficient to explain the observed expression patterns. As a result the proposed network can be a starting point for mechanistic gene regulatory models describing the developing patterns during early flower development. A complete set of testable hypotheses ranked by their effect on the predictive power of the model is given in tables S3 and S4 and we will only discuss here a limited number of striking examples.

As an example, the predictions concerning *AHP6* illustrate well the incomplete nature of the available data. *AHP6* has been described as a direct target of *MP* (Besnard et al., 2014b). Whereas the data summarized in the atlas are compatible with the hypothesis that *MP* is required, the latter has a much broader expression pattern than *AHP6*, suggesting further regulation. The simple hypothesis that *STM* could act as an inhibitor of expression would substantially improve the predicted pattern. This is in contrast to the conclusion by Besnard et al (Besnard et al., 2014a) who observed a temporary co-expression of both genes and concluded that AHP6 was probably not repressed by STM. However, their observations are also compatible with the hypothesis that *STM* inhibits *AHP6* above a certain threshold. In that case, the temporary overlap would correspond to the transition of cell state 7 to 8 or 9 to 16, when STM expression diminishes.

Novel hypotheses also resolve potential contradictions in the regulatory network model. The evidence so far suggests that the maximum levels of *LFY* and *CUC*, respectively involved in organ formation and the establishment of organ boundaries do not overlap (supplemental information). There are indications, however, that *LFY* directly activates *CUC2* (Yamaguchi et al., 2014). Although this seems contradictory, several experiments indicate that during early stage 3, *LFY* is expressed in the future boundary region, but at a lower level than elsewhere in the developing flower (supplementary data file 1 and 2). Likewise, *CUC2* shows a broader expression pattern than just the boundary region, again at a weaker level. In parallel, the expression pattern of *ANT*, an upstream regulator of *LFY*, is much more complementary to *CUC*, and our hypothesis that *ANT* is a negative regulator of *CUC* would substantially enhance the coherence between the predicted and observed patterns (Fig. 6A). This then would lead us to propose the existence of an incoherent feedforward motif between *ANT, LFY, CUC*, where *ANT* positively regulates *CUC* via *LFY* together with the proposed negative direct regulation. This type of motif is common in biological regulation and can tune the level and timing of expression of the individual genes (Goentoro et al., 2009); (Gruel et al., 2016).

### Gene activity and growth control

A number of genes have been explicitly associated with growth control, in particular during organ outgrowth (Nole-Wilson et al., 2005; Yamaguchi et al., 2016); (Besnard et al., 2014a)) or in the slowly growing central zone (Schoof et al., 2000). When looking at the distribution of growth rates for each gene, this correlation was confirmed only for some cases: *AHP6* is always expressed in rapidly growing cells during early organ formation, whereas the *CUC, PUCHI* and *CLV3* genes (Chandler and Werr, 2017; Hibara et al., 2006; Lenhard and Laux, 2003) are active in the slowly growing domains of the flower. However, it was not possible to make such direct correlations for other genes supposed to control morphogenesis (Fig. 7A).

This is well illustrated by the auxin-regulated transcription factor MP and its downstream targets *ANT* and *LFY* have been implicated together with *AIL6* in flower morphogenesis (Elliott et al., 1996; Krizek, 2009; Nole-Wilson and Krizek, 2006; Nole-Wilson et al., 2005; Yamaguchi et al., 2013). There is convincing evidence that ANT is involved in the control of cell proliferation (Mizukami and Fischer, 2000; Nole-Wilson et al., 2005). It was therefore surprising that *ANT* expression patterns did at first sight not correlate with particular growth rates: they are expressed in both slowly and rapidly growing cells throughout development. We therefore carried out a more detailed analysis taking into account co-expression in the different cell states, which finally clearly indicated that *ANT* could indeed be considered as potentially growth-promoting (Fig. 8B). The precise function of *LFY* and *MP* in growth control is less well established. *MP* loss-of-function leads to reduced flower outgrowth suggesting it promotes growth, but we found that it was mainly expressed in more slowly growing subpopulations. This points at a broader role of *MP* and is in line with some observations that over-expression of *MP* can also lead to reduced organ growth (Hardtke and Berleth, 1998). *LFY* is one of the main regulators of floral organ initiation and identity. It has been mainly associated with patterning events, and might also be involved in promoting auxin signalling (Li et al., 2013; Parcy et al., 1998; Yamaguchi et al., 2014). Although the distribution of growth rates in the LFY domains was again very broad (Fig. 7A), the combinatorial analysis indicated one of the strongest correlations with rapid growth of all genes tested (Fig. 8B). We therefore quantitatively analysed the growth patterns in the strong *lfy-12* loss of function mutant at a developmental stage equivalent to stage 3 and 4 in wild type plants. At the equivalent of stage 3 in the mutant, all four sepal-like organ primordia are marked by a local increase in auxin signalling, as revealed by the auxin inducible promoter DR5 (Yamaguchi et al., 2014), but we found that in particular organs in the abaxial and adaxial positions grow out later than the medio-lateral primordia, in contrast to the wild type. In addition, the differences in growth rates between the boundary and the adjacent zones is reduced, mainly when comparing stage 4 wild type flowers with an equivalent stage in *lfy-12*. The quantitative analysis, therefore, further supports the hypothesis that *LFY* is involved in coordinating cell expansion rates within specific subdomains, in particular to maintain sufficient growth rate differences between the sepals and sepal boundaries. The relative weak modifications in growth patterns might at first sight not seem significant. However, In terms of volume doubling times, the sepals in the wild type would achieve volume doubling two times faster than the boundary cells. In *lfy-12* this would be reduced to 1,6 times faster. Since growth is exponential and doubling times are in the order of 24h, these changes have the potential to induce substantial changes in organ shape over a few days.

## Conclusion

The integrated analysis of early flower development presented here suggests a number of further steps forward. The dataset can be easily extended by adding new expression patterns or expression gradients. By providing a dynamic template, it should be very useful in interpreting single cell sequencing efforts, which are precisely missing this type of detailed spatial information. The online atlas can also be enriched by information of different nature, including for instance cell polarity or mechanical properties. Other templates, coming from different time-series of wild type and mutants, can be added for further quantitative comparison as we show here for *lfy*. The ultimate aim would be to produce an artificial flower template using average behavior, or a collection of templates providing information on the variability existing in flower development. The atlas also provides an essential step towards the development of complex mechanistic models. Using the multiscale data of the atlas as input, it now becomes possible to compare quantitatively the results obtained *in vivo* with those coming from simulation where a large range of hypotheses can be tested in parallel, as we showed with boolean models here. Importantly, the interactive atlas is available online and provides a tool that can be used and further developed by the entire scientific community.

## Methods

### Live imaging

For clearly visualising cell edges for subsequent segmentation, we either used *Arabidopsis thaliana* (Col-0) plants containing a modified Yellow Fluorescent Protein (YFP) that is acylated and localised to the cell membrane (Willis et al., 2016) or staining with FM4-64. *lfy-12* (Columbia background, (Maizel and Weigel, 2004)) mutants were imaged after staining with FM4-64. Plants, grown in short days under standard conditions, were removed from soil soon after the transition to reproductive growth when the length of the inflorescence stem was less than 1 cm. These small plantlets, including roots, were carefully transferred into a plastic box containing molten, cooled 1% w/v agar supplemented with 2.2 grams l^−1^ MS salts and Gamborg B5 vitamins. Meristems were then dissected to remove obstructing flowers, the box filled with water and then imaged using either a Zeiss LSM780 or LSM700 upright confocal microscope with a 20x or 40x water dipping objective. Confocal Z-stacks were taken of primordia and detector pixel format, slice interval and zoom were set so that each resulting voxel is less than 300 nm^3^, as specified in the data set provided online (doi: provided on acceptance). Plants were transferred to the growth chamber between successive acquisitions.

### Segmentation and lineage tracking

Z-stacks of 2D (x-y) optical sections of five primordia expressing YFP were collected (numbered as FM1-5). A sixth meristem (FM6) was imaged after staining with FM4-64.

During confocal imaging of primordia, the flowers moved upwards due to meristem growth and stem elongation which led to oversampling of confocal optical sections in z-direction and an artificially stretched primordium after 3D reconstruction (Fig S11). To correct this artefact, we first manually selected cells on the epidermal layer whose thickness were least affected by the movement in z-direction (whose anticlinal wall normal were pointing close to the x-y plane direction). We then computed the average cell thickness of the selected cells (*L*1_*thickness*_). We then selected the epidermal cells whose thickness were the most affected by the movement (whose anticlinal wall normal were pointing close to the z-axis) and computed their thickness (*L*1*s*_*thickness*_). To correct the artefact, we divided the voxel thickness by (*L*1*s*_*thickness*_)/ *L*1_*thickness*_, see Figure S11. On a few time points, we also performed a rapid scan, where only two sections were made, to determine the height (in the Z-direction) of the flower bud as growth was negligible then (see also: (Willis et al., 2016)). These new estimations were close to the values calculated previously (<5%).

To quantify growth, we used the high-throughput 4D (space + time) image segmentation and tracking pipeline that we previously developed (Willis et al., 2016). This 4D imaging pipeline allows precise quantification of cellular growth over multiple cellular generations using MARS-ALT (Fernandez et al., 2010). Using the collection of image stacks, we used a three-dimensional auto-seeded watershed algorithm (Fernandez et al., 2010) to segment the cells. The segmented images were manually checked for segmentation errors (over-segmentation, under-segmentation, missing cell, or shape error). For this purpose, we conducted a visual inspection of segmentation quality of two dimensional (x-y) optical sections by comparing the optical section obtained using a confocal microscope and corresponding segmentation. In case of segmentation error, the contours of the cells were corrected on 2D sections. Cell volumes were calculated by voxel counting and multiplying this count by the voxel volume.

To track mother-daughter cell lineages, we first performed an affine transformation followed by a nonlinear registration using a block-matching framework (Commowick et al., 2008; Malandain and Michelin, 2017; Michelin et al., 2016) between two successive confocal acquisitions which computed the deformation field between them. Using this deformation field, we used ALT (Automatic Lineage Tracking) (Fernandez et al., 2010) to compute cell lineages, between consecutive segmented time points. The mother-daughter pairings were further inspected for errors and manually corrected and validated for L1 and L2 layers, see Figs. 2 and S1.

### Comparison Floral Meristems

We used the surface of flower primordia, represented by a point cloud, as the overall shape to compare the six sets of acquisitions. Although they go through similar developmental stages, each primordium has different cell arrangements. Also, there is no obvious way to synchronise flower development and hence the geometrical shape of individual time points of one series do not exactly correspond to the time points of other series. Therefore, we first computed the spatial and temporal correspondence between the six time series by quantifying shape differences (Michelin et al., 2016). The method uses a rigid transformation based on the hypothesis that two primordia at the same developmental stage have similar size and global shape. Since the shape of the flower primordium does not change sufficiently during stage 1 and 2 for such a comparison, we examined the overall shape changes during stage 3 and 4, when the sepals grow out and more dramatic changes in geometry are observed. To facilitate comparison, the temporal resolution of each time course was first refined to one hour using a dedicated 3D image interpolation method (Malandain and Michelin, 2017).

### Integration of the gene atlas into Morphonet and AtlasViewer

For online display and the introduction of expression patterns, Morphonet was used (http://www.morphonet.org). For access see supplemental information. Morphonet is a web-based interactive platform for visualization and sharing of complex morphological data and metadata (Leggio et al., 2019). Exploiting its Unity (https://unity.com) 3D visual engine, it offers a vast assortment of possible interactions with 2, 3 and 4D datasets. Through a flexible hierarchical representation of biological structures and dedicated formats for associated metadata, users can follow the dynamics of biological shapes, onto which associated quantitative and qualitative properties can be projected.

Cells are represented with meshes in Morphonet, and the meshes generated from the cell segmentation were converted to *obj* format and uploaded to Morphonet together with lineage information.

For the introduction of gene expression patterns, five time points were chosen (Fig. 3, (Smyth et al., 1990) corresponding to:

- the initium stage,
- stage 1 (the flower starts to bulge out),
- stage 2 (a globular bud is formed, separated from the inflorescence meristem),
- stage 3 (the ab- and adaxial sepals start to grow out)
- stage 4 (all sepals are clearly growing out and the four whorls have been specified).

The expression patterns of 28 genes were subsequently introduced. This was in short done as follows (see also Supplementary Information):

1. <u>Collection of data from the literature.</u> As many available image sets as possible were collected from the literature. This included GFP expression patterns and RNA in situ hybridizations.
2. <u>Complete existing data with new data.</u> We completed these data by our own RNA in situ hybridizations. For this purpose, we generated a further 60 sets of serial sections with the expression patterns of 20 genes (Original data available online (doi: provided upon acceptance), Supplementary Information).
3. <u>Manual annotation of timepoints.</u> To facilitate interpretation, we used a binary notation (i.e. genes are either on or off). Since often only 2D data in the form of sections were available and published 3D GFP data were usually partial, it was not possible to project directly the patterns automatically on the atlas. Instead, we used a manual protocol using the annotation tool in Morphonet by clicking on the individual cells. Cells potentially expressing a particular gene were identified by manually projecting sections of in situ hybridizations or confocal sections on the different time points of the atlas. Whenever possible, cell numbers were counted to estimate the size of the expression pattern. For each gene 2-4 datasets available in the literature were identified. We encountered three different cases:

i. The patterns of the in situ hybridization and/or GFP were simple to interpret, and zones of expression could be unambiguously identified (Fig S3). In the absence of GFP patterns, the use of serial sections was crucial.
ii. There was a conflict between results obtained using GFP expression and in situ hybridizations. In that case, the in situ hybridization results were used (Fig S4).
iii. If information on both protein levels and RNA levels were available, the RNA pattern or promoter activity (in case of GFP construct) was retained (Fig S5).
4. The obtained patterns were subsequently refined using information of co-expression (Fig. S5) or based on information on mutual regulation (e.g. AG and AP1 mutually inhibit each other).

The references and images for each gene are summarized in supplementary file 2.

In addition, we integrated the four-dimensional gene expression data together with all segmented and tracked time courses in an open source standalone software platform, called ***AtlasViewer*** (available at https://gitlab.com/slcu/teamHJ/publications/refahi_etal_2020/-/tree/master/atlasviewer/atlasviewer). The patterns were imported directly from Morphonet into AtlasViewer. Visualization of combination of gene expressions using AtlasViewer facilitated the identification and correction of annotation errors.

### Cell states, clustering and transition graphs

After assigning expression values for the 27 genes to individual cells, this boolean element vector is used to define a *cell state* for each cell (Fig. 4A-B). Similarities between cell states were calculated using the Hamming distance, i.e. the sum of the absolute differences between the vector elements (Fig. 4C). The states were clustered using hierarchical agglomerative clustering from SciPy package using Ward’s method, which uses Ward variance minimization algorithm, leading to a dendrogram. Manual flipping at the dendrogram nodes and identification of tissue structures were applied to generate the final graph (Fig. 3D).

To illustrate the evolution of cell states over time, cell lineages were used to generate a *pattern transition graph* whose vertices were cell states. An arrow connected two vertices if and only if any of the descendant cells of the source cell state acquired the target cell state at the next developmental stage. We then assigned weights to arrows as the number of descendant cells in a specific state divided by the total number of descendant cells. More precisely, let *x → y* be an arc of the transition graph, where *x* and *y* are cell states. The assigned weight, *w*, is defined as *w* = (#descendant cells of cells in pattern *x* in pattern *y*) / (#descendant cells of cells in pattern *x*), where # denotes number of cells. To extract the main structure of the pattern transition graph, we then pruned by keeping the arrows whose weight were equal or greater than a manually defined threshold of 0.2. However, in this representation the descendent patterns with a small number of cells are penalized. We therefore also computed a *reverse pattern transition graph* where arcs pointed from *descendent patterns to their ancestors*. For each arrow, *x← y*, in the reverse transition graph, where *x* and *y* were cells states, we assigned a weight *w’*, defined as, *w’=* (#cells in pattern *y* whose ancestors are in pattern *x*) / (#cells of cells in pattern *y*). The weights were then used to prune the reverse transition graph by removing the arrows whose weight were below 0.2. The pruned transition graphs were then merged into a single transition graph, Figure 4.

### Cell growth and anisotropy rates and correlation analysis with gene expression and states

Assuming exponential growth of a cell of volume 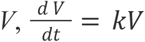, with a constant relative growth rate we compute *k* from cell volumes and lineage information for a cell *c* at time point *t*_*i*_, 0 ≤ *i* ≤ 4 (index corresponding to the 5 developmental stages considered), as:

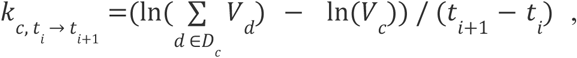

where V_*c*_ is the volume of the mother cell *c*, V_*d*_ is the volume of a daughter cell *d*, and *D*_*c*_ is the set of daughter cells of *c* at time point *t*_*i*+1_.

Projection of computed values on time courses can be done either on the mother cell or on the daughter cells (e.g. Fig. 5C). Correlation analysis between the computed values and gene expression was done for cells expressing a specific gene in both forward (following all cells expressing a gene at one time point to the next) and backward (tracking all cells expressing a gene at a specific time backward in time). Similar correlation analysis is done for averaging cells within a specific cell state. When both forward and backward growth are available, the growth rate of a cell *c*at time point *t*_*i*_ was computed as the average of the two to quantify growth at a specific time (e.g. Fig. 7C):

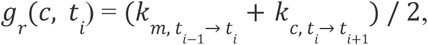

where *m*is the mother cell of *c*. In the cases where only one was available, the reported values are for the first time point the forward growth calculation and for the last time point the backward growth calculation.

3D growth anisotropy was computed using cell and lineage information by identifying and matching key points on the cell and set of daughter cells’ surface. We define key points as the smallest set of points on the cell surface enveloping the cell (Convex Hull, calculated using Python bindings to QHull library, http://www.qhull.org/). The Convex Hulls of each cell and its daughter cells were centred and then mapped according to a nearest neighbour criterion. We then calculated the best (least-squares estimation) linear transformation *A* between the two hulls:

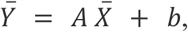

Where 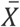 is the collection of the selected points for a cell and 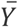 the collection of corresponding points on the daughter cells’ surface.

The transformation was decomposed into three transformations, two rotations and an expansion using the Singular Value Decomposition (SVD):

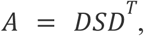

where *D*and *D*^*T*^ are the rotations and *S*is the expansion. The singular values of *S*(*s*_1_, *s*_2_, *s*_3_) represent the length of the expansion along the axes. The anisotropy is then computed using these expansion lengths as the fractional anisotropy *a*:

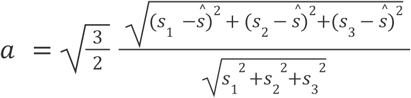

where ŝ = (*s*_1_ + *s*_2_ + *s*_3_)/3. The fractional anisotropy is a scalar value ranging from 0 (expansion was equal in all directions) to 1 (expansion was in only one direction). The anisotropy rate is the fractional anisotropy per hour.

Correlation analysis between growth anisotropy and gene expression and states, like in the case of growth rates, can be done in both forward and backward directions. When both are available the anisotropy rate at time point *t*_*i*_, was calculated as the average of the forward anisotropy rate calculation from *t*_*i*_ to *t*_*i*+1_ and the backward growth anisotropy rate calculation from *t*_*i*−1_ to *t*_*i*_.

### Regulatory network analysis

We extracted the gene regulatory interactions, inhibition and activation, from the published literature based on the available data on direct binding on promoter regions and mutant analyses (Fig. 1).

### Boolean regulatory terms

To analyse the regulatory network, we defined a subset of boolean rules combining the input arrows in the regulation network (i.e. repression and activation), and examined whether they could reproduce the spatial gene expression patterns of the Atlas. Since the input arrows indicate either positive (activation) or negative (repression) regulation, the corresponding boolean regulatory terms that we define have two sub-terms for activation and repression. We next define the types of terms used for the full regulatory term and the repression and activation sub-terms (for further details on network construction see (Moignard et al., 2015)).

Regarding notation we assume we have a set of names where *g*_1_, *g*_2_, … ∈ *Symb* range over symbols representing gene names, ϵ, *t, t* ′ ∈ *TExp* range over (possibly empty) activation/repression terms, and *r, r*′ ∈ *RExpr* over full regulatory terms. We will give the following definitions using the ‘is defined as’ symbol (∷ =) to separate the class being defined (e.g. regulatory expressions) with its definitions and the ‘alternative’ symbol (|) to separate definitions in cases where there is more than one.

The class of regulatory interactions for each gene is defined as

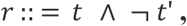

where *r* ∈ *RExpr* is a full regulatory term for a gene including an activation term (*t* ∈ *TExp*) and a repression term (*t*′ ∈ *TExp*) combined by a *logical and* (∧) and *logical not* (¬).

Each term can be empty (ϵ), a gene name (*g* ∈ *Symb*syntactic case), or a combination of gene names with conjunction or disjunction, as described by

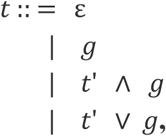

where ∨ represents a *logical or*.

We next define an evaluation function to evaluate the activation and repression terms to values in the set *B* = {*T, F*}(*T* for true, and *F*for false). Evaluation happens in a value environment providing values (True or False) for the gene names in the terms. To represent this context we use a sequence of gene name value bindings σ = *g*_1_ : *b*_1_ …, *g*_*n*_ : *b*_*n*_ where *g*_1_ …, *g*_*n*_ ∈ *Symb*(gene names) and *b*_1_, …, *b*_*n*_ ∈ *B*(True or False representing expression or non-expression of the corresponding gene), where the gene names are required to be distinct. We will sometimes treat this context as a function with finite domain; for example to obtain the value of *g*_1_ in σwe wrote σ(*g*_1_). The evaluation function [| |] : *TExp* → *B* in a value environment σis then (per syntactic case):

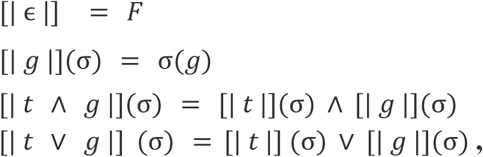

where *t* ∈ *TExp, g* ∈ *Symb*. For both *TExp*’s and *RExpr*’s evaluation functions, we assume that the given environments are well-formed, i.e. they contain mappings for all the gene names that appear in the expression being evaluated. Note that the above evaluation implies left association of expressions so, for example, the expression *a* ∧ *b* ∨ *c* is evaluated as (*a* ∧ *b*) ∨ *c*.

The evaluation function for the full regulatory term [| |] : *RExpr* → *B*is:

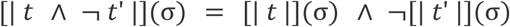

Two *RExpr*’s are *semantically-equivalent* if they contain the same gene names and given the same environment they evaluate to the same value.

### Translating the gene regulatory network into boolean regulatory terms

Given a gene regulatory network, as in Fig. 1, for each gene we have a set of activators (positive regulation) and a set of repressors (negative regulation), but there is no information on how their inputs combine to control expression of their target gene. In order to see the most likely boolean regulatory term between the regulations, we enumerated all the possible *TExpr*’s for the activators, all the possible *TExpr*’s for the repressors, combined them into regulatory terms (*RExpr*’s) and scored them based on how well they agree with the expression data (Supplemental Table 2).

Given a set of input activators/repressors, {*g*_1_, …, *g*_*n*_} for a gene there are *n*! · 2^n−1^ possible *TExp*’s. For each permutation of the input genes (out of *n*! possible), we have a choice of disjunction or conjunction between them. For example for 2 activator genes {*g*_1_, *g*_2_}we can generate the following terms: {*g*_1_ ∧ *g*_2_, *g*_1_ ∨ *g*_2_, *g*_2_ ∧ *g*_1_, *g*_2_ ∨ *g*_1_}. If we also had 2 repressors then the number of terms becomes 16. While the number grows very quickly with *n*, we found that is not prohibitive for the number of genes we have here (*n* < 5).

Each cell in a tissue dataset implies a value environment; for example if in a cell gene *g*_1_ is on, gene *g*_2_ is off, gene *g*_3_ is off, and *g*_4_ is off we get a value environment *g*_1_ : *T, g*_2_ : *F, g*_3_ : *F, g*_4_ : *F*. We can then evaluate the generated boolean regulatory terms in this environment so for example for an expression for *g*_4_ =(*g*_1_ ∧ *g*_2_) ∧ ¬*g*_1_ we can evaluate for that cell [| (*g*_1_ ∧ *g*_2_) ∧ ¬*g*_1_ |](*g*_1_ : *T, g*_2_ : *F, g*_3_ : *F, g*_4_ : *F*) = *F*, which (for this example cell) agrees with the actual value of *g*_4_.

### Hypotheses generation

In order to generate new hypotheses for each gene, we enumerated all possible changes in the form of single regulatory interactions between genes of the published gene regulatory network. Therefore either existing single interactions were removed or new ones were added. For each modified interaction, we generated all possible *TExp*’s and full regulatory terms as before.

Suppose we have a universe of genes *G* = {*g*_1_, *g*_2_, *g*_3_, *g*_4_}. Then, for a gene A with two activators (*g*_1_, *g*_2_)and one repressor *g*_4_, the following set of regulatory interactions for that gene would be generated: {(acts=(*g*_1_, *g*_2_, *g*_3_), reprs=(*g*_4_))), (acts=(*g*_1_, *g*_2_), reprs=(*g*_4_, *g*_3_)), (acts=(*g*_1_), reprs=(*g*_4_)), (acts=(*g*_2_), reprs=(*g*_4_)), (acts=(*g*_1_, *g*_2_), reprs=())}. Note that we only added inputs from genes that were not in the original network, whereas the gene *g*_3_, could represent any other gene in the network not yet connected to gene A. For each set of inputs (activators and repressors) we then generated and scored regulatory terms as before.

### Pattern Evaluation

To score an expression pattern generated from cell lineage, the regulatory network of a particular gene (Fig. 1), or a boolean model including hypothesis for an entire tissue, we evaluate it for all the cells and calculate the *Balanced Accuracy* (BAcc) for its predictions. BAcc is defined as 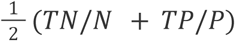 where *TN*is the number of true negatives (# of cells where the expression evaluates to false and the actual value of that gene is false --- like the example above), *TP*is the number of true positives (#of cells where the expression evaluates to true and the actual value of the gene is true), *P*is the number of positives (#of cells where the value of the gene is true), and *N*is the number of negatives (# of cells where the value of the gene is false). We chose BAcc as measure since it also penalizes errors where only few cells have an expression (or opposite). We also tried alternative similarity measures, such as Mutual Information and % correct. This led to the same conclusions.

The scores and therefore the best expressions are not necessarily the same for the tissues at different timepoints (Supplemental Table S3). For each gene and each time point we merged and ranked the generated regulatory expressions for all the proposed interactions keeping only the ones that are within 10% of the best expression for that time point. Starting from the last time point going backwards we then identify expressions that appear near the top in more than one time point for a single coherent hypothesis.

### Model comparison and random models

In order to establish a baseline for the comparison of *RExpr*’s generated by the hypotheses, the *RExpr*’s based on the reference network, and lineage prediction, we also constructed a *random network* with the same structure as the gene regulatory network in Fig. 1, where all the inputs are replaced with random inputs. For the results reported in the main text we generated 100 random *RExpr*’s per gene based on random inputs with the same numbers as they appear in the gene regulatory network(Fig. 1). These were then scored (using BAcc) per time point and averaged.

All the p-values reported in the main text are the result of a paired t-test between the scores of all the genes under the different models, e.g. lineage scores vs best hypothesis scores.

### Growth regulation

In order to examine growth regulation by gene expression we used Relative Growth Difference (RGD) to compare growth differences between populations of cells (defining regions on a tissue), defined as:

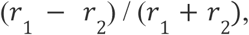

where *r*_1_ is the median growth rate of the first population of cells and *r*_2_ is the median growth rates of the second population of cells. The RGD ranges from 0 to 1 except in some cases early on in development where it can be > 1 when one of the regions considered has a negative median growth rate.

To examine the gene regulation of growth by the action of single genes, we examined the differences between the population of cells expressing a gene versus the population of cells not expressing the gene. Spatially this defines two regions, which can be described as boolean expressions for a gene, *g*, as *g*and ¬*g*. Writing [*e*]_*T*_ for the set of cells where expression *e*is True and 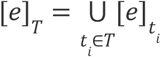 for the union of of all the cells where the expression is True over all time points, the RGD of a gene *g*was calculated as:

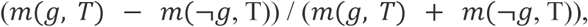

where 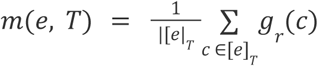 is the median growth rate of the cells in the regions defined by the expression, *e*, over a set of time points, *T*. The evaluation of an expression in a cell follows the procedure described in the ‘Boolean regulatory expressions’ section. The growth rate of a cell is, when possible, the average of backward and forward growth rates as described above.

In order to get a more fine-tuned understanding of growth gene regulation we extended our single-gene analysis to pairs of genes that are co-expressed. Each combination of co-expressed genes implicitly defines two regions (populations of cells) on the flower tissue at any time point. These two regions can be defined using boolean expressions as, *g*_1_ ∧ *g*_2_ (region where they are co-expressed) and *g*_1_ ∧ ¬ *g*_2_ (region where only one of them is expressed). For each pair of co-expressed genes the RGD at time point *t*_*i*_ (colour maps in the heatmaps in Supp. Figs 11, 12, 13) was calculated as:

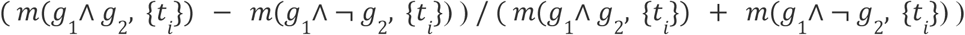

In order to get the most common regional separation implied by gene pairs at a time point we grouped the pairs into categories (numbered annotations in the heatmaps in Supp. Figs 11, 12, 13) defining the same regions. Two pairs of genes *g*_1_, *g*_2_ and *g*′_1_, *g*′_2_define the same regions at *t*if the two boolean expressions they imply select the same set of cells:

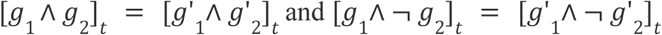

In Supp. Figs 11, 12 and 13 we only display the groupings in the top 50% (by RGD) of the pairs at that time point up to a maximum of 6 groupings. Groupings are also sorted by RGD so the group with index 1 has the highest RGD and so on.

## Supporting information

All supplemental information

## Data Availability

Confocal z-stacks, segmentation files, cell lineage information, and in situ hybridisation images are available online (doi: provided on acceptance). Software scripts for reproducing the analysis performed in this work is available via the Sainsbury Laboratory gitlab repository (https://gitlab.com/slcu/teamhj/publications/refahi_etal_2019). All data is interactively minable via the MorphoNet ATLAS (http://www.morphonet.org).

## Author contribution

Y.R., A.Z., H.J., J.T. designed research; Y.R., A.Z., R.W., G.M., J.L.,H.J., J.T., A.A., L.V., A-E.R., P.D. and F.Z. performed research; Y.R.,A.Z., H.J., J.T., R.W., G.M., J.L., N.P., F.B., C.G., E.M., G.Ma., contributed new reagents/analytic tools; Y.R., H.J., J.T., A.Z., G.M., J.L, G.Ma., A.A., L.V. and P.D. analyzed data; Y.R.,A.Z., H.J., J.T. wrote the paper.

## Acknowledgements

The authors would like to thank Arun Sampathkumar, OIivier Hamant and Marie Monniaux for discussion, Anuradha Kar for help with MorphoNet. Weibing Yang provided the *lfy* mutant and helped with imaging. JT, YR, AA, LV, A-ER and FZ were funded by the ERC grant MORPHOGENETICS. This work was also supported by ANR BIOMOD (ANR-19-CE43-0010) grant (to YR.) and Gene2Shape ERACAPS grant to JT, HJ and AZ. The E.M.M laboratory is funded by the Howard Hughes Medical Institute, and the work here was supported by the U.S. National Science Foundation Division of Integrative Organismal Systems grant IOS1826567.

HJ was funded by the Gatsby Charitable Foundation (GAT3395/PR4B) BBSRC (BB/S004645/1)

