## Supplementary material for "A multiscale analysis of early flower development in Arabidopsis provides an integrated view of molecular regulation and growth control": All supplemental information

**Supplemental Figures S1-S16**
**Supplemental Tables S1-S4**
**Litterature**
**Justification of gene expression**
**How to access the Atlas on Morponet**

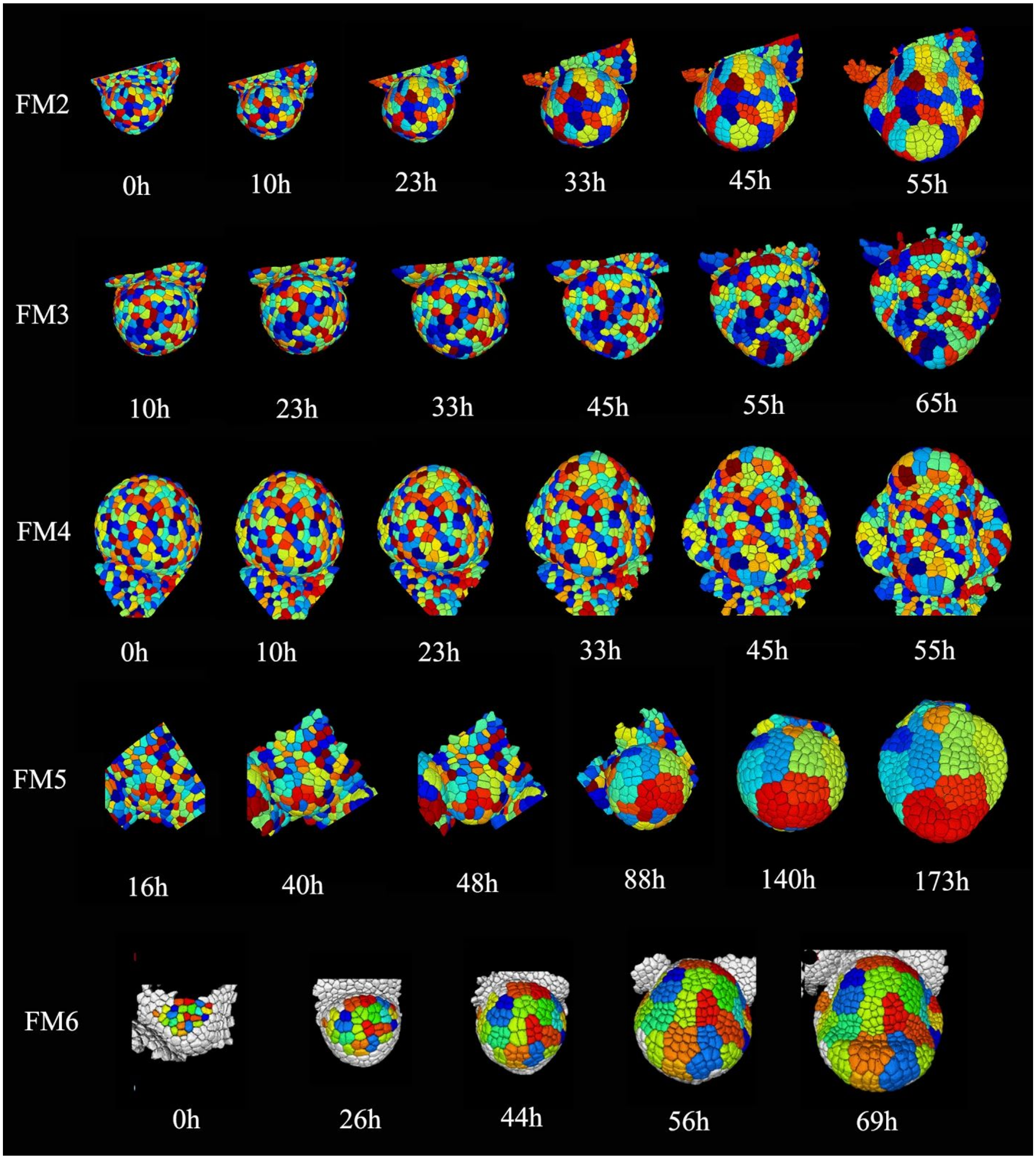

**Figure S1** – Surface rendering of segmented images of FM 2-6 time courses. The cells are colored according to computed lineages. Numbers indicate number of hours after first acquisition.

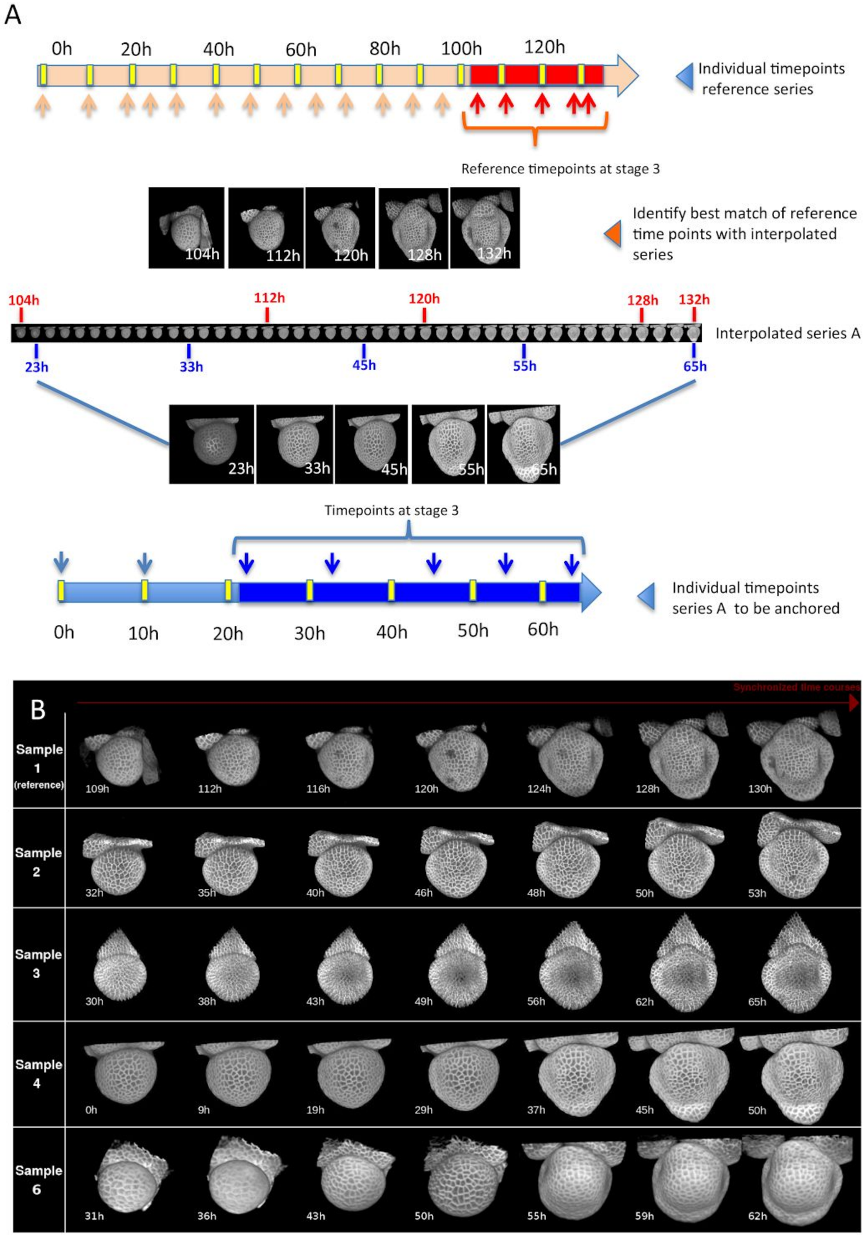

**Figure S2 A)** Comparison of reference series with 18 time points, from initium until late stage 3 with another series 'A' with 7 time points. Since the changes in shape are most striking during stage 3, this developmental window was chosen for the comparison (dark red zone in the reference series, dark blue in series A. First, a 3D image interpolation is performed to improve temporal resolution to1h intervals. The 5 timepoints of stage 3 of the reference series are then compared to this interpolated series. The two series are then 'anchored' via shapes that are closest to each other. Note that there is a difference in time scale, as some meristems grow more quickly than others.

**B)** Result of temporal alignments of confocal time series of floral meristems. The 3D images of the sample 1 (reference sequence) are paired to their best match in shape and size in the samples 2, 3, 4 and 6. This was not possible for series 5 (see text for details).

A

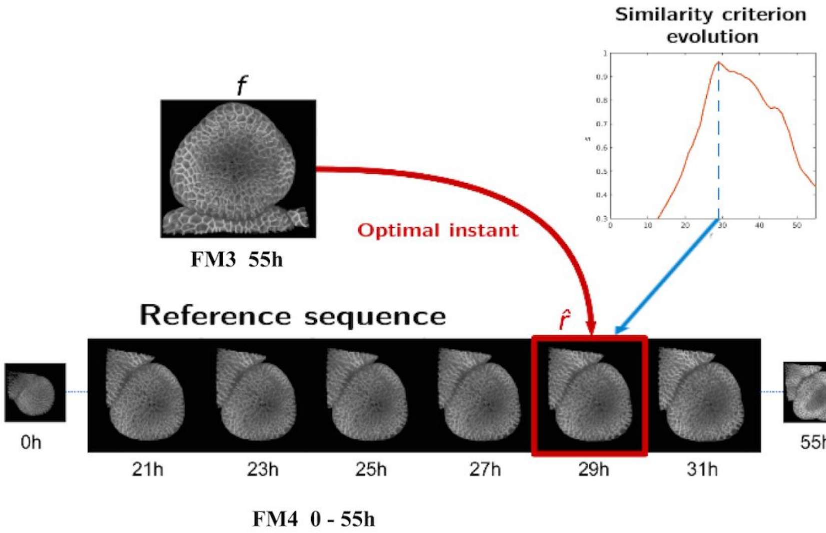

B

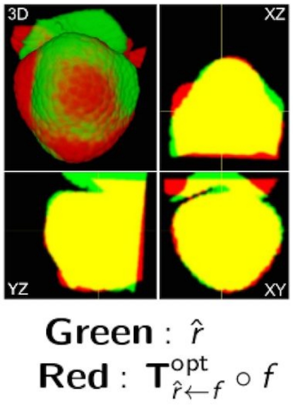

C

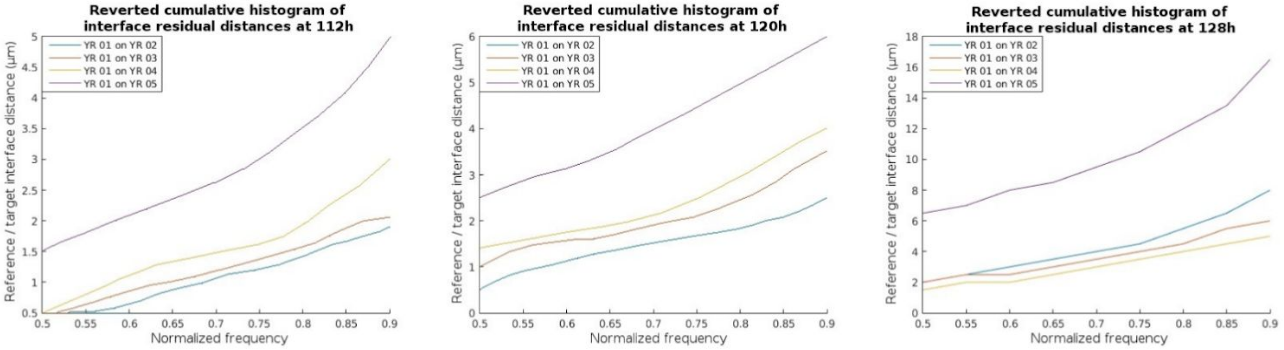

**Fig S3.** Examples of comparison of meristem shapes. **A)** Comparison between meristem 3 and 4. Time point at 55h is systematically compared with all timepoints of the interpolated series of meristem 3 and the residual distance between the two surfaces is used to calculate the similarity. In this example, the similarity is maximal at 29h. **B)** Optimal projection of the two timepoints which most closely resemble each other. **C**) Comparison of optimal alignment between different timepoints of the reference meristem (FM 1) and four other meristems (FM 2-5) at different timepoints. The cumulative distance between the point clouds of both surfaces is given as a normalized frequency (e.g. 4 um has a Normalized frequency of 0.75 at 128h : 75% of the points are at a distance of less than 4 um) . Note that the difference between meristem 5 and meristem 1 is much higher. As a result, this meristem could not be aligned and synchronized with the reference series.

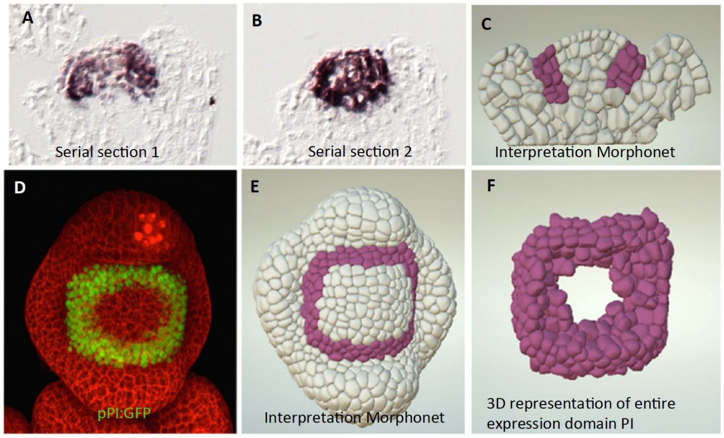

**Fig S4**. Interpretation and annotation of expression pattern (A, B, D) of PI (pPI:GFP and in situ hybridization) in Morphonet (C, E, F) at floral stage 3. This is a case where the patterns of the in situ hybridization (A,B) and (D) are simple to interpret, and zones of expression can be unambiguously identified. Note the importance of serial sections in interpreting the in situ pattern. Both types of markers indicate a ring-like expression pattern of 2-3 cells wide and 3-4 cells deep. (C) Shows a cross section of the pattern in Morphonet, (E) a top view. In (F) only cells expressing PI are represented, to show the overall shape of the expression pattern.

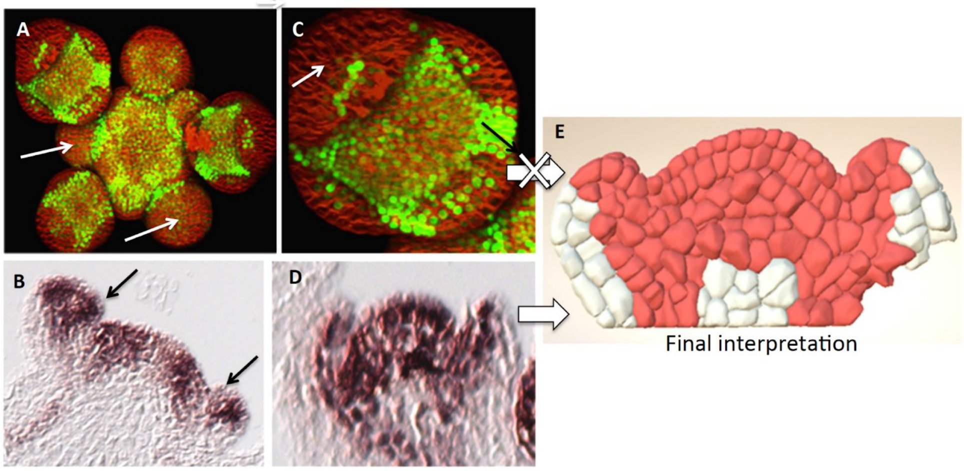

**Fig S5.** Expression pattern of MP. Conflicting information coming from a line expressing pMP: GFP-NS (A,C) and in situ hybridization (B, D). The GFP pattern suggests that there is no expression at early stages of flower development (arrows in A), whereas in situ hybridizations indicate there is strong expression (arrows in B). Likewise, the GFP pattern indicates that there is only expression at the tip of the outgrowing sepals and weaker expression in the floral meristem (C), whereas in in situ hybridisation indicates a strong expression at the adaxial side of the sepal and throughout the floral meristem. (E) in Morphonet, the in situ pattern was retained.

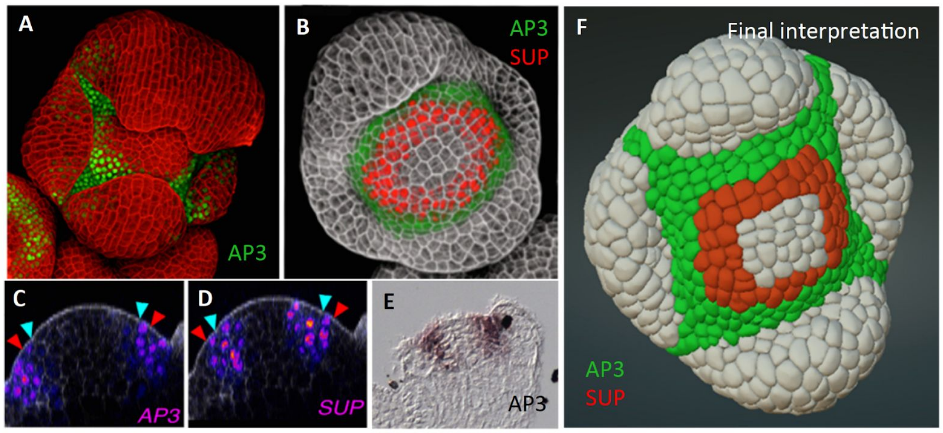

**Fig S6.** Promoter activity **(A)** in *pAP3:GFP* and protein levels **(B)** in *gAP3:GFP*. **(B-D)** double labelling of *pAP3:CFPN7* and gSUP:VenusN7. **(E)** In situ RNA pattern of AP3. AP3 is expressed at the periphery of the floral meristem and the promoter activity extends into the organ boundaries whereas the protein is absent from lateral boundaries between sepals. (compare A and B). *gSUP* is expressed in a ring inside the AP3 domain (B-D). There is a slight overlap (about one cell wide), but SUP expression is stronger in cells that do not express AP3, and vice versa (C,D). In the binary interpretation the patterns do not overlap (see (Prunet et al., 2017) for details on the lines used).

A

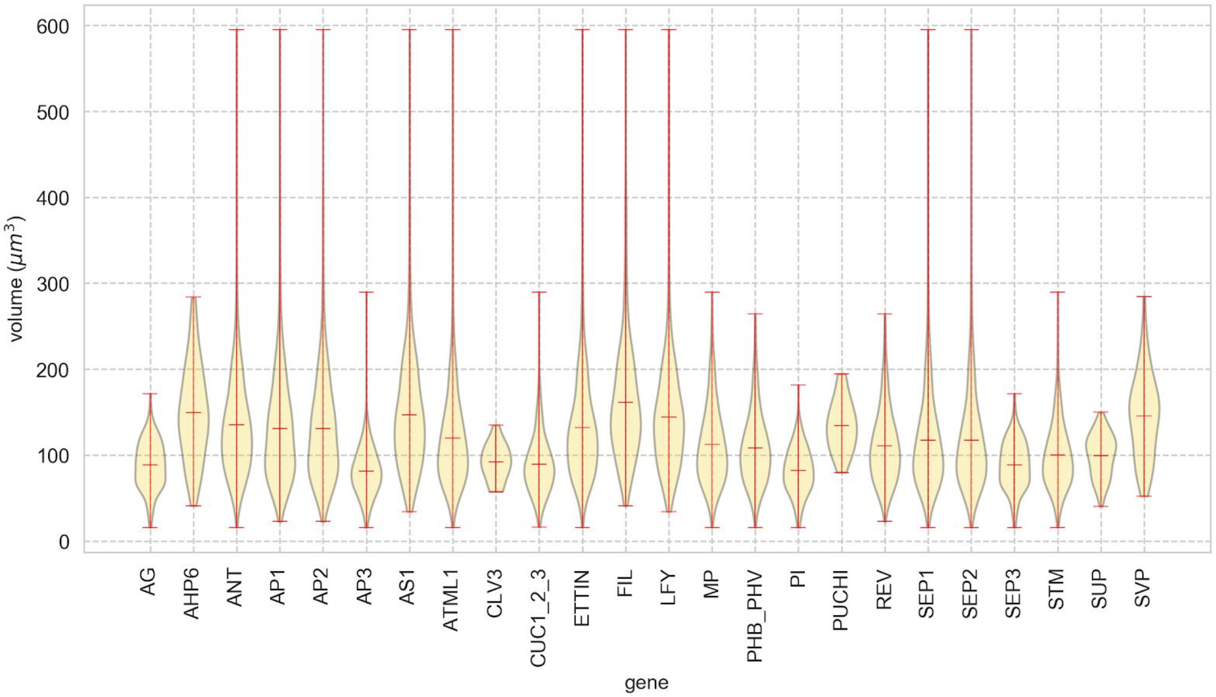

B

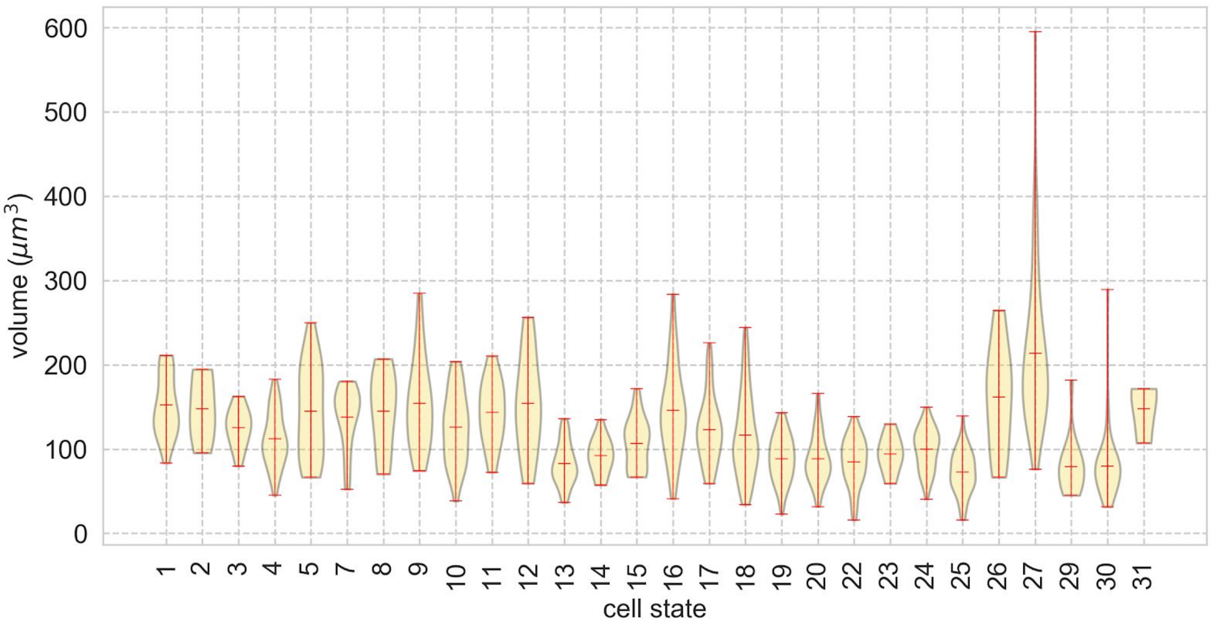

**Figure S7 A**) Distribution of volumes of cells expressing specific genes. **B)** Distribution of volumes of cells in particular states.

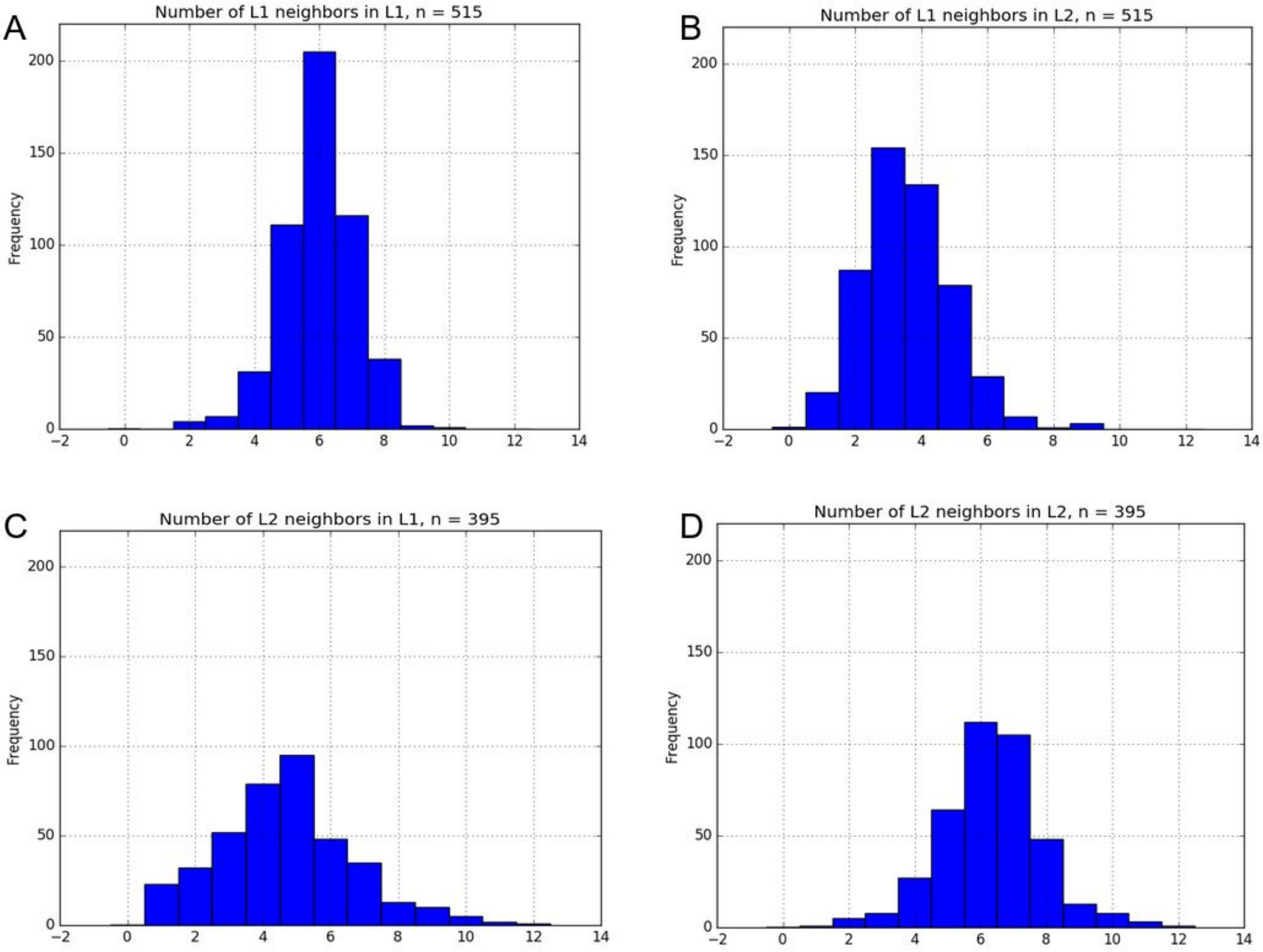

**Figure S8.** Neighbor number distributions of cells in L1 and L2 at 132h for FM1. (A) Distribution of the number of L1 neighbors surrounding a cell in L1. (B) Distribution of the number of L1 neighbors in L2. (C) Distribution of the number of L2 neighbors in L1. (D) Distribution of the number of L21 neighbors surrounding a cell in L2.

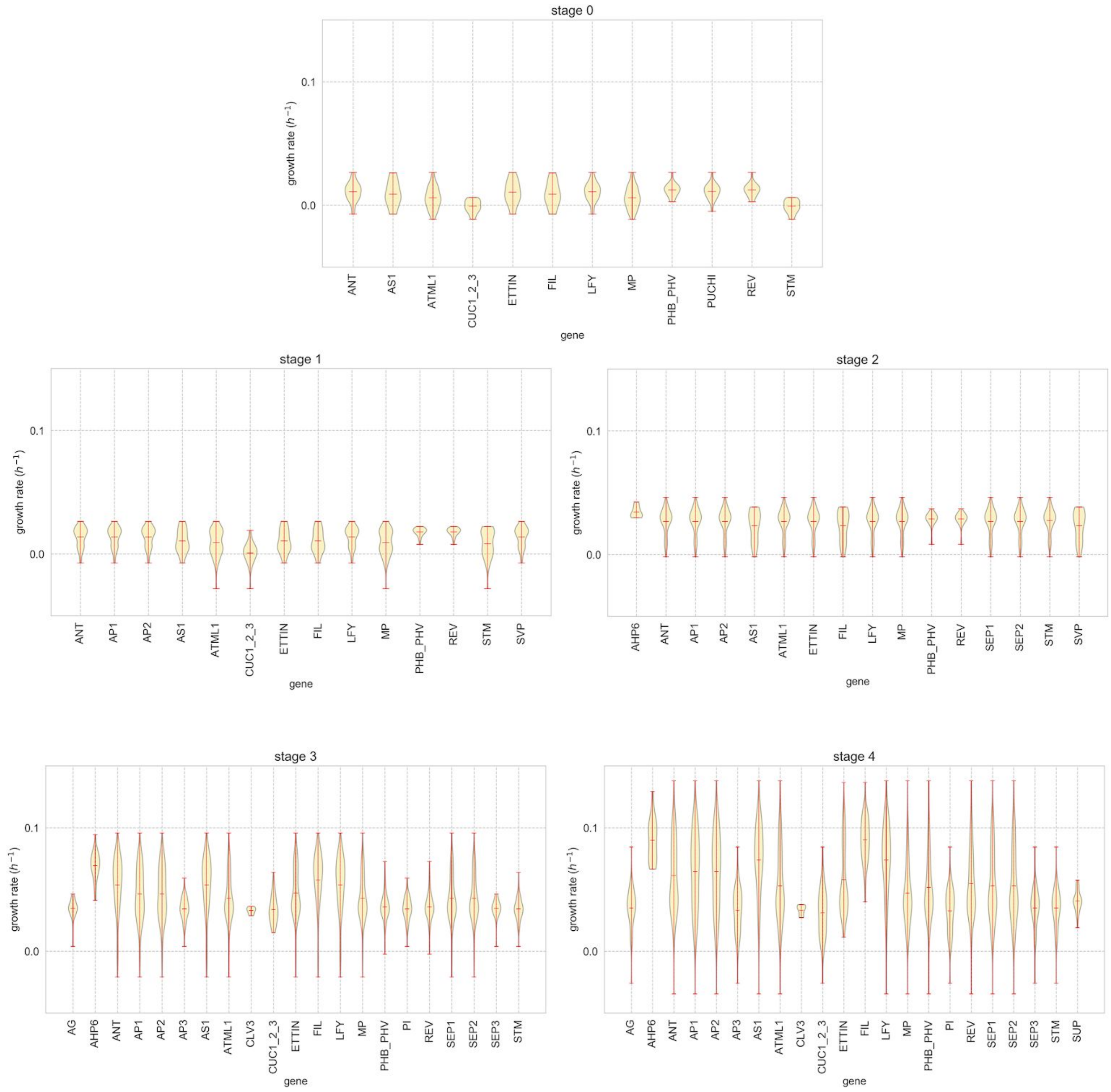

.

**Figure S9.** Growth rates/hour at different timepoints, for gene expression domains at different stages of development. Certain gene domains can have different behaviours, depending on the developmental stage. FIL, for example, is associated with more slowly growing cells at stage 1 and 2, and then amongst the most rapidly growing ones later on.

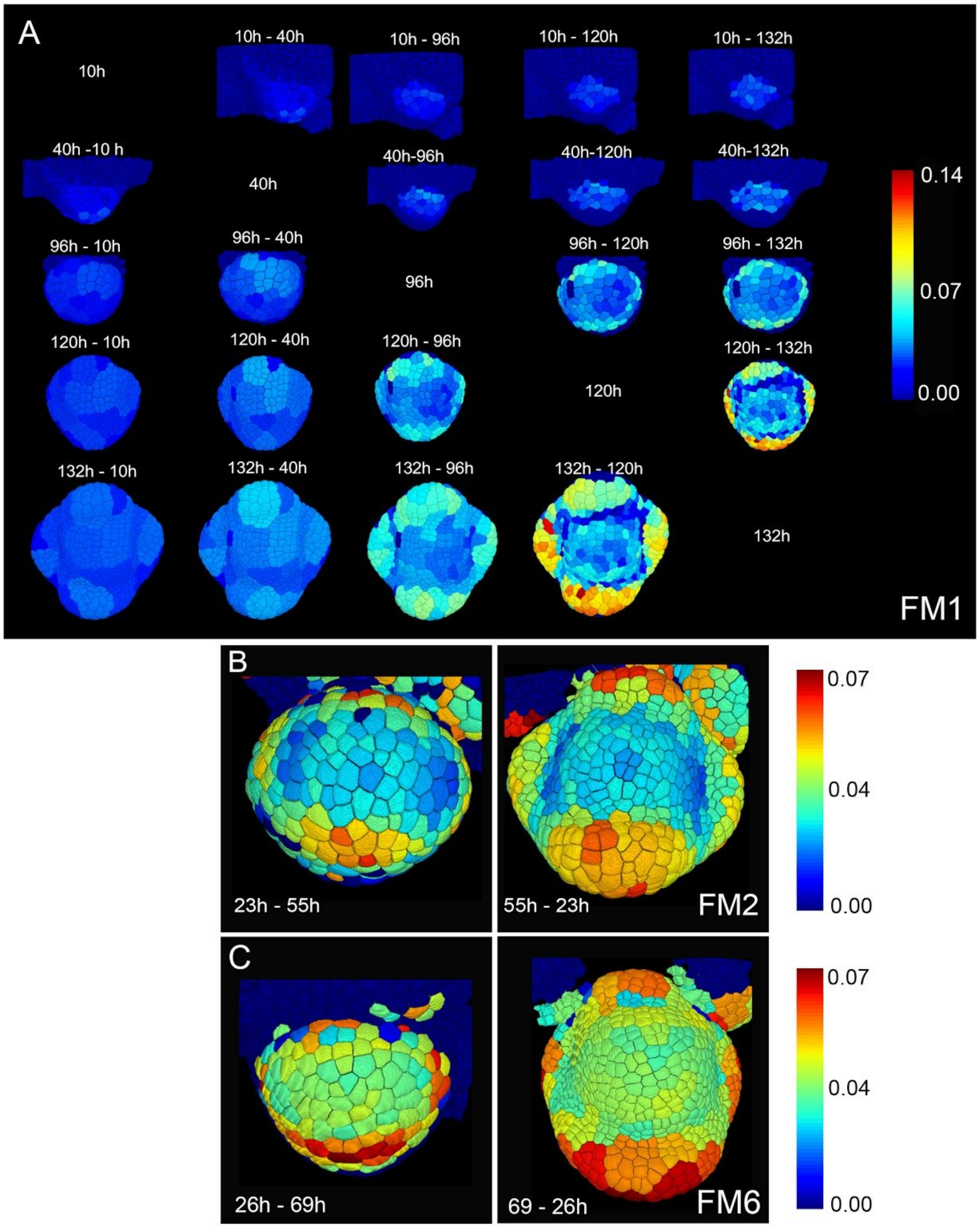

**Figure S10** (**A**) Backward (how much cells have grown, below diagonal) and forward (how much cells will grow, above diagonal) growth rates as um3 per hour between time points (Flower Meristem 1), illustrating that the increased growth rate in the sepal is determined from 96h (stage 2) onwards, when the bud has still a globular shape. Bar indicates color code for growth rates (um3 per hour). The untracked cells (not generating the cells at 132h) are marked as having now growth (dark blue). (**B**) Forward (23h to55h) and backward growth rate (55h - 23h) of cells in flower meristem 2 (FM2), also showing predetermined growth rates from globular stage onward. (**C)** Forward and backward growth rates in flower meristem 6 (FM6) also showing predetermined growth from globular stage onwards. Color codes in (B) and (C) as in (A).

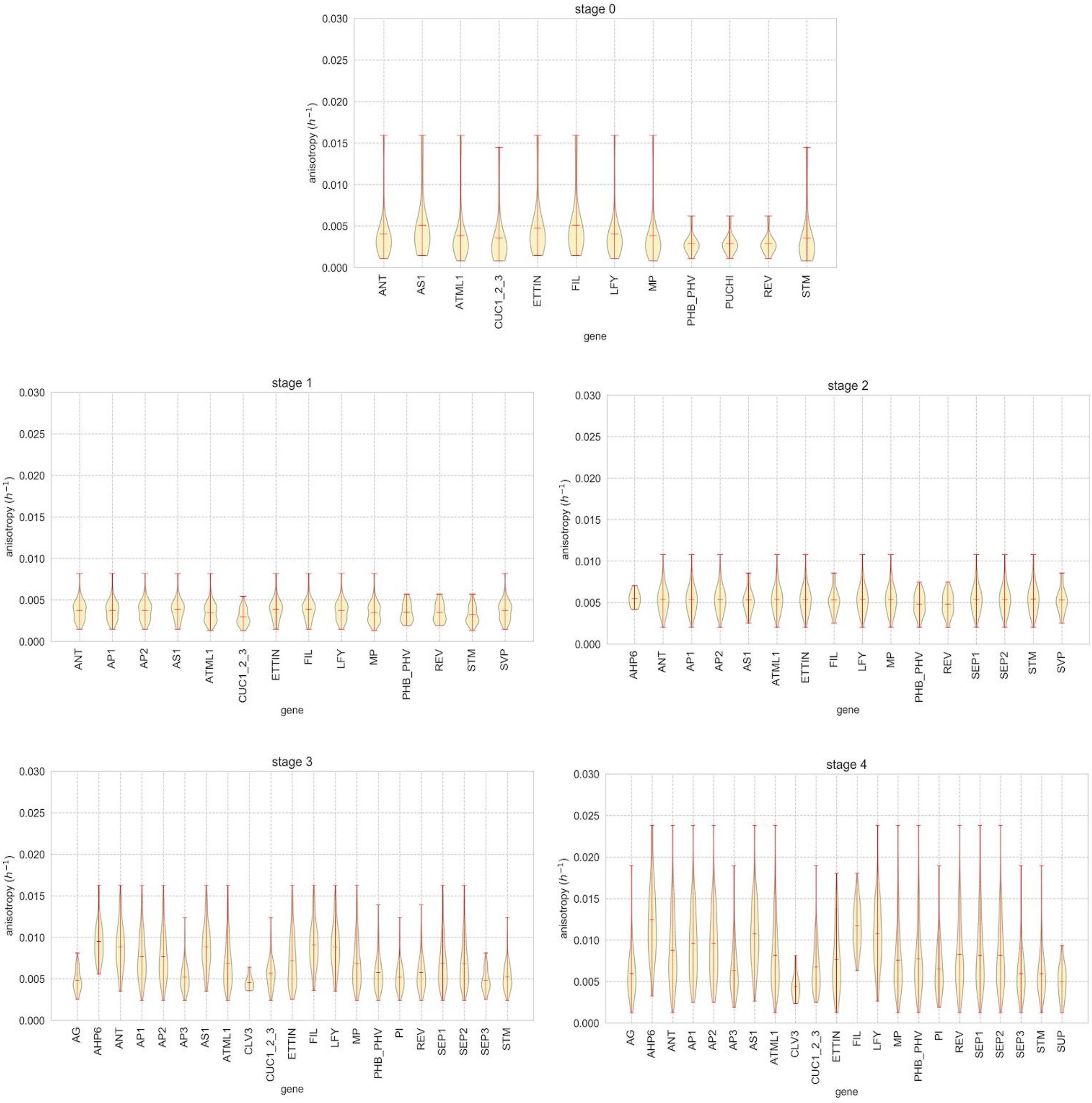

**Figure S11** Growth anisotropy for gene expression domains at different stages of flower development

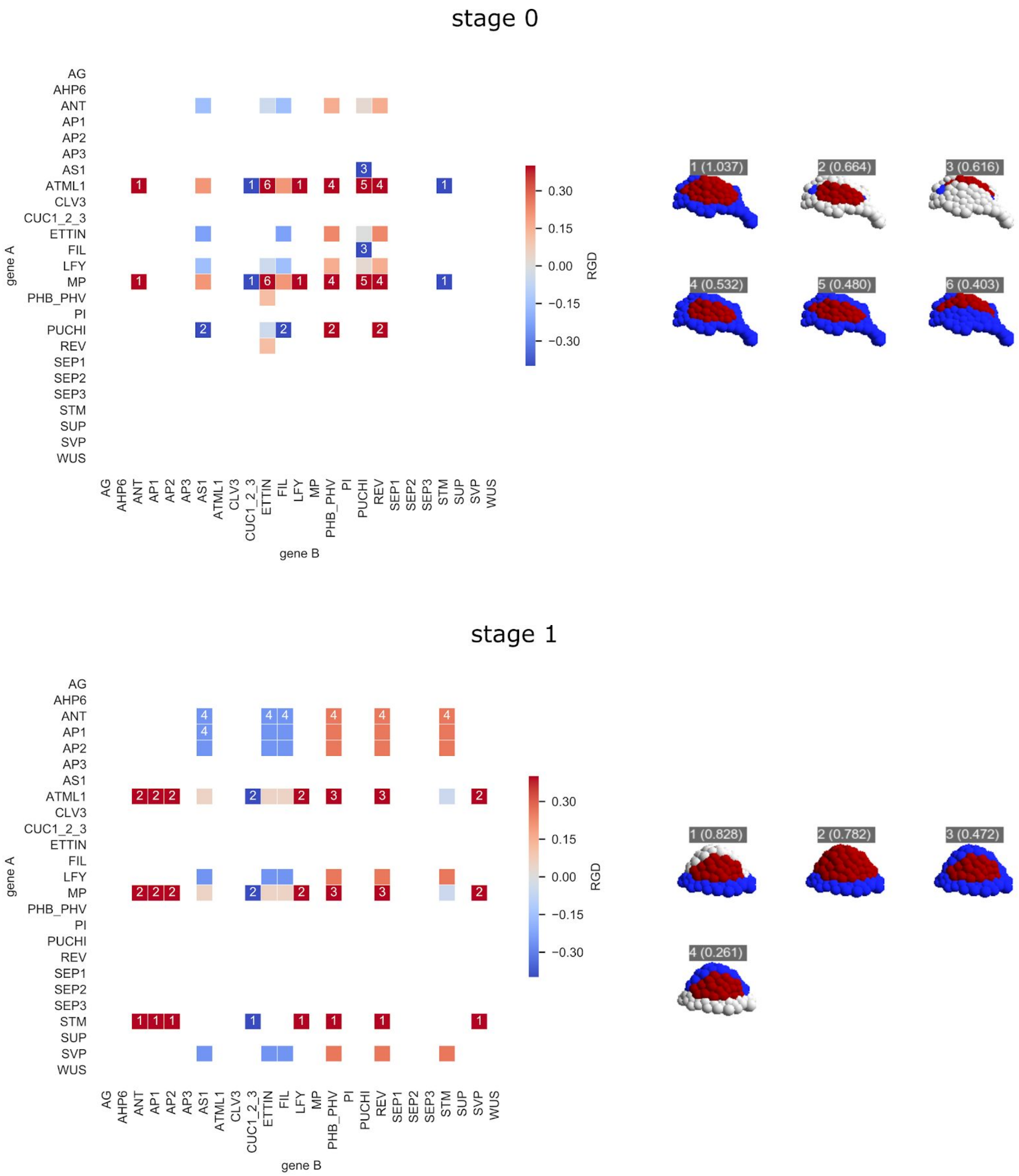

**Figure S12** Relative Growth Differences (RGD) between gene pairs (left) and the regions they define (right) for stages 0 and 1**.** For each pair of genes on the heatmap the colour refers to the RGD between the mean growth rate of the population of cells co-expressing gene A and B versus the mean growth rate of the population of cells expressing only A. The RGD for pairs of genes where either population is empty (i.e completely overlapping or not overlapping at all) is not reported (blank cells). The numbered annotations refer to the region separation implied by the gene pairs. The regions are shown along with their RGD in parentheses on the tissue geometry sorted by RGD where the region with the higher mean growth rate of the two is shown in red and the region with the lower growth rate is shown in blue.

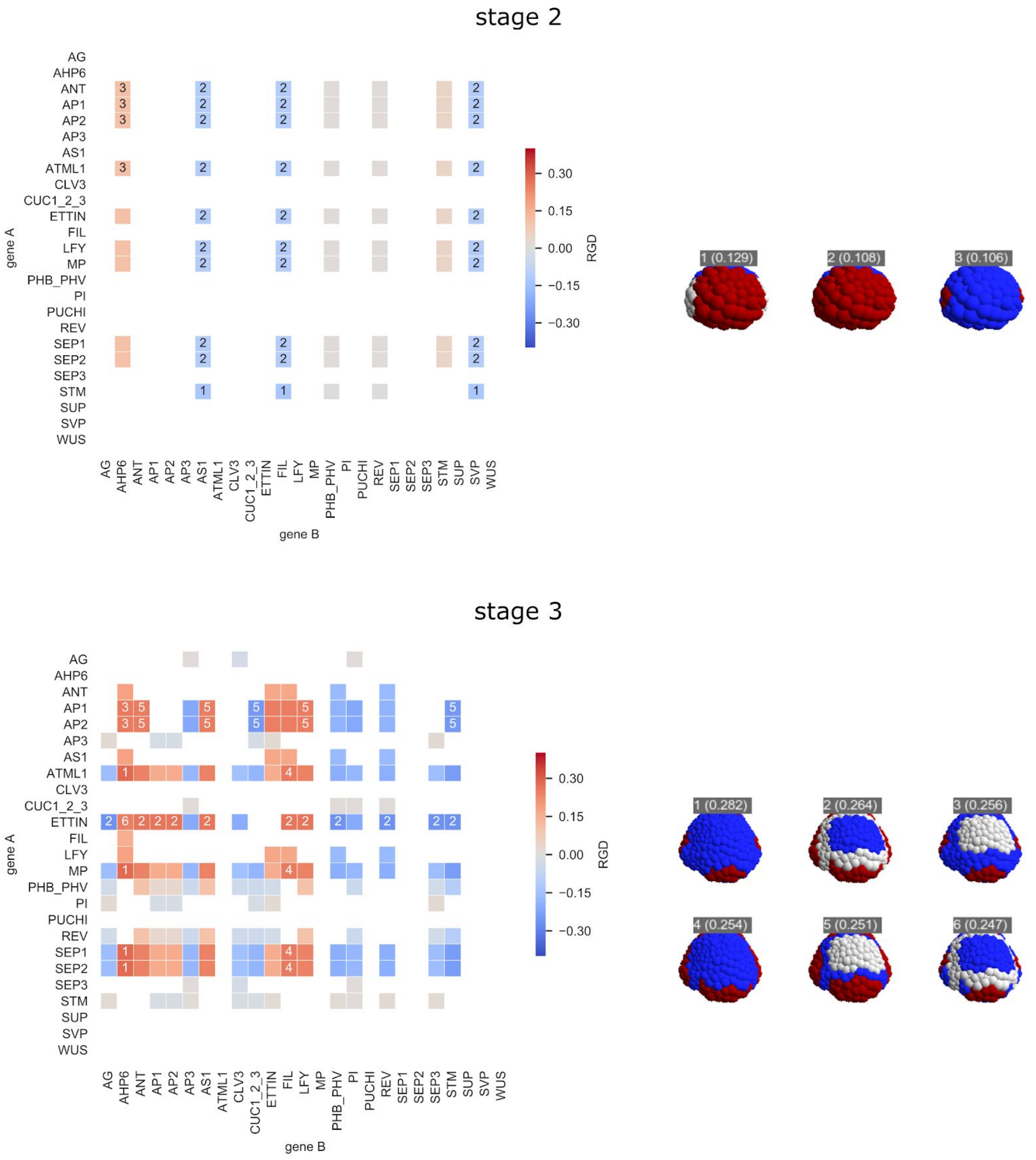

**Figure S13** Relative Growth Differences (RGD) between gene pairs (left) and the regions they define (right) for stages 2 and 3. Colours and annotations as for Fig. S11.

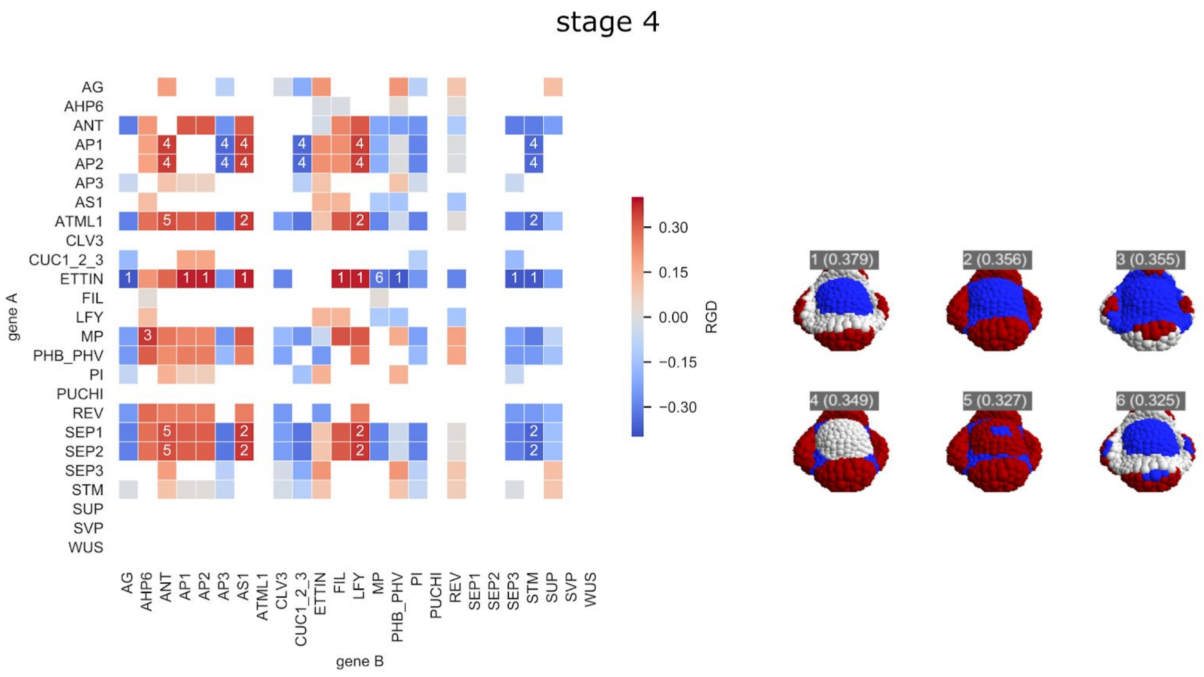

**Figure S14** Relative Growth Differences (RGD) between gene pairs (left) and the regions they define (right) for stage 4. Colours and annotations as for Fig. S11.

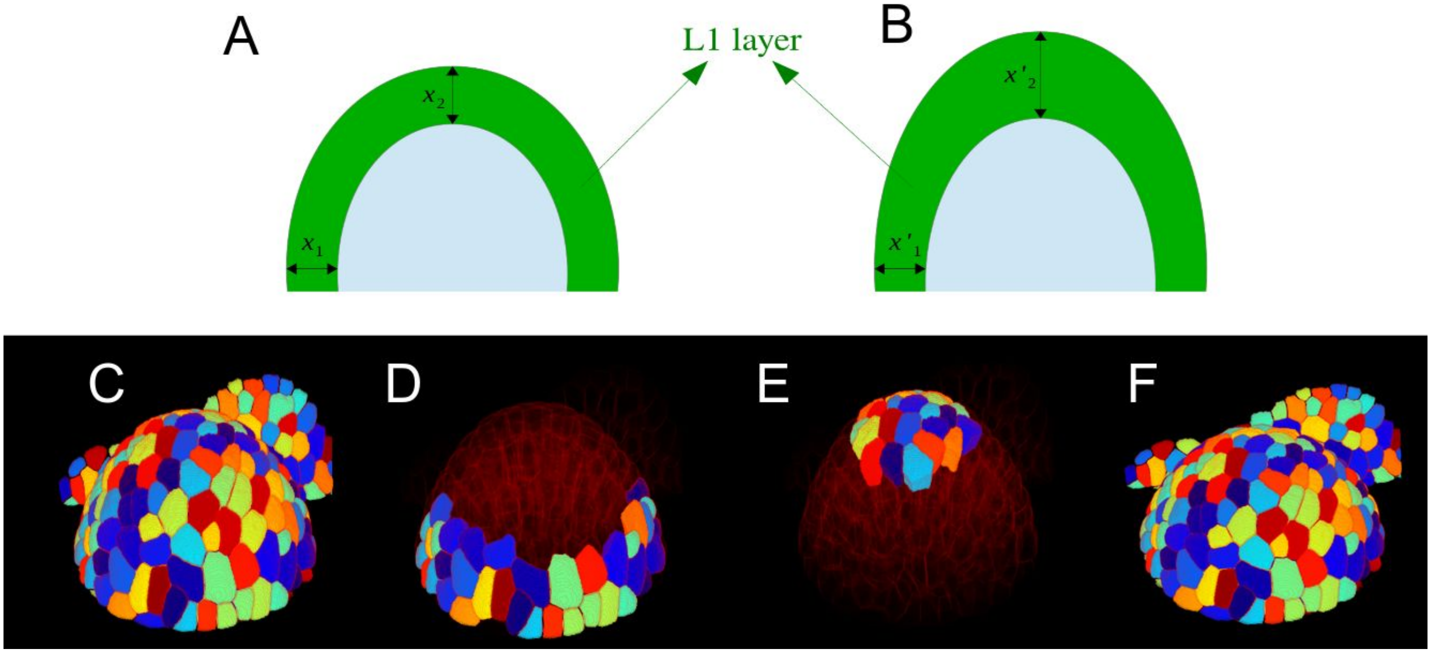

**Figure S15** Estimation of image resolution using L1 layer thickness. **A & B )** Schematic image of unstretched (A) and stretched primordium due to plantlets movement in z-direction (B). We assume that L1 layer thickness is constant in the absence of the movement in z-direction. We call $x'_2/x'_1$ stretching factor, where $x'_1$ is the L1 layer thickness not affected by the movement and $x'_2$ is the L1 layer thickness most affected by the movement **C**) Stretched primordium **D**) Cells least-affected by the movement in z-direction **E**) Cells most-affected with the movement in z-direction **F**) Primordium after correction of stretching factor.

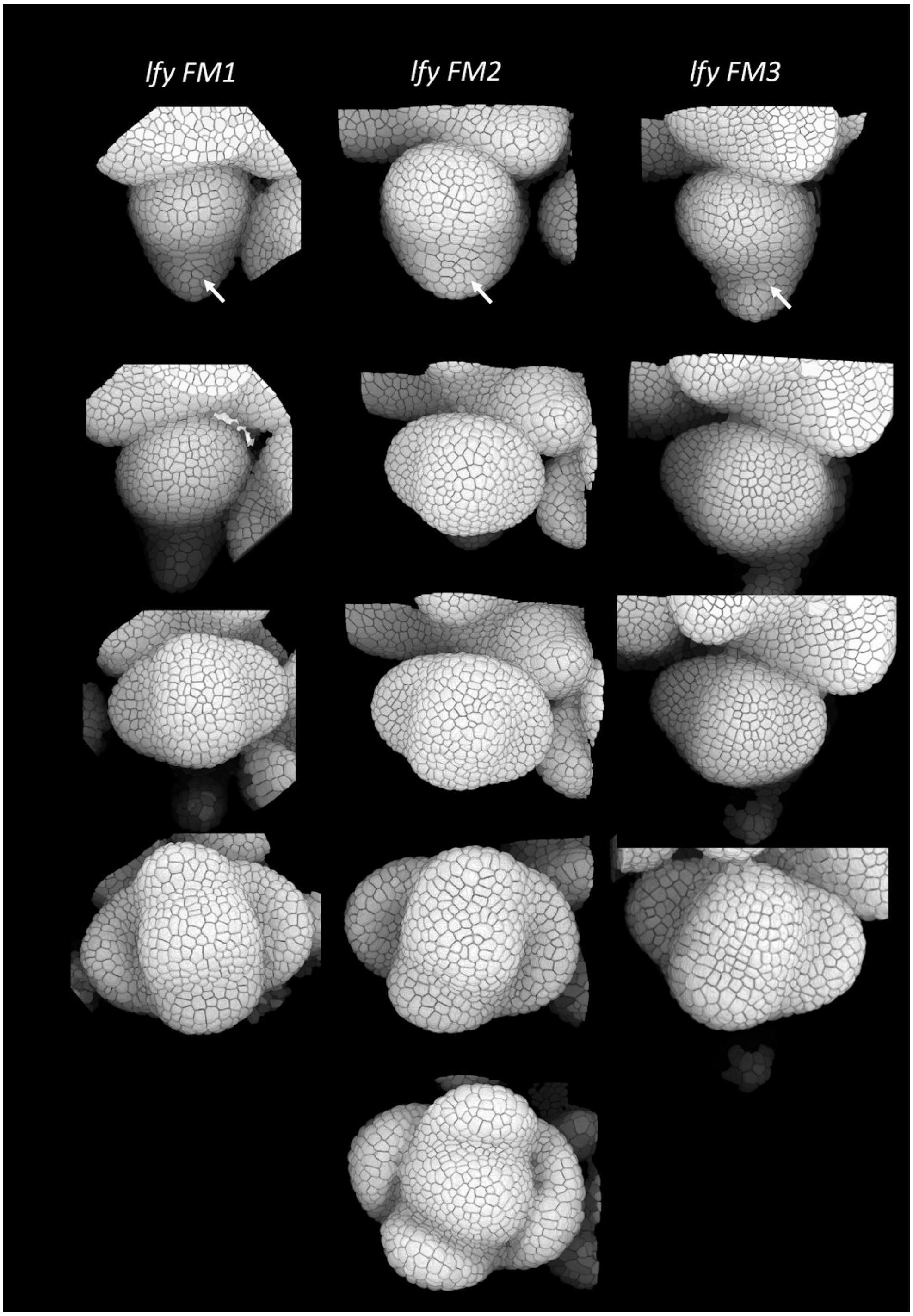

**Fig S16.** Three examples of early development in *lfy-12*. FM1 was analyzed in detail (see main text). Arrows point at outgrowing bracts (on the abaxial side of the flowers. In all three cases, the lateral sepals grow out first, followed by the abaxial and adaxial ones. Note that phyllotaxis is slightly perturbed. This is particularly visible in the last time-point of *lfy* FM2, where the adaxial sepal is not perpendicularly positioned to the meristem radius, suggesting a tendency towards spiralled phyllotaxis.

**Table 1: Justification gene interactions :**

| **Meristem function** | | |
|---|---|---|
| *AUX (via MP/FIL) ¬ STM* | *(Vernoux et al., 2000) (Chung et al., 2019; Vernoux et al., 2000)* | *Genetic evidence, and in situ hybridisation. STM is ectopically expressed in the pin1 mutant which has reduced auxin levels at the meristem. This inhibition might at least in part involve MP and its target FIL.* |
| CLV3 ¬ WUS | (Mayer et al., 1998; Schoof et al., 2000) | Genetic evidence (opposing phenotypes and epistasy of *wus* over *clv3* mutant, reciprocal effect of mutations on each others' expression patterns). CLV3 acts via CLV1 receptor. |
| CUC > STM | ((Mayer et al., 1998; Schoof et al., 2000; Takada et al., 2001); (Hibara et al., 2006); (Spinelli et al., 2011) | Partial overlap in expression. Ectopic CUC1 expression induces ectopic STM. Mutants have reduced STM expression. |
| ETT ¬ WUS | (Liu et al., 2014) | ETT and WUS overlap in the early flower. Yeast 1H and ChIP binding of ETT to pWUS. Prolonged WUS (in situ) signal in flowers of *ett/arf3* mutants. |
| ETT ¬ STM | (Chung et al., 2019) | STM is upregulated in *ett/arf4/fil* triple mutant. ETTIN binds directly to the STM promoter. Inactivation involves histone-deacetylation mediated transcriptional silencing |
| FIL ¬ STM | (Chung et al., 2019) | STM is upregulated in *ett/arf4/fil* triple mutant. FIL binds directly to the *STM* promoter. Inactivation involves epigenetic processes. |

| | | |
|---|---|---|
| STM > CUC1-3 | (Spinelli et al., 2011) | Ectopic STM induces CUC1 and later on CUC2-3. CUC1 has putative STM binding sites |
| WUS > CLV3 | (Mayer et al., 1998; Schoof et al., 2000; Takada et al., 2001; Yadav et al., 2011) | Genetic evidence (opposing phenotypes and epistasy of *wus* over *clv3* mutant, reciprocal effect of mutations on each others' expression patterns). CLV3 acts via CLV1 receptor. WUS directly binds to *CLV3* promoter. |
| **Organ outgrowth** | | |
| AUX > MP | (Wenzel et al., 2007)see e.g. (Vernoux et al., 2011) for general overview. | Post-transcriptional regulation. MP is an ARF (ARF5), which is activated by AUXIN via degradation of AUX/IAA. |
| ANT > LFY | (Hirota et al., 2007; Yamaguchi et al., 2016) | ANT with AIL6 induces *LFY* in parallel with MP. Delayed activation of *LFY* in *ant/ail6* mutants. Inducible ANT expression induces LFY in *ant.* ANT and AIL6 are expressed slightly prior to LFY and bind directly to promoter (ChIP qPCR). |
| AUX > PUCHI | (Hirota et al., 2007; Yamaguchi et al., 2016) | pPUCHI has aux-re elements required for its expression, and auxin increases PUCHI levels. |
| MP > AHP6 | (Besnard et al., 2014; Bishopp et al., 2011) | AHP6 has several Auxin responsive elements in its promoter. AHP6 is not activated in *mp* mutants. |
| MP > ANT | (Yamaguchi et al., 2013) | Reduced expression in *mp* mutant, induction of expression after inducible overexpression, direct binding of MP to promoter (CHiP qPCR). |
| MP > LFY | (Yamaguchi et al., 2013) | Reduced expression in *mp* mutant, induction of |

| | | |
|---|---|---|
| | | expression after inducible overexpression, direct binding of MP to promoter. |
| PUCHI > AUX | (Chandler and Werr, 2017) | Synergy with DRN/DRNL/BOP genes in controlling Auxin biosynthesis and promoting auxin transport. |
| LFY > CUC | (Yamaguchi et al., 2014) | Reduced expression of CUC in lfy mutant, induction in LFY-GR line, direct binding. |
| **Organ identity** | | |
| AG/SEP3 ¬ AP1 | (Gustafson-Brown, 1994; Kaufmann et al., 2009; Ó'Maoiléidigh et al., 2014; Pajoro et al., 2014; Sakai et al., 2000) | Genetic interactions and in situ hybridisation. AP1 is repressed in inner two whorls when AG is expressed there. Confirmed by CHiP seq. Probably in combination with SEP3, which also binds to AP1 promoter. |
| AG ¬ AP2 | (Grigorova et al., 2011; Mizukami and Ma, 1992; Wollmann et al., 2010; Zhao et al., 2007) | Likely indirect, the precise relationship is not known. The genes are not co-expressed (Wollmann et al) and have an antagonistic effect. |
| AG > AP3 | (Gustafson-Brown, 1994; Ó'Maoiléidigh et al., 2014); reviewed in: (Kaufmann et al., 2009; Pajoro et al., 2014; Sakai et al., 2000); (Pajoro et al., 2014; Sakai et al., 2000) | Genetic evidence (mutant phenotypes) and CHiPseq |
| AG > SEP3 | (Gustafson-Brown, 1994; Ó'Maoiléidigh et al., 2014) (Kaufmann et al., 2009) | Genetic evidence (mutant phenotypes) and CHiPseq |
| AG > SUP | (Pajoro et al., 2014; Sakai et al., 2000) | Expression studies in mutants (AG required to maintain expression of SUP), CHiPseq. |
| AG ¬ WUS | (Lenhard et al., 2001; Lohmann et al., 2001) | WUS expression is maintained in ag mutants. AG directly binds to pWUS and inhibition |

| | | |
|---|---|---|
| | | involves polycomb group proteins. |
| AP1 > LFY | (Ferrándiz et al., 2000; Kaufmann et al., 2010; Liljegren et al., 1999); reviews: (Liu and Mara, 2010; Liu et al., 2007; Pajoro et al., 2014) | AP1 overexpression prematurely induces LFY (Liljegren), LFY downregulation in *ap1*-mutants (Ferrandiz), confirmed by CHiPseq (e.g. Kaufmann) |
| SVP ¬AP3 (early development)<br><br>AP1/SEP3 > AP3 | (Gregis et al., 2009; Kaufmann et al., 2009; Ng and Yanofsky, 2001) (Kaufmann et al., 2009; Liu and Mara, 2010; Liu et al., 2007; Ng and Yanofsky, 2001; Pajoro et al., 2014); | AP3 induced by AP1-VP16, reduced AP3 expression in *ap1* mutant. Direct binding to pAP3.<br>Inhibitory action proposed when combined with SVP or SOC1 during early flower development. Direct binding of AP1 to AP3 promoter. To avoid contradiction the effect of AP1 during early development was represented as a negative regulation by SVP |
| SVP ¬PI (early development) | (Gregis et al., 2009; Kaufmann et al., 2009; Ng and Yanofsky, 2001) (Kaufmann et al., 2009; Liu and Mara, 2010; Liu et al., 2007; Ng and Yanofsky, 2001; Pajoro et al., 2014); | Inhibitory action proposed for AP1 when combined with SVP or SOC1 during early flower development. Direct binding of AP1 to PI3 promoter. To avoid contradiction this was represented as a negative regulation by SVP |
| AP1/SEP3 > PI | (Kaufmann et al., 2009; Ng and Yanofsky, 2001)) reviewed in : (Liu and Mara, 2010; Liu et al., 2007; Pajoro et al., 2014); (Gregis et al., 2006, 2009) | PI induced by AP1-VP16, reduced expression in *ap1* mutants (Ng and Yanofsky). Direct binding of AP1 to pPI. This light require SEP3 (Gregis et al 2009) |
| AP1 ¬ AG | (Gregis et al., 2006, 2009) Discussed in: (Drews et al., 1991; Grigorova et al., 2011) | Both genes have complementary expression patterns. AP1, together with SVP and AGL24 control AG expression (phenotype in triple mutant shows carpelloid structures in 1st |

| | | |
|---|---|---|
| | | whorl). AP1 binds to pAG (ChIP) |
| SVP ¬ SEP3 (stage 1 and stage 2) | (Kaufmann et al., 2009, 2010) (Gregis et al., 2006, 2008, 2009); (Kaufmann et al., 2009; Liu et al., 2009); review: (Liu and Mara, 2010; Liu et al., 2007; Pajoro et al., 2014) | AP1 binds directly to pSEP3 (ChIP) and inhibits SEP3 during stage 1 and 2 with SVP). Lateron AP1 interacts with SEP3 to induce B genes. CHiP data and genetic/phenotypic analysis, in situ hybridisation.<br>To avoid contradiction in the model, we represented the negative regulation as an input from SVP |
| AP1 > SEP3 | (Kaufmann et al., 2009, 2010) see also: (Gregis et al., 2006, 2008, 2009) | AP1 binds to the SEP3 promoter, and SEP3 expression is rapidly up-regulated after AP1 activation using an inducible system. |
| AP1 ¬ SOC1 SVP (with SEP3 during later stages) | (Liu et al., 2007) | Increased expression in *ap1* mutant and direct binding to promoters (CHiP) |
| AP2 ¬ AG | (Drews et al., 1991; Grigorova et al., 2011; Yant et al., 2010)) | Expression of *AG* in all whorls of *ap2* mutants. Binding of AP2 to pAG (CHiP qPCR). |
| AP2 ¬ AP1 | (Liu and Mara, 2010; Liu et al., 2007; Pajoro et al., 2014; Yant et al., 2010) | CHiP seq data indicate binding, Yant et al conclude a negative interaction, but this seems in contradiction with their co-expression. |
| AP2 ¬ ETT | (Liu et al., 2014; Yant et al., 2010) | pETT has AP2 binding sites, confirmed by ChIP qPCR. Increased ARF3 expression in *ap2* mutant, reduced in inducible overexpresser. |
| AP2 > PI | (Goto and Meyerowitz, 1994; Liu et al., 2014; Yant et al., 2010) | CHiP seq (Yant et al), lower expression levels, but normal pattern in *ap2* flower (in situ hybridisation) |
| AP3/ PI > SUP | (Sakai et al., 2000); (Wuest et al., 2012) | Lack of SUP expression in mutants. CHiP seq. Reduction |

| | | |
|---|---|---|
| | | of expression in AP3 inducible knock-down. |
| AP3/PI ¬ AP1 | (Sundström et al., 2006) (Wuest et al., 2012) | ChIP seq. Inducible *AP3/PI* expression, shows reduced *AP1* expression. *ap3/pi* mutants show increased *AP1* levels. |
| AP3 > PI | (Wuest et al., 2012) | Reduction of PI in inducible knock down. CHiP seq |
| LFY > AP1 | (Parcy et al., 1998); (Wagner et al., 1999); Review: (Liu and Mara, 2010; Liu et al., 2007; Pajoro et al., 2014) | In situ hybridisations, mutant phenotypes, LFY-VP16 induction of AP1. |
| LFY > AP3 | (Parcy et al., 1998) | In situ hybridisations, mutant phenotypes, LFY-VP16 induction of AP3. |
| LFY > PI | (Kaufmann et al., 2009; Ng and Yanofsky, 2001) (Winter et al., 2011) | LFY required for PI expression. Relation suggested direct with UFO. CHiP suggests direct binding. |
| LFY > AG | (Lohmann et al., 2001); (Parcy et al., 1998); review: (Liu and Mara, 2010; Liu et al., 2007; Pajoro et al., 2014) | In situ hybridisations, mutant phenotypes, LFY-VP16 induction of AG. |
| PUCHI > LFY | (Karim et al., 2009) | lfy phenotype in *puchi/bop1/bop2* mutant. No LFY expression. |
| LFY/SEP (1-)3 > AP3 | (Kaufmann et al., 2009) (Liu et al., 2009) review: (Liu and Mara, 2010; Liu et al., 2007; Pajoro et al., 2014) | CHiCHIP (Kaufmann et al) for SEP3, AP3 reduced expression in sep1-4 quadruple mutant (Liu et al 2009). However, not in sep1-3 triple (Pelaz et al 2000). SEP3 binds to LFY. |
| SEP (3) > AG | (Kaufmann et al., 2009); review: (Liu and Mara, 2010; Liu et al., 2007; Pajoro et al., 2014) | ChiP-CHIP (Kaufmann et al), AG reduced expression in sep1-4 quadruple mutant (Liu et al 2009). However, not in sep1-3 triple (Pelaz et al 2000). SEP3 binds to LFY. |

| | | |
|---|---|---|
| SVP ¬ SEP 3 | (Kaufmann et al., 2009; Liu et al., 2009); review: (Liu and Mara, 2010; Liu et al., 2007; Pajoro et al., 2014)<br>For further details on SVP: see also: (Gregis et al., 2013; Hartmann et al., 2000; Liu et al., 2013) | ChIP (Kaufmann et al), Genetic evidence combined with in situ hybridisations (Liu et al.). |
| SUP ¬ AP3 PI | (Liu and Mara, 2010; Liu et al., 2007; Pajoro et al., 2014; Prunet et al., 2017; Sakai et al., 2000); (Sakai et al., 2000) | Extended AP3/PI expression in mutants (review: Liu and Mara), due to reconversion of 4>th whorl cells to 3rd whorl (Prunet et al 2017). Way of action unknown. |
| WUS > AG | (Lenhard et al., 2001; Lohmann et al., 2001) | AG is required for WUS repression and meristem termination. Ectopic WUS can induce AG, this involves direct binding (tests using GUS with AG intron in ectopic WUS expressing lines, …) |
| **Organ polarity** | | |
| ANT > FIL | (Nole-Wilson and Krizek, 2006) | ANT directly binds FIL promoter, but no reduced expression of FIL in *ant* mutant. Slightly induction of FIL in inducible overexpressing line (RNAseq) |
| AS1 > REV | (Fu et al., 2007) | Reduced expression in *as1* mutant. Increased expression in as1 overexpresser and dominant gain of AS2 function |
| DOF5.1 > REV | (Kim et al., 2010) | Activation tagging (35S): induction of REV, DOF binds directly to REV promoter. DOF 5.1 is expressed in the entire apex. |
| AS2 ¬ KAN | (Iwakawa et al., 2007; Iwasaki et al., 2013; Nurmberg et al., 2007; Wu et al., 2008)) | Plants transformed with35S:AS2 have reduced levels of KAN1 and KAN2mRNA and have a leaf phenotype reminiscent of the |

| | | |
|---|---|---|
| | | kan1 kan2 double mutant, whereasas2–1mutants have elevated levels ofKAN2 |
| AS1(with AS2)¬ ETT | (Iwakawa et al., 2007; Iwasaki et al., 2013; Nurmberg et al., 2007; Wu et al., 2008) (Husbands et al., 2015); review: (Kuhlemeier and Timmermans, 2016) | AS1 and AS2 act in a complex. AS1 and 2 bind directly to the ETT promoter, and ETT is maintained longer in the as2 mutant. ETTIN expression increased in as1 mutant. The complex might also act via the tasiARF pathway |
| AS1(with AS2) > PHB PHV | (Fu et al., 2007) | Reduced expression in *as1* mutant. Increased expression in AS1 overexpresser and dominant gain of AS2 function |
| FIL > KAN | (Bonaccorso et al., 2012); see also: (La Rota et al., 2011) | 35S::FIL:GR induces expression of KAN even in the presence of CHX. Expression greatly reduced in *fil yab2 yab3 yab5.* |
| FIL > AS1 | (Bonaccorso et al., 2012) | 35S::FIL:GR induces expression of AS1, even in the presence of CHX , note, however, that expression is not altered in *fil yab2 yab3 yab5* |
| *KAN > FIL* | *(Bonaccorso et al., 2012)* | *No clear evidence for direct interactions. Evidence based on their complementary expression patterns and roles.* |
| KAN ¬ PHB/PHV (directly and via MiR 166A) | (Eshed et al., 2001; Kuhlemeier and Timmermans, 2016; Pekker et al., 2005)see also (Eshed et al., 2001; Kuhlemeier and Timmermans, 2016; Pekker et al., 2005; Zhou et al., 2007); (Merelo et al., 2013). Reviews : (Emery et al., 2003); (Kuhlemeier and Timmermans, 2016) | KAN binds directly to the miR166A and F promoters, and to pPHV, and downregulates these genes (in 3SS::KAN-GR line) (Merelo et al).<br><br>Note: Overexpression of miR 165 reduces the expression of PHB/PHV, and causes phenotypes reminiscent of HDZIP III mutants (Zhou et al). |

| | | |
|---|---|---|
| LFY > ETT | (Yamaguchi et al., 2014) | Reduced ETT in *lfy* mutant, reduced ETT in *pin1* mutant, induction in LFY-GR line. Binding to p*ETT* |
| KAN¬ AS2 (and the AS1AS2 complex) | (Wu et al., 2008) | 35S KAN GR reduces AS2 expression (but not AS1). pAS2 binds KAN and mutating the KAN binding site causes AS2 overexpression. Since AS2 acts in a complex with AS1, this also affects the activity of AS1. |
| MP > FIL | (Wu et al., 2015); (Chung et al., 2019) | Expression of FIL reduced in *mp* mutant. FIL induced by Inducible MP. Direct binding in ChIP. |
| *PHB/PHV/REV ¬ KAN* | (Pekker et al., 2005); (Emery et al., 2003) | In plants where *HD-ZIPIII* genes are ectopically expressed in the abaxial leaf domain, *KAN1* expression is reduced and vice versa. Suggested in different articles the discussion. Only indirect evidence (mutant phenotypes, effects of REV on KAN3, ...). The mechanism remains unclear and the effects concern rather downstream events. |
| *Sterol > PHB PHV & REV* | *(McConnell et al., 2001); (Eshed et al., 2001) review: (Emery et al., 2003)* | *The HD ZIPIII genes have a sterol binding domain, but this sterol has not been identified and remains hypothetical.* |
| *STM ¬ AS1* | (Byrne et al., 2000) (Scofield et al 2014) | Genes not co-expressed, AS1 domain extended in the *stm* mutant. Mechanism of inhibition in the meristem unknown, but probably very indirect: STM overexpression does not repress AS1. |

| Domain | Detailed description | General description | Organ identity |
|---|---|---|---|
| 2 | Abaxial floral meristem | Floral meristem | Undifferentiated |
| 3 | Floral meristem initium stage | | |
| 6 | Floral meristem stage 1(L2) |  |  |
| 7 | Floral meristem stage 1 (L1) | | |
| 10 | Floral meristem stage 2 | | |
| 18 | Sepal, adaxial domain | Adaxial domain organ | Sepal |
| 26 | Sepal tip, adaxial domain |  |  |
| 21 | Basal boundary between sepals stage 3 (L2) | Boundary | Boundary |
| 12 | Adaxial domain flower primordium | Primordium | Flower |
| 8 | Centre future lateral sepal | Lateral initium | Sepal |
| 11 | Periphery future lateral sepal | | |
| 28 | Sepal abaxial domain stage 3 (L2) | Abaxial domain organ | |
| 27 | Sepal abaxial domain stage 4 (L1) | | |
| 16 | Sepal tip, abaxial domain | | |
| 17 | Sepal, abaxial domain stage 3 | | |
| 9 | Abaxial domain flower primordium | Flower primordium | Primordium |
| 1 | Bract initium stage | bract | Bract |
| 31 | Lateral domain bract initium stage | | |
| 5 | Bract stage 1 | | |
| 4 | Boundary young flower primordium | Boundary | Boundary |
| 20 | Boundary between sepals stage 3 | | |
| 30 | Boundary between sepals 4 | | |
| 29 | Petal precursors, sepal boundary | Organ precursors, boundary | Petal/ |
| 19 | Petal and Stamen precursors, boundary | Organ precursors and boundaries | Petal/stamen |
| 25 | Stamen precursor stage 4, sepal boundary | | Stamen |
| 13 | Stamen precursor stage 1, sepal boundary | | |
| 22 | Stamen precursor stage 4, meristem | Organ precursors/meristem | |
| 24 | Boundary between stamen and carpel, meristem | Boundary | Boundary |
| 15 | Carpel precursor stage 3, periphery meristem | Organ precursors and meristem | Carpel |
| 23 | Carpel precursor stage 4, meristem | | |
| 14 | Carpel precursor stage 4, central meristem, | | |

Supplementary Table S2: List of cell states and clusters with the description of their identity.

**Table S3: Hypotheses from the literature.**

| Gene | Stage 0 (10h) | Stage 1 (40h) | Stage 2 (96h) | Stage 3 (120h) | Stage 4 (132h) | Summ. |
|---|---|---|---|---|---|---|
| AG | LFY∨WUS∧SEP3<br>¬AP1/2<br>1.0<br>LFY∨SEP3∧WUS<br>¬AP1/2<br>1.0<br>WUS∨SEP3∧LFY<br>¬AP1/2<br>1.0<br>LFY∧WUS∨SEP3<br>¬AP1/2<br>1.0<br>LFY∧SEP3∨WUS<br>¬AP1/2<br>1.0<br>(+1 others) | LFY∨WUS∨SEP3<br>¬AP1/2<br>1.0<br>LFY∨WUS∧SEP3<br>¬AP1/2<br>1.0<br>LFY∨SEP3∧WUS<br>¬AP1/2<br>1.0<br>WUS∨SEP3∧LFY<br>¬AP1/2<br>1.0<br>LFY∧WUS∨SEP3<br>¬AP1/2<br>1.0<br>(+3 others) | LFY∨WUS∨SEP3<br>¬AP1/2<br>1.0<br>LFY∨WUS∧SEP3<br>¬AP1/2<br>1.0<br>LFY∨SEP3∧WUS<br>¬AP1/2<br>1.0<br>WUS∨SEP3∧LFY<br>¬AP1/2<br>1.0<br>LFY∧WUS∨SEP3<br>¬AP1/2<br>1.0<br>(+3 others) | LFY∨WUS∨SEP3<br>¬AP1/2<br>0.5<br>LFY∨WUS∧SEP3<br>¬AP1/2<br>0.5<br>LFY∨SEP3∧WUS<br>¬AP1/2<br>0.5<br>WUS∨SEP3∧LFY<br>¬AP1/2<br>0.5<br>LFY∧WUS∨SEP3<br>¬AP1/2<br>0.5<br>(+3 others) | LFY∨WUS∨SEP3<br>¬AP1/2<br>1.0<br>LFY∧WUS∨SEP3<br>¬AP1/2<br>1.0 | LFY∧WUS∨SEP3<br>¬AP1/2<br>(10h, 40h, 96h, 120h, 132h) |
| AHP6 | MP<br>0.663 | MP<br>0.554 | MP<br>0.5 | MP<br>0.5 | MP<br>0.58 | MP<br>(10h, 40h, 96h, 120h, 132h) |
| ANT | MP<br>0.769 | MP<br>0.638 | MP<br>1.0 | MP<br>0.5 | MP<br>0.398 | MP<br>(10h, 40h, 96h, 120h, 132h) |
| AP1/2 | LFY∧PUCHI<br>¬AG<br>0.878<br>LFY∨PUCHI<br>¬AG<br>0.803 | LFY∨PUCHI<br>¬AG<br>1.0 | LFY∨PUCHI<br>¬AG<br>1.0 | LFY∨PUCHI<br>¬AG<br>0.725 | LFY∨PUCHI<br>¬AG<br>0.881 | LFY∨PUCHI<br>¬AG<br>(10h, 40h, 96h, 120h, 132h) |
| AP3 | LFY∨AP∧SEP3<br>¬SUP<br>1.0<br>LFY∨SEP3∧AP1/2<br>¬SUP<br>1.0<br>AP∨SEP3∧LFY<br>¬SUP<br>1.0<br>LFY∧AP∨SEP3<br>¬SUP<br>1.0<br>LFY∧SEP3∨AP1/2<br>¬SUP<br>1.0<br>(+1 others) | LFY∨AP∧SEP3<br>¬SUP<br>1.0<br>LFY∧AP∧SEP3<br>¬SUP<br>1.0 | LFY∨AP∧SEP3<br>¬SUP<br>1.0<br>LFY∧AP∧SEP3<br>¬SUP<br>1.0 | LFY∨AP∧SEP3<br>¬SUP<br>0.644 | LFY∨AP∨SEP3<br>¬SUP<br>0.595 | SEP3∨LFY∨AP1/2<br>¬SUP<br>(132h) |

| | | | | | | |
|---|---|---|---|---|---|---|
| AS1 | REV∨FIL<br>¬STM<br>0.863 | REV∨FIL<br>¬STM<br>1.0 | REV∨FIL<br>¬STM<br>0.5<br>REV∧FIL<br>¬STM<br>0.5 | REV∨FIL<br>¬STM<br>1.0 | REV∨FIL<br>¬STM<br>1.0 | FIL∨REV<br>¬STM<br>(10h, 40h, 96h, 120h, 132h) |
| CLV3 | WUS<br>1.0 | WUS<br>1.0 | WUS<br>1.0 | WUS<br>0.5 | WUS<br>0.5 | WUS<br>(10h, 40h, 96h, 120h, 132h) |
| CUC1_3 | STM∨LFY<br>0.726 | STM∨LFY<br>0.576 | STM∧LFY<br>0.611 | STM∨LFY<br>0.5<br>STM∧LFY<br>0.5 | STM∨LFY<br>0.5<br>STM∧LFY<br>0.5 | STM∧LFY<br>(96h, 120h, 132h) |
| ETT | AP∨LFY<br>¬AS1∧REV<br>0.92 | AP∨LFY<br>¬AS1∧REV<br>0.79<br>AP∧LFY<br>¬AS1∧REV<br>0.79 | AP∨LFY<br>¬AS1∧REV<br>1.0<br>AP∧LFY<br>¬AS1∧REV<br>1.0 | AP∧LFY<br>¬AS1∧REV<br>0.767 | AP∧LFY<br>¬REV∧AS1<br>0.68 | LFY∧AP1/2<br>¬REV∧AS1<br>(40h, 96h, 120h, 132h) |
| FIL | ANT∨MP∧ETT<br>¬REV<br>1.0<br>ANT∨ETT∧MP<br>¬REV<br>1.0<br>MP∨ETT∧ANT<br>¬REV<br>1.0<br>ANT∧MP∨ETT<br>¬REV<br>1.0<br>MP∧ETT∨ANT<br>¬REV<br>1.0<br>(+1 others) | ANT∨MP∧ETT<br>¬REV<br>1.0<br>ANT∨ETT∧MP<br>¬REV<br>1.0<br>MP∨ETT∧ANT<br>¬REV<br>1.0<br>ANT∧MP∨ETT<br>¬REV<br>1.0<br>MP∧ETT∨ANT<br>¬REV<br>1.0<br>(+1 others) | ANT∨MP∨ETT<br>¬REV<br>0.741<br>ANT∨MP∧ETT<br>¬REV<br>0.741<br>ANT∨ETT∧MP<br>¬REV<br>0.741<br>MP∨ETT∧ANT<br>¬REV<br>0.741<br>ANT∧MP∨ETT<br>¬REV<br>0.741<br>(+3 others) | ANT∨MP∧ETT<br>¬REV<br>1.0<br>ANT∨ETT∧MP<br>¬REV<br>1.0<br>MP∨ETT∧ANT<br>¬REV<br>1.0<br>ANT∧MP∨ETT<br>¬REV<br>1.0<br>MP∧ETT∨ANT<br>¬REV<br>1.0<br>(+3 others) | MP∧ETT∨ANT<br>¬REV<br>0.94<br>ANT∨MP∧ETT<br>¬REV<br>0.904<br>MP∨ETT∧ANT<br>¬REV<br>0.904<br>ANT∧MP∨ETT<br>¬REV<br>0.904 | MP∧ANT∨ETT<br>¬REV<br>(10h, 40h, 96h, 120h, 132h)<br>ETT∨MP∧ANT<br>¬REV<br>(10h, 40h, 96h, 120h, 132h)<br>ETT∧MP∨ANT<br>¬REV<br>(10h, 40h, 96h, 120h, 132h)<br>ANT∨MP∧ETT<br>¬REV<br>(10h, 40h, 96h, 120h, 132h) |
| LFY | MP∨AP∨PUCHI∧ANT<br>1.0<br>ANT∨AP∨PUCHI∧MP<br>1.0<br>MP∨AP∧ANT∨PUCHI<br>1.0<br>MP∨AP∧PUCHI∨ANT<br>1.0<br>MP∨PUCHI∧ANT∨AP1/2<br>1.0<br>(+10 others) | MP∨ANT∨PUCHI∧AP1/2<br>1.0<br>MP∨AP∨PUCHI∧ANT<br>1.0<br>ANT∨AP∨PUCHI∧MP<br>1.0<br>MP∨ANT∧AP∨PUCHI<br>1.0<br>MP∨ANT∧PUCHI∨AP1/2<br>1.0<br>(+25 others) | MP∨ANT∨AP∨PUCHI<br>1.0<br>MP∨ANT∨PUCHI∧AP1/2<br>1.0<br>MP∨AP∨PUCHI∧ANT<br>1.0<br>ANT∨AP∨PUCHI∧MP<br>1.0<br>MP∨ANT∧AP∨PUCHI<br>1.0<br>(+33 others) | MP∨AP∨PUCHI∧ANT<br>1.0<br>MP∨AP∧ANT∨PUCHI<br>1.0<br>MP∨AP∧PUCHI∨ANT<br>1.0<br>MP∨PUCHI∧ANT∧AP1/2<br>1.0<br>ANT∨PUCHI∧MP∧AP1/2<br>1.0<br>(+10 others) | MP∧PUCHI∨ANT∧AP1/2<br>1.0<br>MP∧PUCHI∨AP∧ANT<br>1.0 | MP∧PUCHI∨ANT∧AP1/2<br>(40h, 96h, 120h, 132h)<br>MP∧PUCHI∨AP∧ANT<br>(40h, 96h, 120h, 132h) |
| PHB/ PHV | ANT∨AS1<br>0.893 | ANT∨AS1<br>0.773 | ANT∨AS1<br>0.5 | ANT∨AS1<br>0.236<br>ANT∧AS1<br>0.236 | ANT∨AS1<br>0.783 | AS1∨ANT<br>(10h, 40h, 96h, 120h, 132h) |

| | | | | | | |
|---|---|---|---|---|---|---|
| PI | LFY∨SEP3∧AP1/2<br>¬SUP<br>1.0<br>LFY∨AP∧SEP3<br>¬SUP<br>1.0<br>SEP3∨AP∧LFY<br>¬SUP<br>1.0<br>LFY∧SEP3∨AP1/2<br>¬SUP<br>1.0<br>LFY∧AP∨SEP3<br>¬SUP<br>1.0<br>(+1 others) | LFY∨AP∧SEP3<br>¬SUP<br>1.0<br>LFY∧SEP3∧AP1/2<br>¬SUP<br>1.0 | LFY∨AP∧SEP3<br>¬SUP<br>1.0<br>LFY∧SEP3∧AP1/2<br>¬SUP<br>1.0 | LFY∨AP∧SEP3<br>¬SUP<br>0.644 | LFY∨SEP3∨AP1/2<br>¬SUP<br>0.583<br>LFY∧AP∨SEP3<br>¬SUP<br>0.555 | AP∨LFY∨SEP3<br>¬SUP<br>(132h)<br>LFY∧AP∨SEP3<br>¬SUP<br>(132h) |
| REV | AS1<br>0.393 | AS1<br>0.273 | AS1<br>0.338 | AS1<br>0.236 | AS1<br>0.649 | AS1<br>(10h, 40h, 96h, 120h, 132h) |
| STM | CUC<br>¬AHP6<br>1.0 | CUC<br>¬AHP6<br>0.75 | CUC<br>¬AHP6<br>0.5 | CUC<br>¬AHP6<br>0.745 | CUC<br>¬AHP6<br>0.748 | CUC<br>¬AHP6<br>(10h, 40h, 96h, 120h, 132h) |
| SUP | AP3∨PI∨AG<br>1.0<br>AP3∨PI∧AG<br>1.0<br>AP3∨AG∧PI<br>1.0<br>PI∨AG∧AP3<br>1.0<br>AP3∧PI∨AG<br>1.0<br>(+3 others) | AP3∨PI∨AG<br>1.0<br>AP3∨PI∧AG<br>1.0<br>AP3∨AG∧PI<br>1.0<br>PI∨AG∧AP3<br>1.0<br>AP3∧PI∨AG<br>1.0<br>(+3 others) | AP3∨PI∨AG<br>1.0<br>AP3∨PI∧AG<br>1.0<br>AP3∨AG∧PI<br>1.0<br>PI∨AG∧AP3<br>1.0<br>AP3∧PI∨AG<br>1.0<br>(+3 others) | AP3∨PI∧AG<br>0.921<br>AP3∧PI∧AG<br>0.921 | AP3∧PI∨AG<br>0.821<br>AP3∨PI∨AG<br>0.766 | PI∨AP3∨AG<br>(132h)<br>PI∧AP3∨AG<br>(132h) |

**Table S3:** Summary of inputs for every gene at the 5 different time points (in hours), based on a literature search. Rules are combining logical AND (∧) and OR (∨), where negative regulation is implemented as logical NOT (¬). Optimal rules for individual time points are shown. The numbers below the rules indicate the Bacc score. Where alternative rules are equally good, several are listed. 'Summ' column indicates gene interactions with most significant impact throughout development (stages at which they occur indicated between brackets).

#### Table S4: novel hypotheses

| Gene | Stage 0 (10h) | Stage 1 (40h) | Stage 2 (96h) | Stage 3 (120h) | Stage 4 (132h) | Summ. |
|---|---|---|---|---|---|---|
| AG | AHP6 1.0<br>PHB 1.0<br>STM 1.0<br>CUC 1.0<br>ATML1 1.0<br>(+37 others) | AHP6 1.0<br>PHB 1.0<br>STM 1.0<br>CUC 1.0<br>ATML1 1.0<br>(+37 others) | AHP6 1.0<br>PHB 1.0<br>STM 1.0<br>CUC 1.0<br>ATML1 1.0<br>(+37 others) | ¬AHP6 1.0<br>¬CUC 1.0<br>¬AS1 1.0<br>¬ANT 1.0<br>¬PUCHI 1.0<br>(+4 others) | AHP6 1.0<br>PHB 1.0<br>STM 1.0<br>CUC 1.0<br>ATML1 1.0<br>(+36 others) | (¬CLV3)*<br>¬ANT<br>(120h, 132h)<br><br>*already in the literature via WUS |
| AHP6 | AP3 1.0<br>SUP 1.0<br>SEP1_2 1.0<br>PI 1.0<br>AG 1.0<br>(+7 others) | AP3 1.0<br>SUP 1.0<br>SEP1_2 1.0<br>PI 1.0<br>AG 1.0<br>(+5 others) | ¬STM 0.923 | FIL 0.885<br>¬REV 0.833<br>¬PHB 0.833<br>LFY 0.806<br>AS1 0.806<br>(+2 others) | LFY 0.871<br>AS1 0.871<br>¬STM 0.871<br>AP1/2 0.791<br>¬AG 0.791<br>(+1 others) | ¬STM<br>(96h, 120h, 132h) |
| ANT | LFY 1.0<br>¬STM 1.0<br>¬CUC 1.0 | LFY 1.0<br>AP1/2 1.0<br>¬CUC 1.0 | AP3 1.0<br>REV 1.0<br>SUP 1.0<br>AHP6 1.0<br>PHB 1.0<br>(+29 others) | LFY 1.0<br>AS1 1.0<br>¬STM 1.0 | ¬CUC 0.868<br>PHB 0.845<br>¬AP3 0.799 | ¬CUC<br>(10h, 40h, 132h)<br>PHB<br>(132h) |
| AP1/2 | WUS 1.0<br>AHP6 1.0<br>SEP3 1.0<br>STM 1.0<br>AP3 1.0<br>(+19 others) | WUS 1.0<br>AHP6 1.0<br>PHB 1.0<br>SEP3 1.0<br>STM 1.0<br>(+37 others) | WUS 1.0<br>AHP6 1.0<br>PHB 1.0<br>SEP3 1.0<br>STM 1.0<br>(+37 others) | STM 0.865<br>MP 0.865<br>SEP1_2 0.865<br>CUC 0.86<br>PHB 0.818<br>(+6 others) | STM 1.0<br>AP3 1.0<br>CUC 1.0<br>MP 1.0<br>SEP1_2 1.0<br>(+4 others) | AP3<br>(40h, 96h, 120h, 132h) |
| AP3 | WUS 1.0<br>AHP6 1.0<br>PHB 1.0<br>AG 1.0<br>CUC 1.0<br>(+37 others) | WUS 1.0<br>AHP6 1.0<br>PHB 1.0<br>AG 1.0<br>CUC 1.0<br>(+36 others) | WUS 1.0<br>AHP6 1.0<br>PHB 1.0<br>AG 1.0<br>CUC 1.0<br>(+36 others) | PI 1.0 | CUC 0.962<br>STM 0.962<br>¬AS1 0.962<br>PI 0.866 | STM<br>¬AS1<br>CUC<br>PI<br>(132h) |
| AS1 | WUS 1.0<br>AHP6 1.0<br>SEP3 1.0<br>CLV3 1.0<br>AG 1.0<br>(+16 others) | WUS 1.0<br>AHP6 1.0<br>PHB 1.0<br>SEP3 1.0<br>CLV3 1.0<br>(+33 others) | ¬PHB 1.0 | WUS 1.0<br>AHP6 1.0<br>PHB 1.0<br>SEP3 1.0<br>CLV3 1.0<br>(+28 others) | WUS 1.0<br>AHP6 1.0<br>PHB 1.0<br>SEP3 1.0<br>CLV3 1.0<br>(+33 others) | ¬PHB<br>(96h, 120h, 132h) |
| CLV3 | AP3 1.0<br>REV 1.0<br>SUP 1.0<br>AHP6 1.0<br>PHB 1.0<br>(+40 others) | AP3 1.0<br>REV 1.0<br>SUP 1.0<br>AHP6 1.0<br>PHB 1.0<br>(+40 others) | AP3 1.0<br>REV 1.0<br>SUP 1.0<br>AHP6 1.0<br>PHB 1.0<br>(+40 others) | AG 0.873<br>SEP3 0.873 | AG 0.812<br>SEP3 0.812<br>ETT 0.788<br>REV 0.776<br>STM 0.738 | SEP3<br>(10h, 40h, 96h, 120h, 132h) |

| | | | | | | |
|---|---|---|---|---|---|---|
| CUC1-3 | WUS 1.0<br>AHP6 1.0<br>SEP3 1.0<br>AG 1.0<br>AP1/2 1.0<br>(+9 others) | ¬AP1/2 1.0<br>¬ANT 1.0 | WUS 1.0<br>AHP6 1.0<br>SEP3 1.0<br>AG 1.0<br>AP3 1.0<br>(+11 others) | ¬ETT 0.895<br>WUS 0.808<br>SEP3 0.808<br>AG 0.808<br>AP1/2 0.808<br>(+10 others) | ¬ANT 0.988<br>AP3 0.933 | ¬ANT<br>(10h, 40h, 96h, 120h, 132h) |
| ETTIN | WUS 0.92<br>AHP6 0.92<br>PHB 0.92<br>SEP3 0.92<br>AG 0.92<br>(+34 others) | FIL 1.0<br>¬WUS 1.0<br>¬AHP6 1.0<br>¬PHB 1.0<br>¬SEP3 1.0<br>(+13 others) | WUS 1.0<br>AHP6 1.0<br>PHB 1.0<br>SEP3 1.0<br>AG 1.0<br>(+36 others) | SEP3 1.0<br>AG 1.0<br>¬CUC 0.987 | PHB 0.988<br>ANT 0.969 | PHB<br>ANT<br>(132h) |
| FIL | STM 1.0<br>AHP6 1.0<br>PHB 1.0<br>SEP3 1.0<br>CUC 1.0<br>(+37 others) | STM 1.0<br>AHP6 1.0<br>PHB 1.0<br>SEP3 1.0<br>CUC 1.0<br>(+37 others) | AS1 1.0 | STM 1.0<br>AHP6 1.0<br>PHB 1.0<br>SEP3 1.0<br>CUC 1.0<br>(+37 others) | AP1/2 1.0<br>AS1 1.0<br>LFY 1.0<br>¬STM 1.0<br>¬PHB 1.0<br>(+27 others) | AS1<br>(10h, 40h, 96h, 120h, 132h) |
| LFY | STM 1.0<br>AHP6 1.0<br>PHB 1.0<br>SEP3 1.0<br>AG 1.0<br>(+28 others) | STM 1.0<br>AHP6 1.0<br>PHB 1.0<br>SEP3 1.0<br>AG 1.0<br>(+28 others) | STM 1.0<br>AHP6 1.0<br>PHB 1.0<br>SEP3 1.0<br>AG 1.0<br>(+29 others) | STM 1.0<br>AHP6 1.0<br>PHB 1.0<br>SEP3 1.0<br>AG 1.0<br>(+26 others) | STM 1.0<br>AHP6 1.0<br>PHB 1.0<br>SEP3 1.0<br>AG 1.0<br>(+25 others) | ¬STM<br>¬SEP3<br>AS1<br>(120h, 132h) |
| PHB/PHV | REV 1.0<br>¬FIL 1.0<br>PUCHI 0.991 | STM 1.0<br>REV 1.0<br>¬ETT 1.0<br>¬FIL 1.0 | REV 1.0 | REV 0.619<br>¬ETT 0.619<br>¬FIL 0.619 | ¬FIL 0.985<br>MP 0.948<br>REV 0.896 | REV<br>(10h, 40h, 96h, 120h, 132h) |
| PI | WUS 1.0<br>AHP6 1.0<br>PHB 1.0<br>AG 1.0<br>CUC 1.0<br>(+37 others) | WUS 1.0<br>AHP6 1.0<br>PHB 1.0<br>AG 1.0<br>CUC 1.0<br>(+36 others) | WUS 1.0<br>AHP6 1.0<br>PHB 1.0<br>AG 1.0<br>CUC 1.0<br>(+36 others) | AP3 1.0 | AP3 0.934<br>CUC 0.901<br>STM 0.901<br>¬AS1 0.901<br>WUS 0.873<br>(+5 others) | AP3<br>(10h, 40h, 96h, 120h, 132h) |
| REV | PHB 0.893<br>LFY 0.893<br>PUCHI 0.893<br>ANT 0.893 | PHB 0.773<br>LFY 0.773<br>AP1/2 0.773<br>ANT 0.773 | PHB 0.838 | PHB 0.619<br>¬ETT 0.619<br>¬FIL 0.619 | PHB 0.814<br>¬FIL 0.814<br>¬ETT 0.802<br>MP 0.784<br>SUP 0.782<br>(+1 others) | ¬ETT<br>PHB<br>(120h, 132h) |
| STM | WUS 1.0<br>CLV3 1.0<br>SEP3 1.0<br>AG 1.0<br>AP1/2 1.0 | PHB 1.0<br>REV 1.0 | PHB 0.748<br>REV 0.748<br>AP1/2 0.679<br>ANT 0.679<br>MP 0.679 | SEP3 0.99<br>AG 0.99 | SEP3 1.0<br>AG 1.0 | AG<br>(120h, 132h) |

| | | | | | | |
|---|---|---|---|---|---|---|
| | (+27 others) | | (+3 others) | | | |
| SUP | WUS 1.0<br>AHP6 1.0<br>PHB 1.0<br>CUC 1.0<br>CLV3 1.0<br>(+38 others) | WUS 1.0<br>AHP6 1.0<br>PHB 1.0<br>CUC 1.0<br>CLV3 1.0<br>(+38 others) | WUS 1.0<br>AHP6 1.0<br>PHB 1.0<br>CUC 1.0<br>CLV3 1.0<br>(+38 others) | WUS 1.0<br>AHP6 1.0<br>CUC 1.0<br>CLV3 1.0<br>AS1 1.0<br>(+37 others) | REV 0.972<br>ANT 0.926<br>PHB 0.917<br>ETT 0.917<br>¬CUC 0.917 | ANT<br>(10h, 40h, 96h, 120h, 132h) |

**Table S4:** Summary of novel inputs and their effect for every gene at different time points. 'Summ' column indicates gene interactions with most significant impact throughout development (stages at which they occur indicated between brackets as time points). The notation is the same as in Table S3. Note that when the pattern is good already for the literature-based regulatory inputs (see table 2), many possible hypotheses are regarded as good by the algorithm as long as they do not degrade the pattern. They are mentioned here for completeness.

**Note that these hypotheses come in addition to those in table 3. For example, SUP requires at least AP3/PI or AG to be switched on. If this is not the case (as in the first three timepoints), any other gene will not be able to switch on SUP.**

### AGAMOUS

| | Stages with approx. zones of future meristem (green) and sepals (yellow) marked | AGAMOUS top view | AGAMOUS section | Rozier et al | Wolmann et al | Urbanus et al | Unpublished results |
|---|---|---|---|---|---|---|---|
| Initium stage | 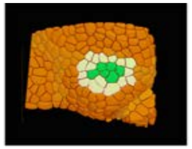 | 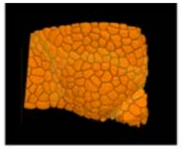 | 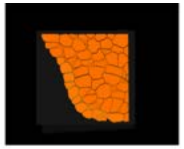 | 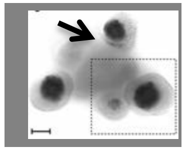 | | | 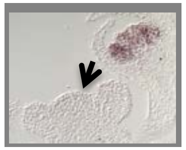 |
| stage 1 | 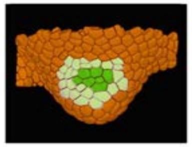 | 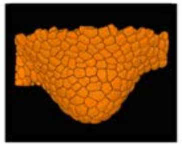 | 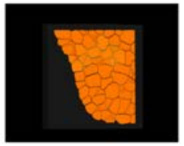 | 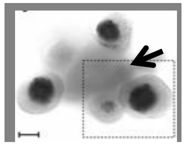 | | | 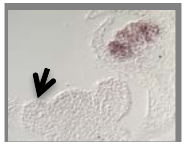 |
| stage 2 | 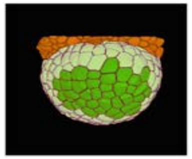 | 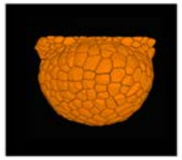 | 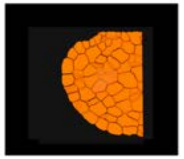 | 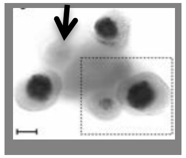 | | | 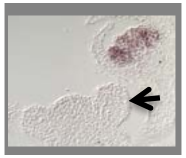 |
| stage 3 | 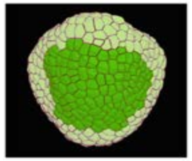 | 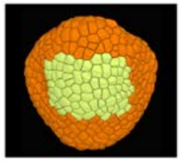 | 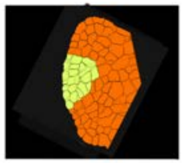 | 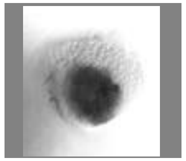 | 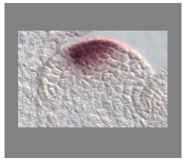 | | 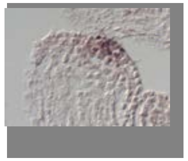 |
| stage 4 | 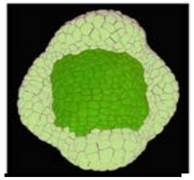 | 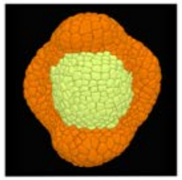 | 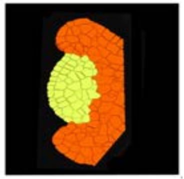 | 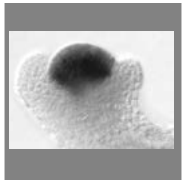 | 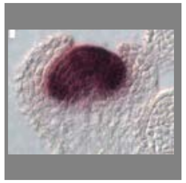 | 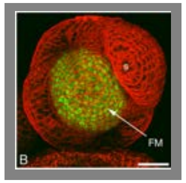 | 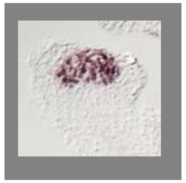 |
| Remarks: | - AGAMOUS is expressed from stage 2 onwards, in a zone that extends to one or two cells from the sepal boundary (to leave space for the A function genes in the second whorl) | | | | | | |

### AHP6

| | Stages of flower development with future meristem and sepals marked | topview | section | Rozier et al | Besnard et al. | Cellier and Traas unpublished | Das et al unpublished |
|---|---|---|---|---|---|---|---|
| Initium stage | 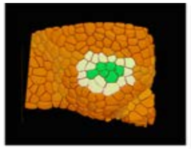 | 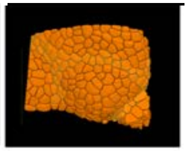 | 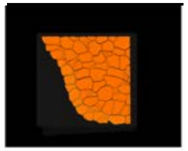 | 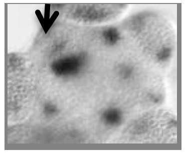 | 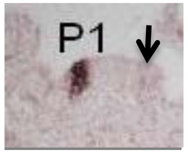  | | |
| stage 1 | 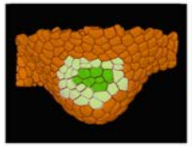 | 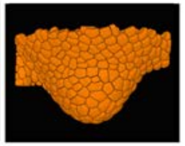 | 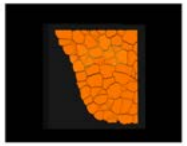 | 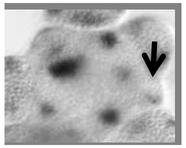 | 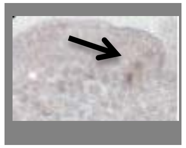 | | |
| stage 2 | 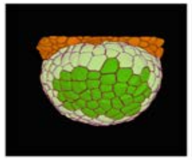 | 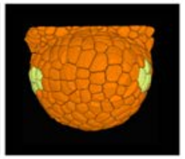 | 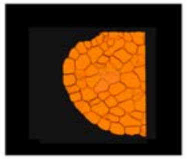 | 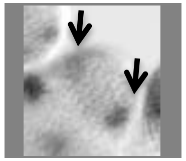 | 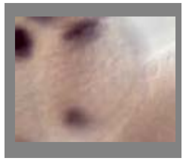 | | 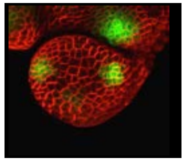 |
| stage 3 | 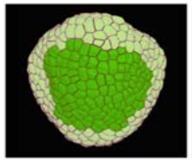 | 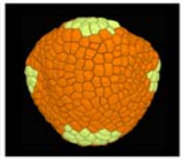 | 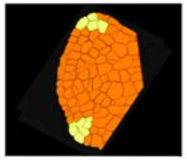 | 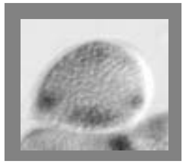 | 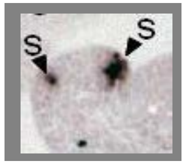  | | 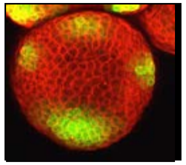 |
| stage 4 | 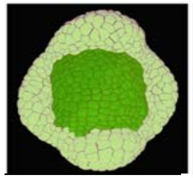 | 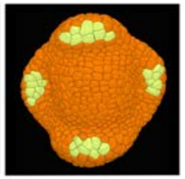 | 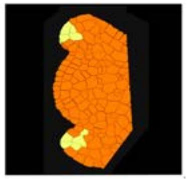 | | | 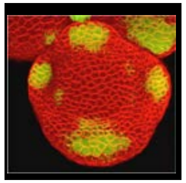 | |
| Remarks: | There is some ambiguity regarding the early stages.In situs in Besnard et al suggest that AHP6 is switched very early during organ formation. At what stage precisely is impossible to determine. This also seems to vary between individual meristems. pAHP6:GFP transcriptional fusion line as described in Besnard et al | | | | | | |

### AINTEGUMENTA

| | Stages of flower development with future meristem and sepals marked | AINTEGUMENTA topview | AINTEGUMENTA section | Elliott et al 1996 | Yamaguchi et al 2012 | Krizek 2009 | unpublished (in situ hybridization and GFP (for ref on GFP line see Grandjean et al |
|---|---|---|---|---|---|---|---|
| Initium stage | 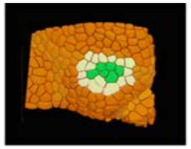 | 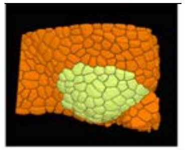 | 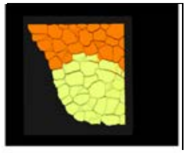 | | 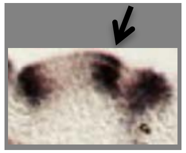 | | 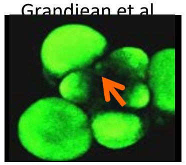 |
| stage 1 | 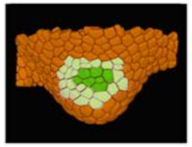 | 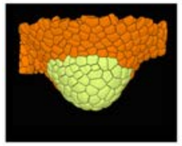 | 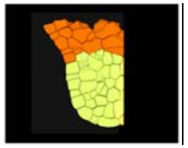 | 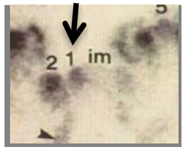 | 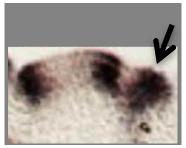 | | 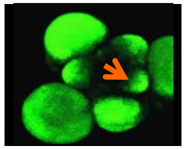 |
| stage 2 | 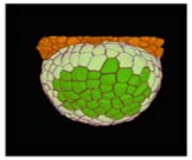 | 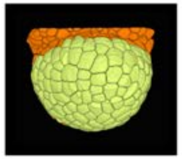 | 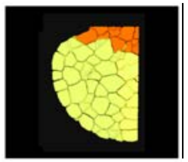 | 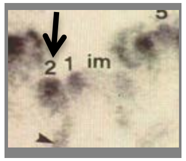 | | | 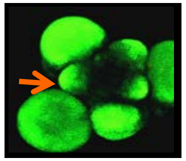 |
| stage 3 | 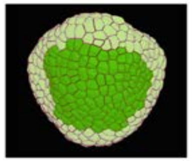 | 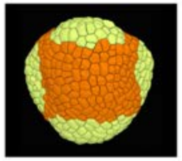 | 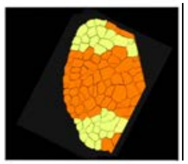 | | 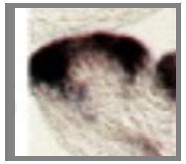 | 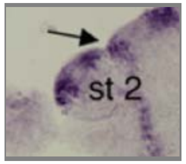 | 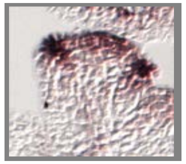 |
| stage 4 | 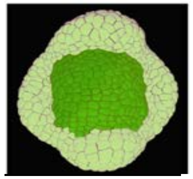 | 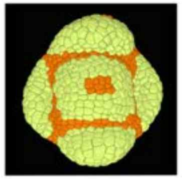 | 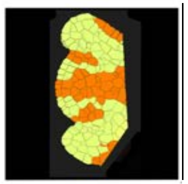 | 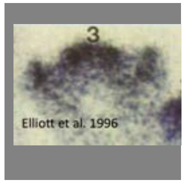 | | | 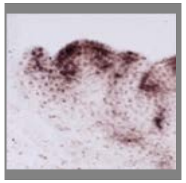 |
| Remarks: | No clear image found of stage 2, but ANT seems to be expressed in the bract | | | | | | |

### APETALA1

| | Stages of flower development with future meristem and sepals marked | APETALA 1top view | APETALA1 section | Yu et al (supp figure) | urbanus et al | unpublished | unpublished |
|---|---|---|---|---|---|---|---|
| Initium stage | 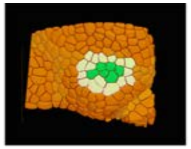 | 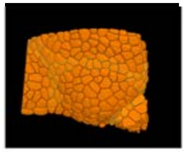 | 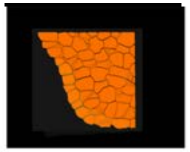 | | 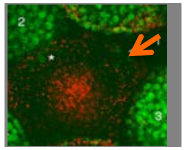 | | |
| stage 1 | 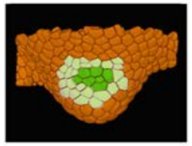 | 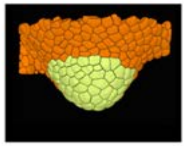 | 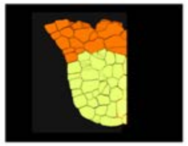 | | 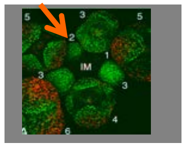 | | 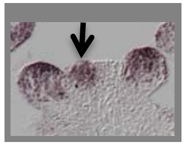 |
| stage 2 | 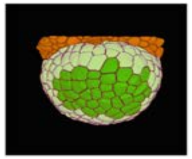 | 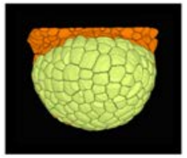 | 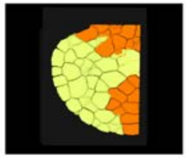 | 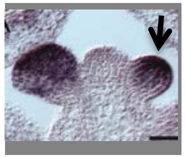 | 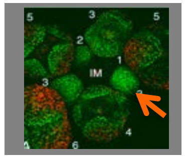 | | 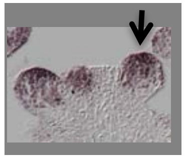 |
| stage 3 | 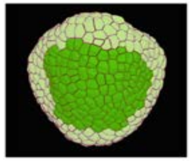 | 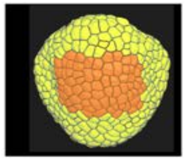 | 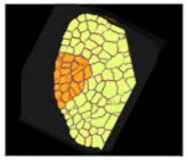 | 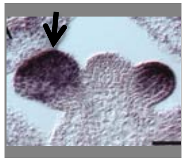 | 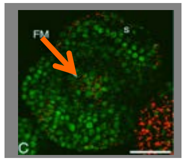 | 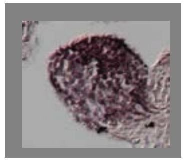 | 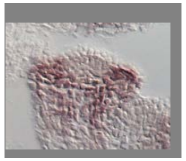 |
| stage 4 | 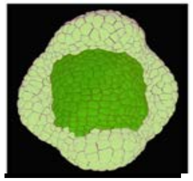 | 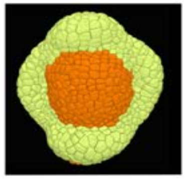 | 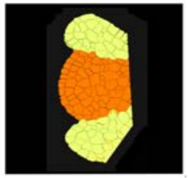 | | 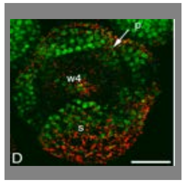 | | 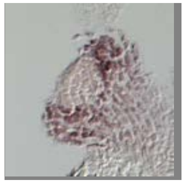 |
| Remarks: | AP1 is weakly expressed during the very early stages (urbanus et al), when the sepals start to grow out (stage 3) expression becomes rapidly weaker in the centre. Expression in the sepals is slightly stronger adaxially. Note also expression in the future petal region in late stage 3. Contrasting results when it comes to maintenance of expression of throughout stage 3, which likely the stage where AP1 is inactivated. | | | | | | |

### APETALA2

| | Stages of flower development with future meristem and sepals marked | APETALA2 top view | APETALA2 section | Wollmann et al 2010 | Jofoku et al 1994 | | unpublished data |
|---|---|---|---|---|---|---|---|
| Initium stage | 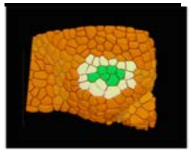 | 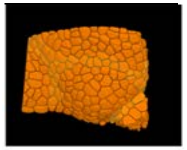 | 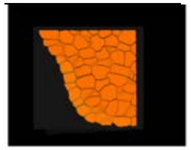 | | | | |
| stage 1 | 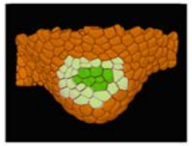 | 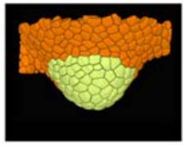 | 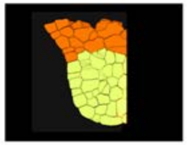 | 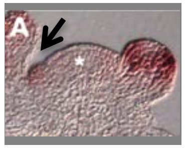 | | | 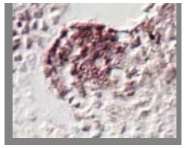 |
| stage 2 | 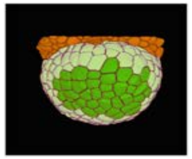 | 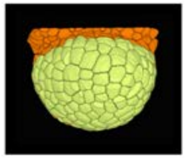 | 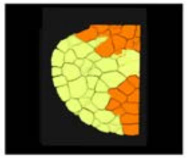 | | 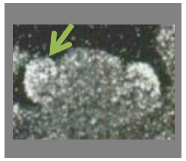 | | 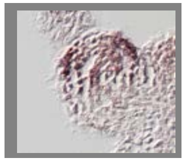 |
| stage 3 | 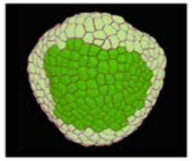 | 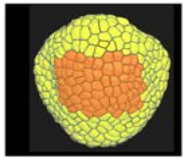 | 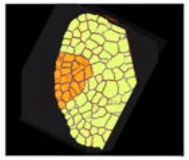 | 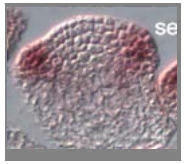 | | | 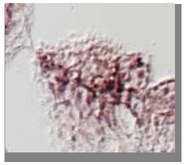 |
| stage 4 | 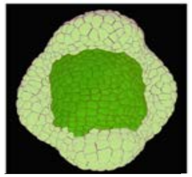 | 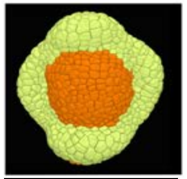 | 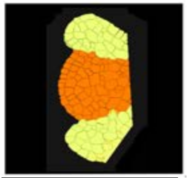 | 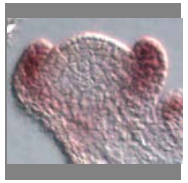 | 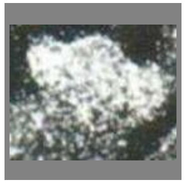 | | |
| Remarks: | Very few articles show the precise expression patterns of AP2. The patterns published by Jofuku et al seem in contradiction with those published by Wollmann et al. when it comes to late stage 3. We note, however, that Jofuku et al do not publish consistent patterns (one figure showing labelling in the inflorescence meristem, the other not). In addition, our results confirm a pattern which is very close or identical to the one published by Wollmann et al. Together, the data indicate a pattern very close to that of AP1. | | | | | | |

### APETALA3

| | Stages of flower development with future meristem and sepals marked / top view / section / Yu et al 2004 / Goto et al 1994 / Prunet et al unpublished / unpublished |
|---|---|
| Initium stage | 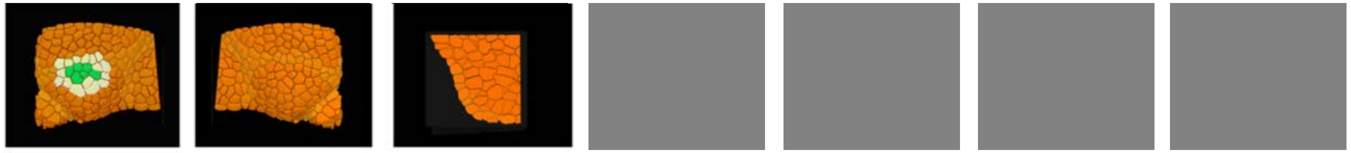 |
| stage 1 | 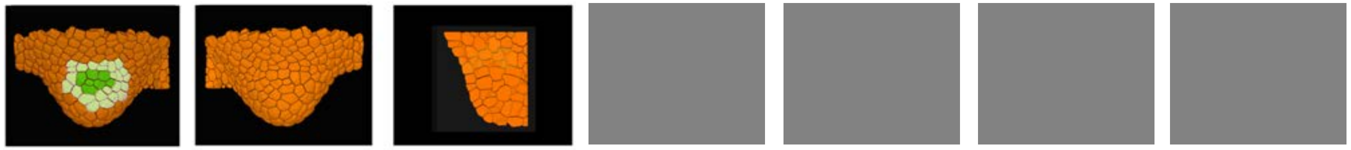 |
| stage 2 | 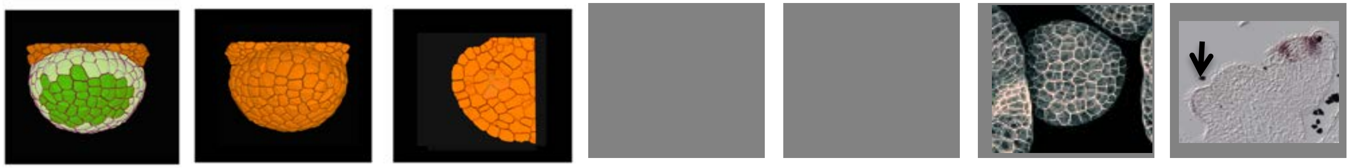 |
| stage 3 | 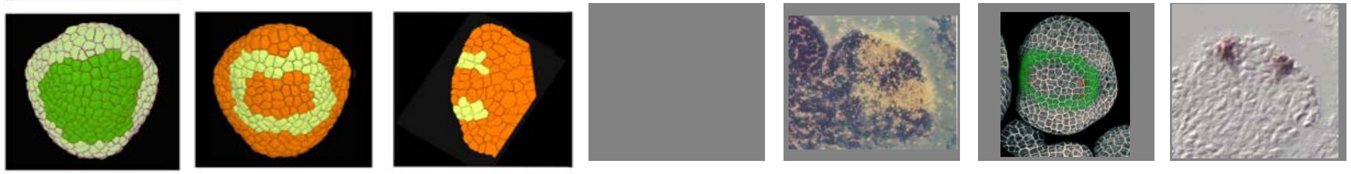 |
| stage 4 | 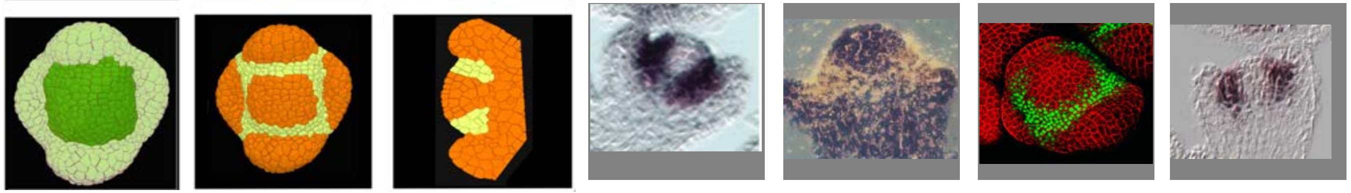 |

Remarks: Note that on some images of the late stage 3, the domain of AP3 seems to cover a broader domain than recorded in the Atlas. This might be due to the fact that the sections were not median . We have therefore chosen to base the interpretation on the 3D pattern visualized by the GFP line, covering a domain of about 4 cells wide at the boundary between sepal and meristem.

### ASYMMETRIC LEAVES 1

| | Stages of flower development with future meristem and sepals marked | top view | section | Byrne et al | unpublished | References |
|---|---|---|---|---|---|---|
| Initium stage | | | | | | 1. Byrne ME, Barley R, Curtis M, Arroyo JM, Dunham M, Hudson A, et al. Asymmetric leaves1 mediates leaf patterning and stem cell function in Arabidopsis. Nature. 2000;408(6815):967-71 |
| stage 1 | | | | | | |
| stage 2 | | | | | | |
| stage 3 | | | | | | |
| stage 4 | | | | | | |

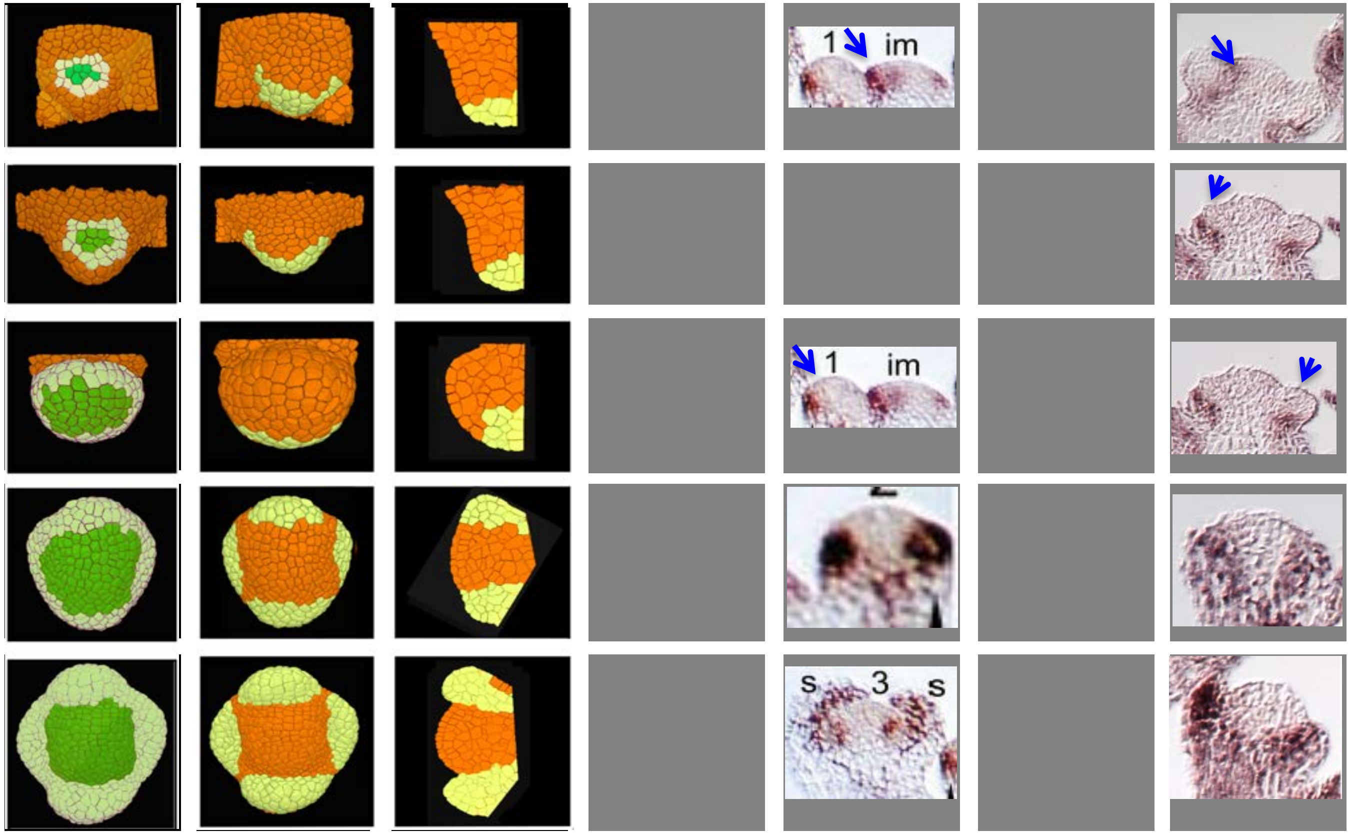

Remarks: AS1 is first expressed in the cryptic bract, than in the sepal primordia. Weaker on abaxial side from late stage 3 onwards

### ATML1

| | Stages of flower development with future meristem and sepals marked | top view | section | Lu et al | Sessions et al | Unpublished |
|---|---|---|---|---|---|---|
| Initium stage | 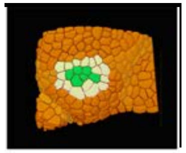 | 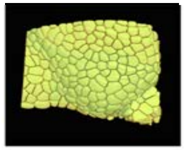 | 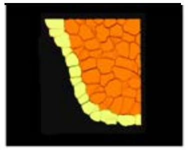 | 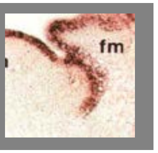 | | |
| stage 1 | 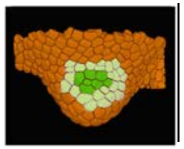 | 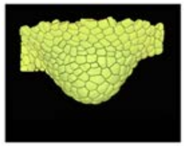 | 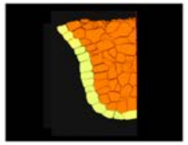 | | 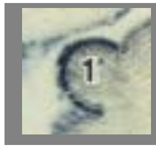 | |
| stage 2 | 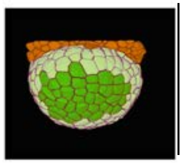 | 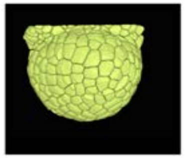 | 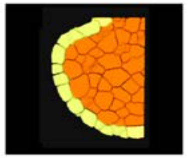 | | 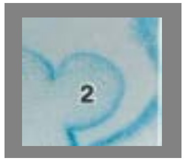 | 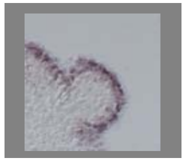 |
| stage 3 | 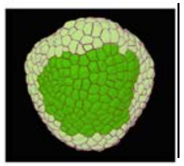 | 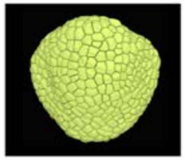 | 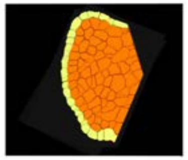 | | 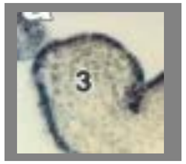 | |
| stage 4 | 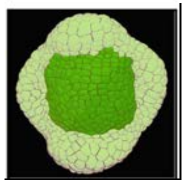 | 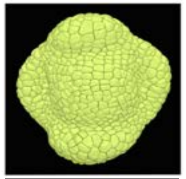 | 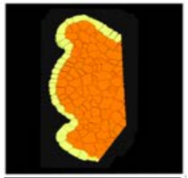 | | 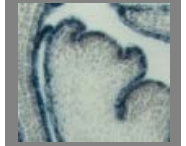 | 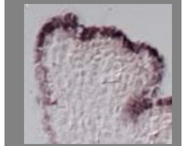 |
| Remarks: | | | | | | |

### CLAVATA3

| | Stages of flower development with future meristem and sepals marked | top view | section | Maier et al 2011 | Zhang et al 2011 | Lenhard and Laux 2003 | Rozier et al 2014 |
|---|---|---|---|---|---|---|---|
| Initium stage | | | | | | | |
| stage 1 | | | | | | | |
| stage 2 | | | | | | | |
| stage 3 | | | | | | | |
| stage 4 | | | | | | | |

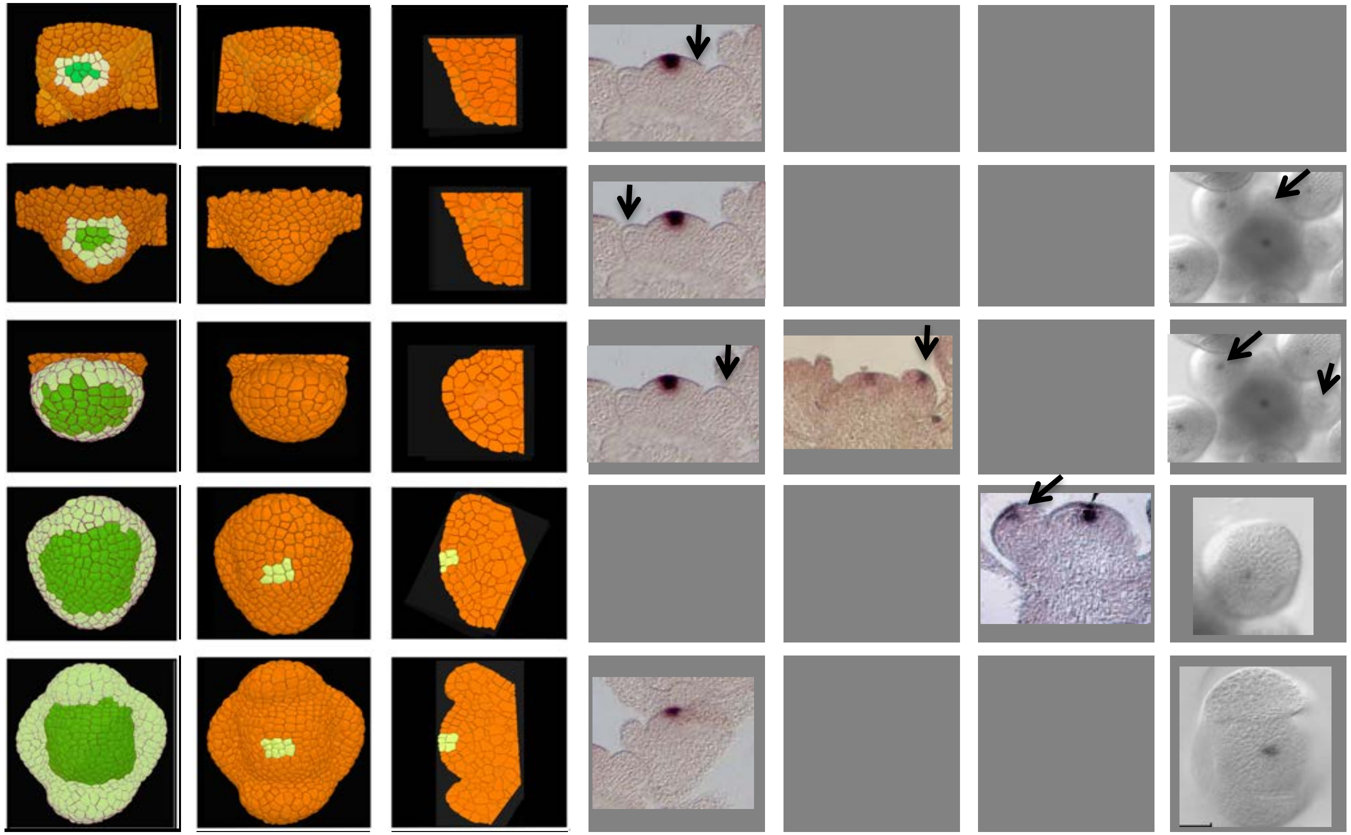

Remarks: There is some ambiguity as to when CLV3 is activated during stage 2. The absence of label in the image coming from Maier et al might be due to the plane of section. On the other hand the positive labeling at stage 2 described Zhang et al could also be due to the plane of section, and in fact represent and older stage. The whole mount described by Rozier et al suggest at most a weak expression at this stage.

### CUC 1, 2 3

| | Stages of flower development with future meristem and sepals marked | top view | section | Vachez et al unpublished (CUC1) | Vachez et al unpublished (CUC1) | Yamaguchi et al 2014 (CUC2) | Hibara et al 2006 (CUC3) | References |
|---|---|---|---|---|---|---|---|---|
| Initium stage | 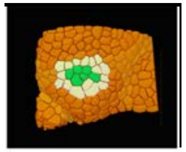 | 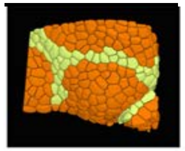 | 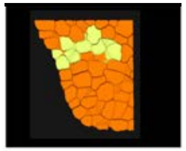 | | | | 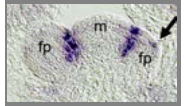  | |
| stage 1 | 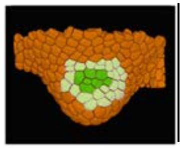 | 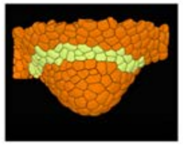 | 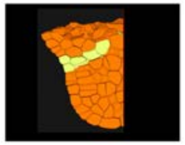 | | | | | |
| stage 2 | 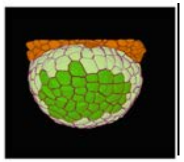 | 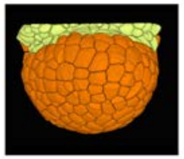 | 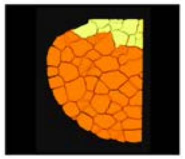 | | | | 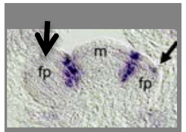  | |
| stage 3 | 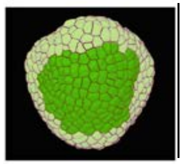 | 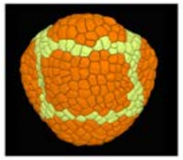 | 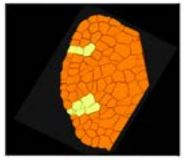 | 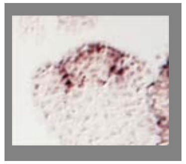 | 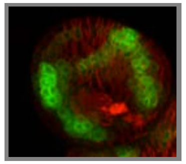 | 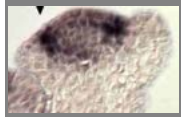 | | |
| stage 4 | 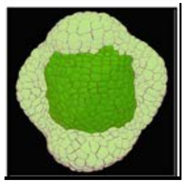 | 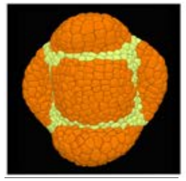 | 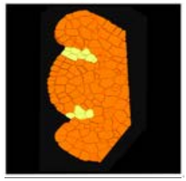 | | | 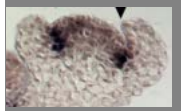 | 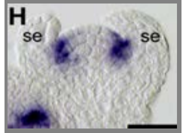  | |

Remarks: Very few studies provide expression data. The avalaible data indicate, that CUCs have a similar or identical expression pattern. In the very young priordium, they could potentially be expressed on the abaxial side of the flower bud (Hibara et al), but we could not confirm this. Lateron, the two studies and our own results indicate that the CUC genes are expressed at the periphery of the floral meristem, at the boundary. They are not detected in the adaxial boundary region of the sepals.

### ETTIN/ARF3

| | Stages of flower development with future meristem and sepals marked | topview | section | Pekker et al 2005 | Liu et al | Rozier et al unpublished |
|---|---|---|---|---|---|---|
| Initium stage | 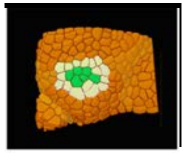 | 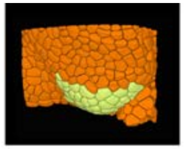 | 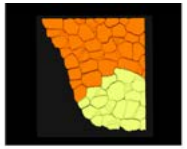 | 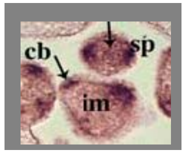  | 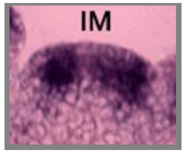  | 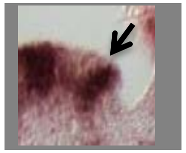 |
| stage 1 | 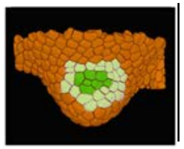 | 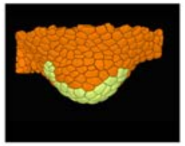 | 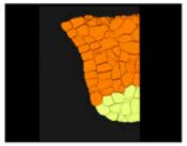 | 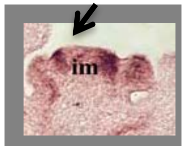  | | 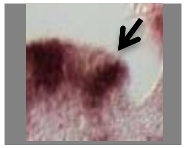 |
| stage 2 | 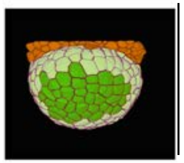 | 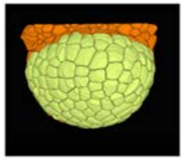 | 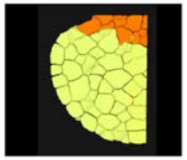 | 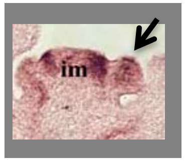  | 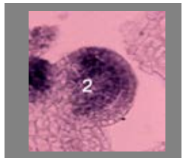  | 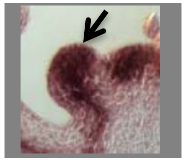 |
| stage 3 | 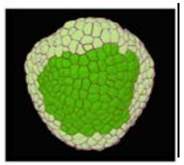 | 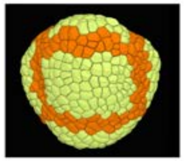 | 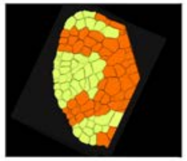 | 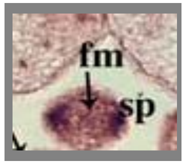  | 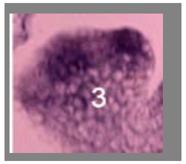  | 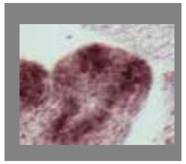 |
| stage 4 | 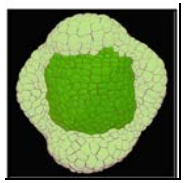 | 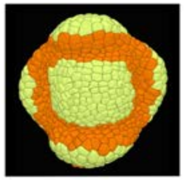 | 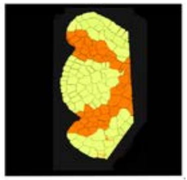 | | | 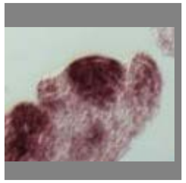 |
| Remarks: | There seemed to be a contradiction between the patterns published by Pekker et al (2005) and Liu et al (2014). This might be due to the plane of section. We therefore performed in situ hybridizations and examined serial sections. This confirmed the observations of Pekker et al that ETT/ARF3 is strongly expressed in the crypric bract (cb in the images form Pekker et al, arrow in Rozier et al.). Our data also confirmed the results by Liu et al, that ETT is strongly expressed in the floral meristem during stage 2 and 3. All results confrim the expression of ETT_ARF3 in the sepal primordia (sp in images from | | | | | |

### FILAMENTOUS FLOWER

| | Stages of flower development with future meristem and sepals marked | topview | section | Eshed et al 2001 | Sawa et al 1999 (line a and d), Dinneny et al | Heilser et al 2005 (a,b), Zhao et al unpublished (d,e) | Vachez et al unpublished |
|---|---|---|---|---|---|---|---|
| Initium stage | 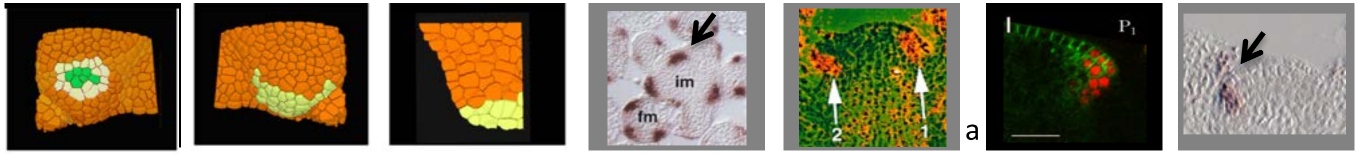 | | | | | | |
| stage 1 | 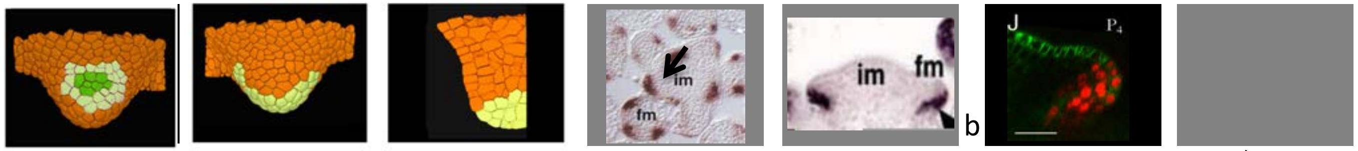 | | | | | | |
| stage 2 | 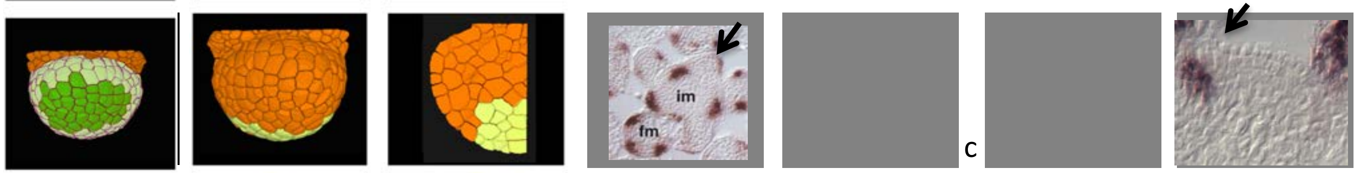 | | | | | | |
| stage 3 | 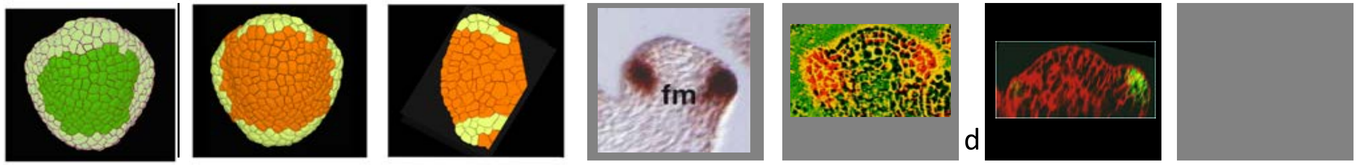 | | | | | | |
| stage 4 | 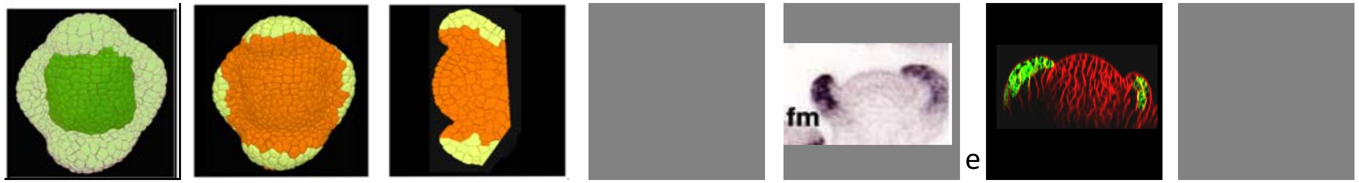 | | | | | | |
| Remarks: | Labelling of the cryptic bract also illustrated by Dinneny et al. | | | | | | |

### LEAFY

| | Stages of flower development with future meristem and sepals marked | topview | section | Blazquez et al. 1997 | Vernoux et al 2000 | Karim et al 2009 | Vachez et al unpublished |
|---|---|---|---|---|---|---|---|
| Initium stage | 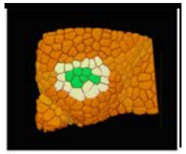 | 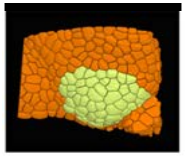 | 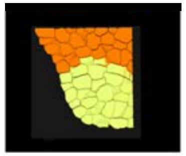 | 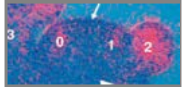 | | | 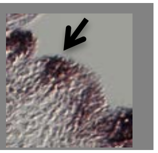 |
| stage 1 | 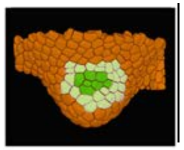 | 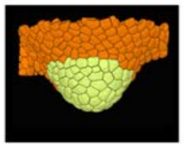 | 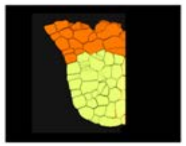 | 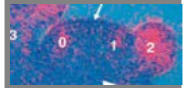 | | | 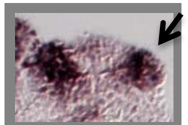 |
| stage 2 | 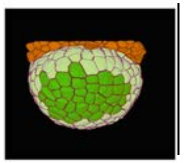 | 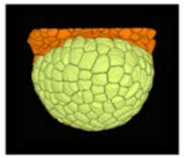 | 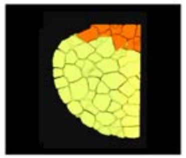 | 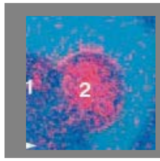 | | 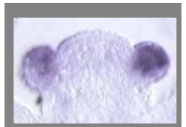 | 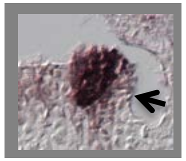 |
| stage 3 | 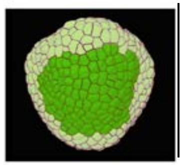 | 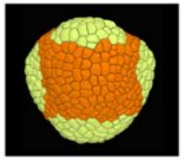 | 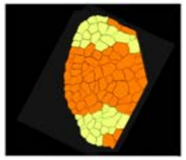 | | 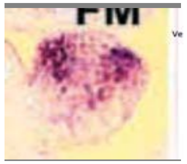 | | 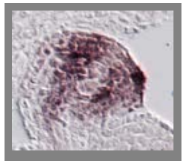 |
| stage 4 | 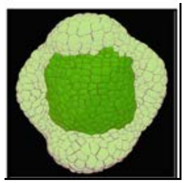 | 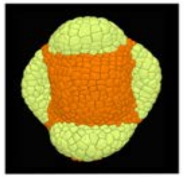 | 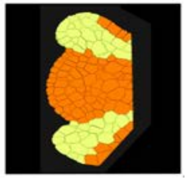 | 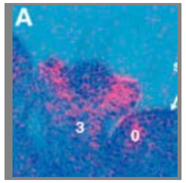 | | | 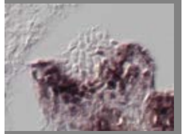 |
| Remarks: | LEAFY is expressed from the earliest stages onwards. It is not clear if the gene is expressed in the cryptic bract (Karim et al, our own results: see arrows stage 2). In most early stage 3 meristems low expression is observed in the central part . We observed, however in a few of these meristems that LFY was still high throughout the flower. We conclude that LFY becomes rapidly downregulated in the floral meristem proper during early stage 3. Note that this seems in contradiction with certain GFP lines (e.g. Chandler and Werr 2017, fig 7. | | | | | | |

### MONOPTEROS_ARF5

<table>
<tr><th></th><th>Stages of flower development with future meristem and sepals marked</th><th>topview</th><th>section</th><th>Yamaguchi et al 2012</th><th>Hardtke and Berleth, 1998</th><th>Vachez et al unpublished</th><th>Vachez et al unpublished</th><th>References</th></tr>
<tr><td>Initium stage</td><td colspan="7">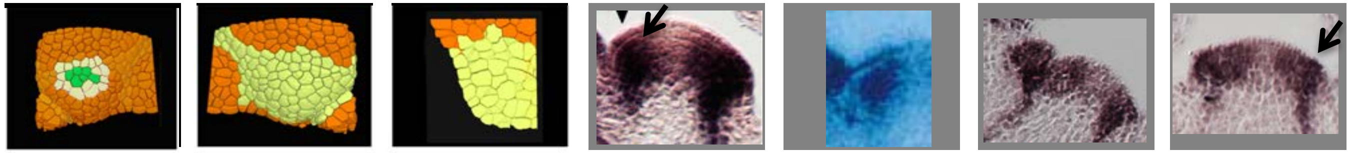</td><td rowspan="5"></td></tr>
<tr><td>stage 1</td><td colspan="7">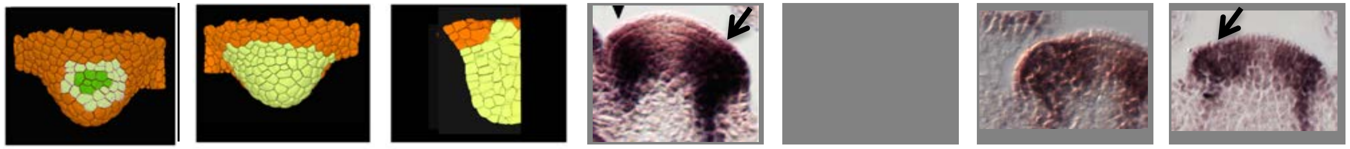</td></tr>
<tr><td>stage 2</td><td colspan="7">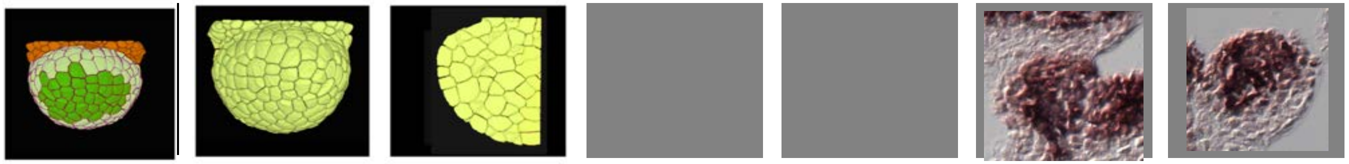</td></tr>
<tr><td>stage 3</td><td colspan="7">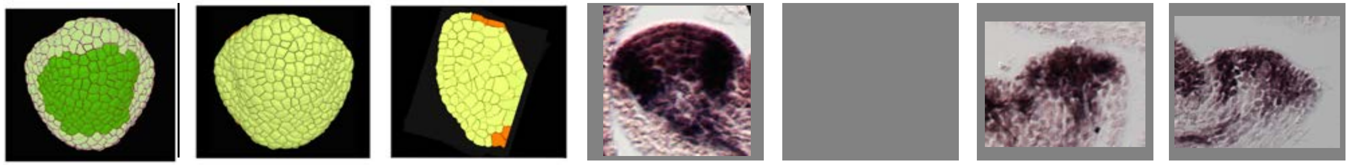</td></tr>
<tr><td>stage 4</td><td colspan="7">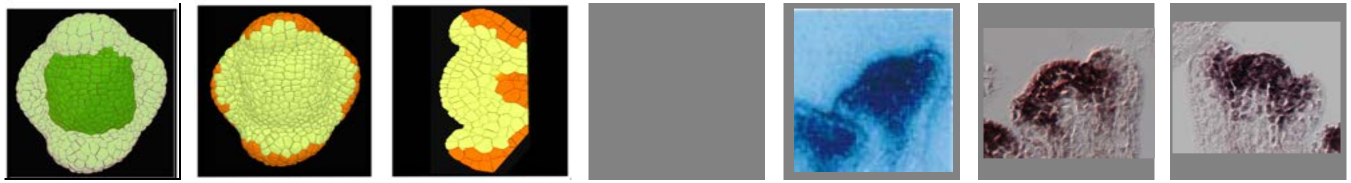</td></tr>
<tr><td>Remarks:</td><td colspan="7">Results from Hardtke and Berleth suggest that MP is not expressed at the meristem surface. Results from Yamaguchi et al as well our own data indicate that MP is expressed at the surface, albeit at somewhat lower levels. We also note a clear expression at the adaxial side of the sepal primordia from stage 3 onwards. There remains some incertainty regarding the level of expression in organ boundaries, where expression is lower in some samples.</td><td></td></tr>
</table>

1. Hardtke CS, Berleth T. The Arabidopsis gene MONOPTEROS encodes a transcription factor mediating embryo axis formation and vascular development. EMBO J. 1998;17(5):1405-11.
2. Vernoux T, Brunoud G, Farcot E, Morin V, Van den Daele H, Legrand J, et al. The auxin signalling network translates dynamic input into robust patterning at the shoot apex. Mol Syst Biol. 2011;7:508.
3. Yamaguchi N, Wu MF, Winter CM, Berns MC, Nole-Wilson S, Yamaguchi A, et al. A molecular framework for auxin-mediated initiation of flower primordia. Dev Cell. 2013;24(3):271-82.

### PHABULOSA

| | Stages of flower development with future meristem and sepals marked | topview | section | Nole-Wilson and Krizek 2006 | | Vachez et al unpublished |
|---|---|---|---|---|---|---|
| Initium stage | 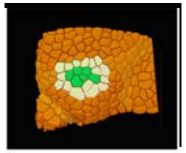 | 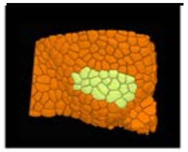 | 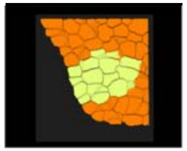 | 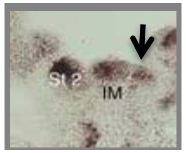 | | 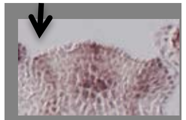 |
| stage 1 | 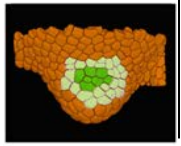 | 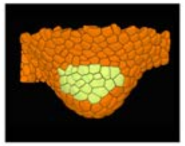 | 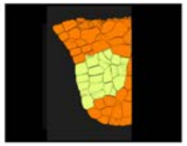 | | | 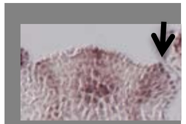 |
| stage 2 | 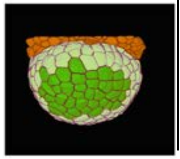 | 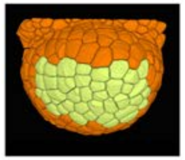 | 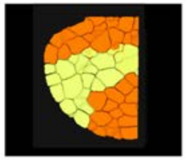 | 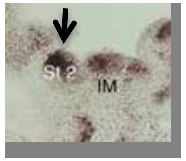 | | 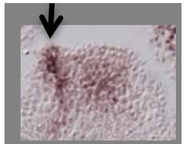 |
| stage 3 | 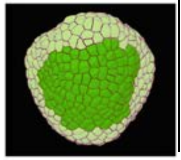 | 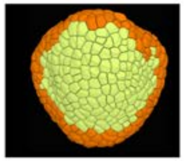 | 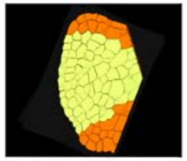 | | | 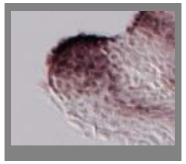 |
| stage 4 | 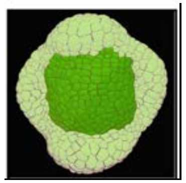 | 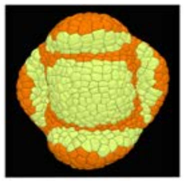 | 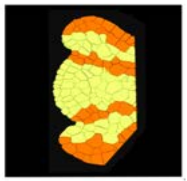 | 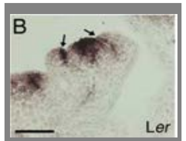 | | 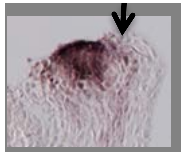 |
| Remarks: | According to Emery et al (2003) very similar to PHV and REV. Note that the adaxial expression in sepals and young flower primordia is weaker. PHB is systematically strong in floral meristems. | | | | | |

### PHAVOLUTA

| | Stages of flower development with future meristem and sepals marked | topview | section | Emery et al 2003 | | Vachez et al unpublished |
|---|---|---|---|---|---|---|
| Initium stage | 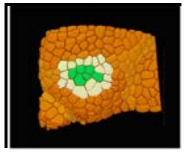 | 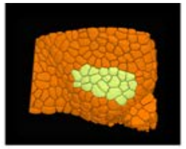 | 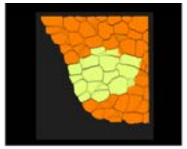 | | | |
| stage 1 | 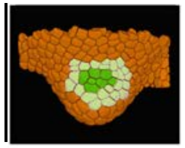 | 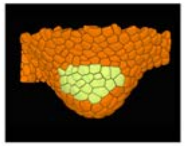 | 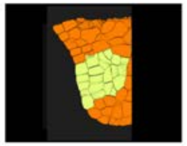 | | | |
| stage 2 | 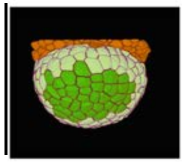 | 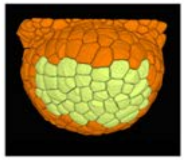 | 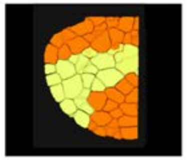 | 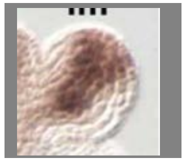 | | 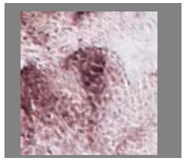 |
| stage 3 | 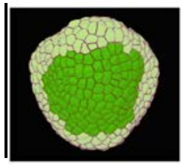 | 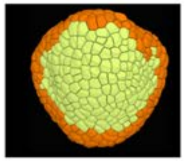 | 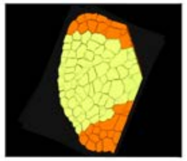 | | | 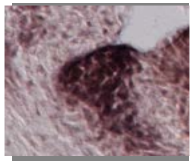 |
| stage 4 | 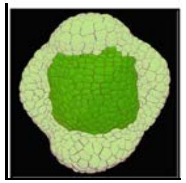 | 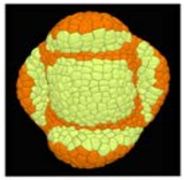 | 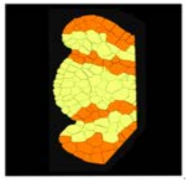 | | | 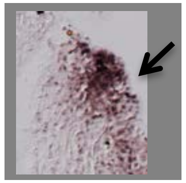 |
| Remarks: | There are little data related to PHV expression. According to Emery et al: the class III HDSummary ZIP genes REVOLUTA (REV), PHABULOSA (PHB), and PHAVOLUTA (PHV) exhibit similar mRNA expression in apical and floral meristems Note that small area without PHV expression in the boundary between sepal primordia and floral meristem (as PHB (Nole-Wilson and Krizek (2006), REV Otsuga et a 2001l). | | | | | |

### PISTILLATA

| | Stages of flower development with future meristem and sepals marked | topview | section | Goto and Meyerowtiz 1994 | Prunet and Meyerowitz unpublished | | Vachez et al unpublished |
|---|---|---|---|---|---|---|---|
| Initium stage | 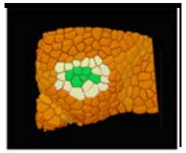 | 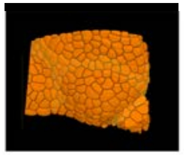 | 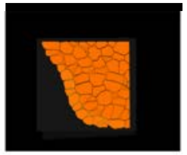 | | | | |
| stage 1 | 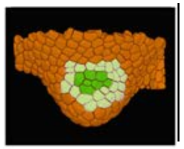 | 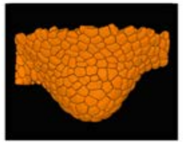 | 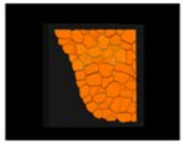 | | | | 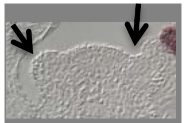 |
| stage 2 | 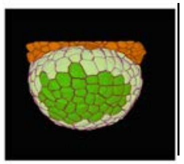 | 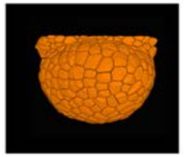 | 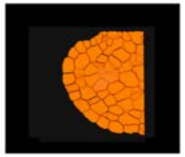 | | 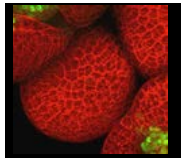 | | 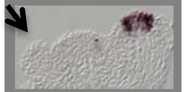 |
| stage 3 | 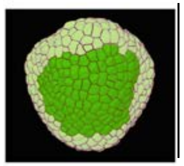 | 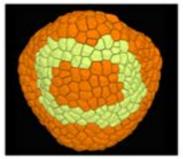 | 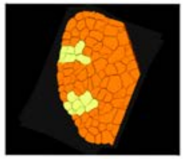 | 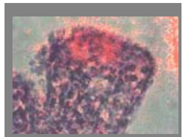 | | | 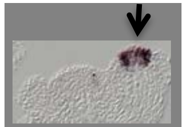 |
| stage 4 | 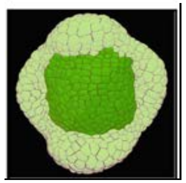 | 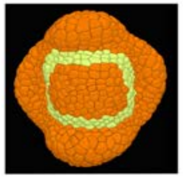 | 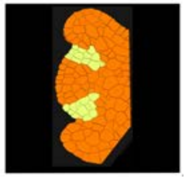 | 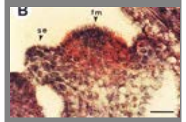 | 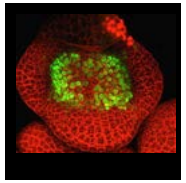 | | 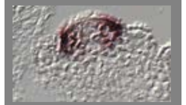 |
| Remarks: | Very little references with reliable expression patterns found. | | | | | | |

### PUCHI

| | Stages of flower development with future meristem and sepals marked | topview | section | Karim et al 2009 | Karim et al 2009 |
|---|---|---|---|---|---|
| Initium stage | 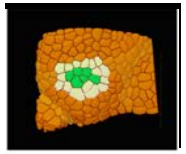 | 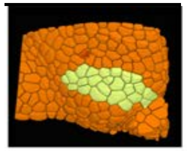 | 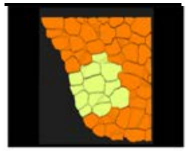 | 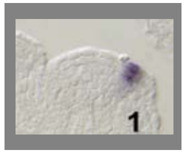 | 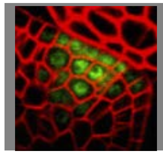 |
| stage 1 | 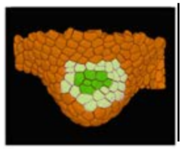 | 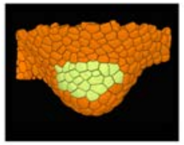 | 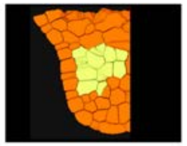 | 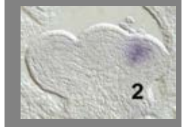 | |
| stage 2 | 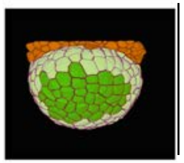 | 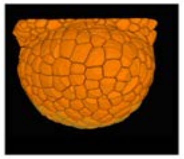 | 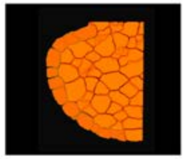 | 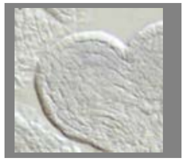 | |
| stage 3 | 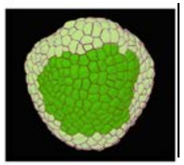 | 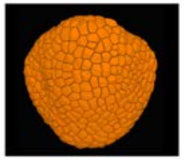 | 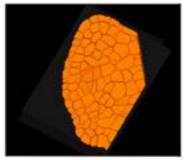 | 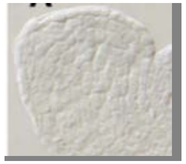 | |
| stage 4 | 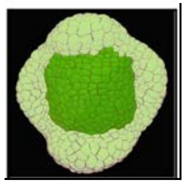 | 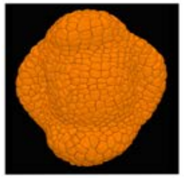 | 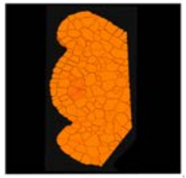 | | |
| Remarks: | | | | | |

### REVOLUTA

| | Stages of flower development with future meristem and sepals marked | topview | section | Heisler et al 2005 | Otsuga et al 2001 | Emery et al 2003 | Vachez et al unpublished |
|---|---|---|---|---|---|---|---|
| Initium stage | 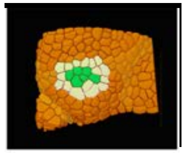 | 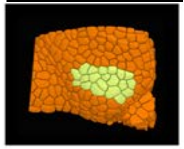 | 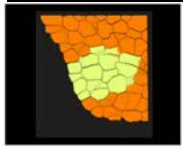 | 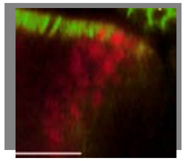 | | | |
| stage 1 | 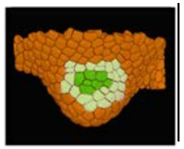 | 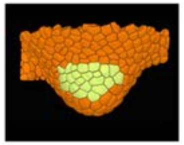 | 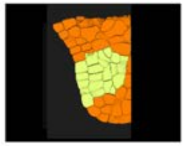 | 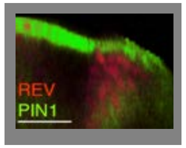 | | | 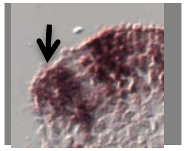 |
| stage 2 | 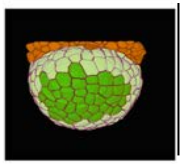 | 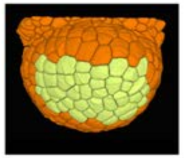 | 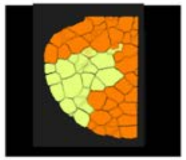 | | 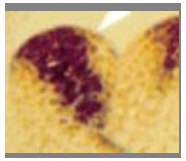 | 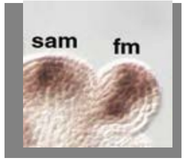 | |
| stage 3 | 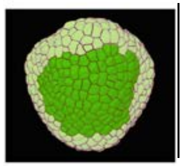 | 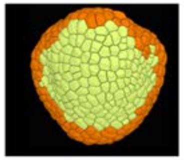 | 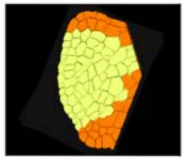 | | | | |
| stage 4 | 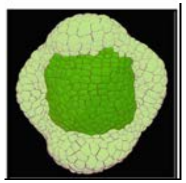 | 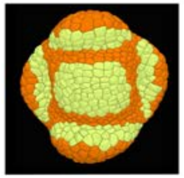 | 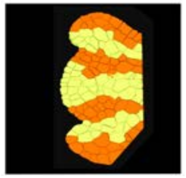 | | 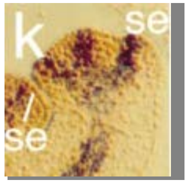 | | 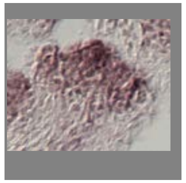 |
| Remarks: | All results agree on the patterns until late stage 3. The image of Otsuga suggests expression at a narrow zone at the floral meristem and the abaxial side of t | | | | | | |

### SEPALLATA1/AGL2

| | Stages of flower development with future meristem and sepals marked | top view | section | Flanagan and Ma, 1994 | Flanagan and Ma, 1994 (bright field of previous column | | | References |
|---|---|---|---|---|---|---|---|---|
| Initium stage | 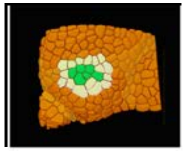 | 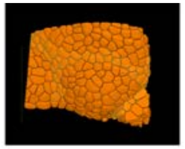 | 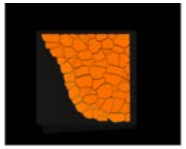 | 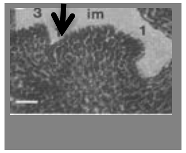 | 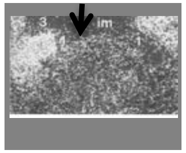 | | | 1. Flanagan CA, Ma H. Spatially and temporally regulated expression of the MADS-box gene AGL2 in wild-type and mutant arabidopsis flowers. Plant Mol Biol. 1994;26(2):581-95.<br>2. Pelaz S, Ditta GS, Baumann E, Wisman E, Yanofsky MF. B and C floral organ identity functions require SEPALLATA MADS-box genes. Nature. 2000;405(6783):200-3. |
| stage 1 | 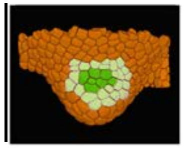 | 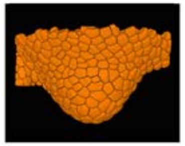 | 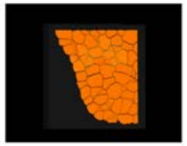 | 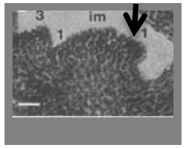 | 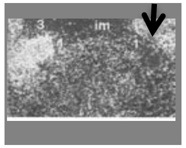 | | | |
| stage 2 | 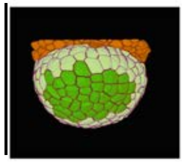 | 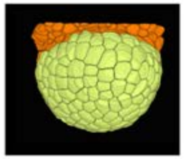 | 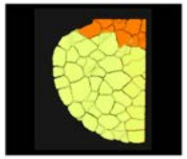 | 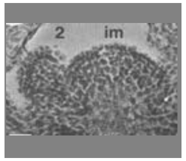 | 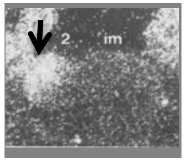 | | | |
| stage 3 | 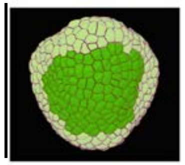 | 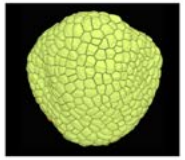 | 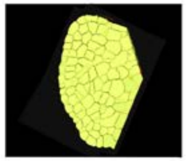 | | | | | |
| stage 4 | 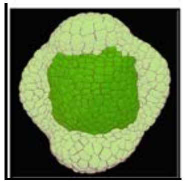 | 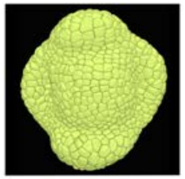 | 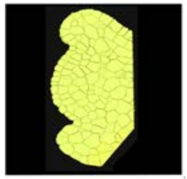 | 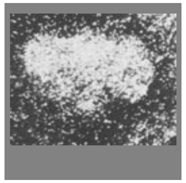 | 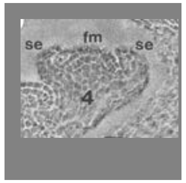 | | | |

Remarks: Only one clear expression pattern found in the litterature.

### SEPALLATA2

| | Stages of flower development with future meristem and sepals marked | top view | section | Savidge et al 1995 | References |
|---|---|---|---|---|---|

Initium stage

stage 1

stage 2

stage 3

stage 4

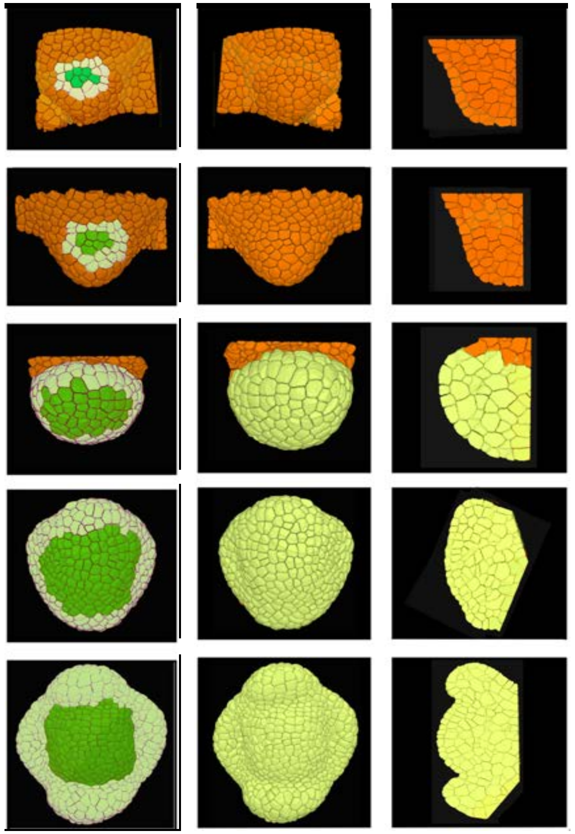

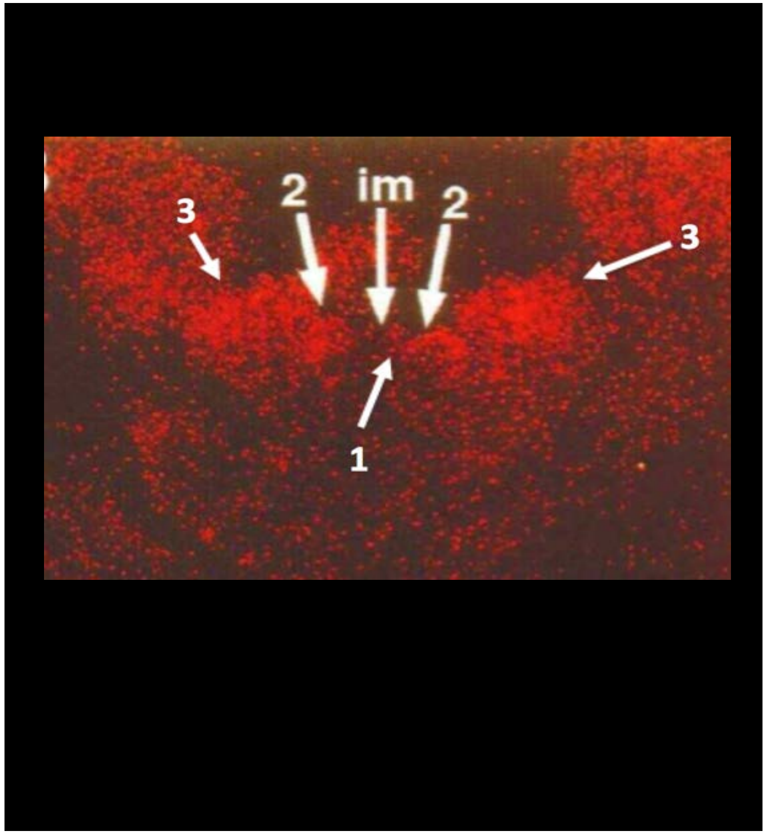

1. Krizek BA, Fletcher JC. Molecular mechanisms of flower development: an armchair guide. Nat Rev Genet. 2005;6(9):688-98.
2. Pelaz S, Ditta GS, Baumann E, Wisman E, Yanofsky MF. B and C floral organ identity functions require SEPALLATA MADS-box genes. Nature. 2000;405(6783):200-3.
3. Savidge B, Rounsley SD, Yanofsky MF. Temporal relationship between the transcription of two Arabidopsis MADS box genes and the floral organ identity genes. Plant Cell. 1995;7(6):721-33.

Remarks: Only one image found, that indicates that SEP2 expression is very similar if not identical to the pattern of SEP1 (see also summary of expression patterns in Krizek and Fletcher (2005).

### SEPALLATA3/AGL9

Columns: Stages of flower development with future meristem and sepals marked | top view | section | Mandel et al 1998 | Urbanus et al 2009 | Vachez et al unpublished | Vachez et al unpublished

Rows: Initium stage | stage 1 | stage 2 | stage 3 | stage 4

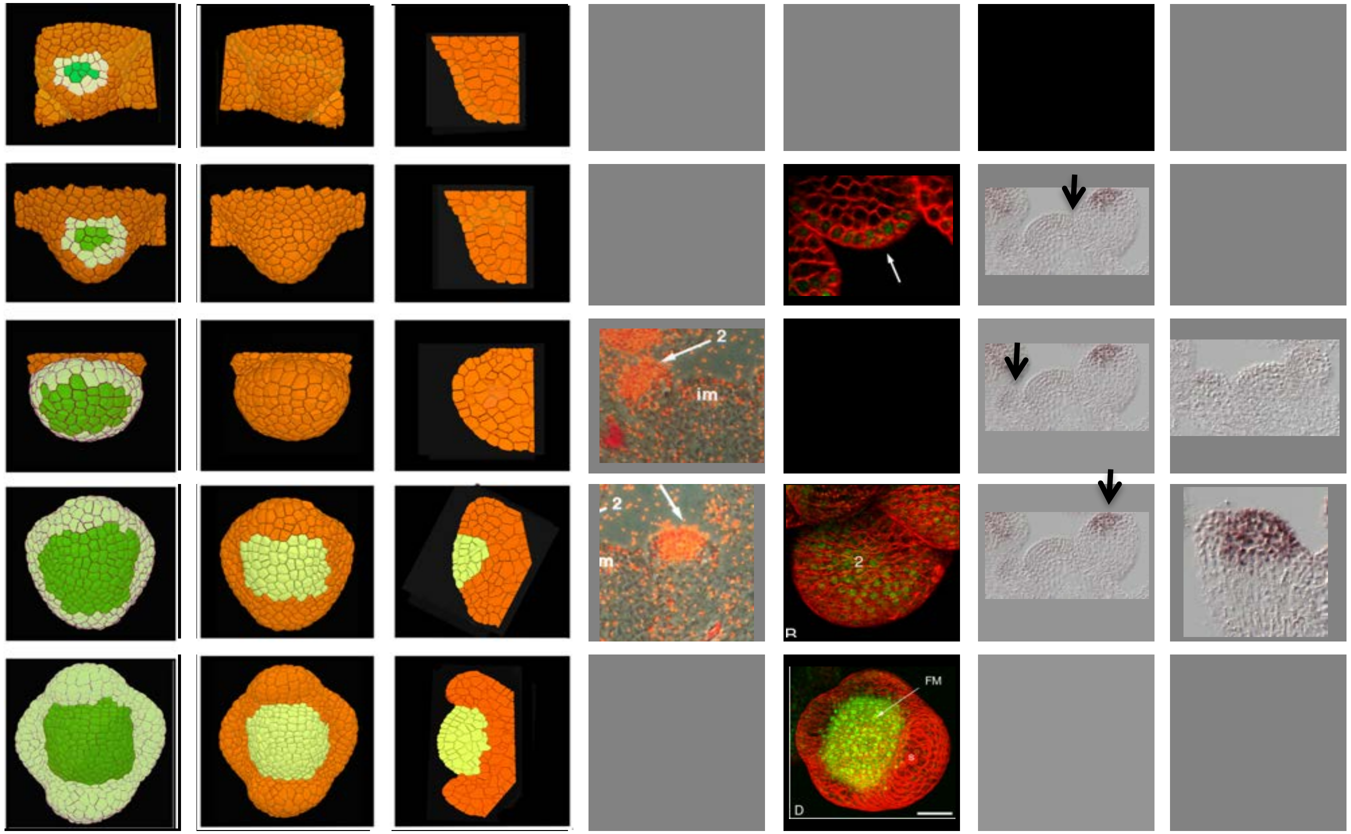

### SHOOTMERISTEMLESS

| | Stages of flower development with future meristem and sepals marked | topview | section | Long and Barton, 2000 | Maier et al 2011 | Landrein et al 2015 | | References |
|---|---|---|---|---|---|---|---|---|
| Initium stage | | | | | | | | |
| stage 1 | | | | | | | | |
| stage 2 | | | | | | | | |
| stage 3 | | | | | | | | |
| stage 4 | | | | | | | | |
| Remarks: | In particular the study by Barton and Long is very detailed. According to the serial sections (not reproduced here), STM reactivates during stage 1 (labelled 5 on the figure) in the future meristem and not in the cryptic bract. The reactivation at this stage is also confirmed by the GFP line used by Landrein et al. The gene is also clearly not expressed in the future sepal primordia at early stage 3. STM is not active in the lateral sepals, even before they grow out and AHP6 is active. | | | | | | | |

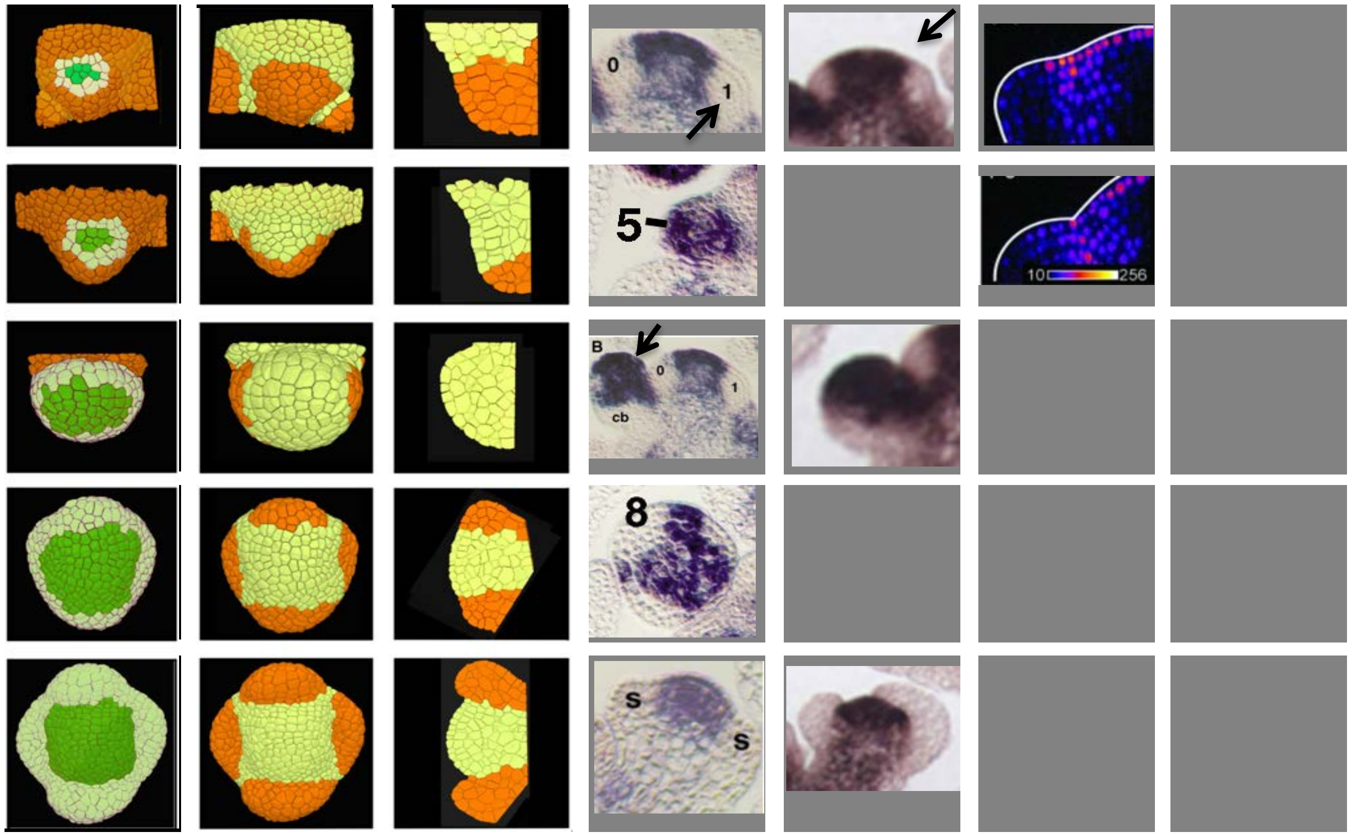

1. Landrein B, Kiss A, Sassi M, Chauvet A, Das P, Cortizo M, et al. Mechanical stress contributes to the expression of the STM homeobox gene in Arabidopsis shoot meristems. Elife. 2015;4:e07811.
2. Long J, Barton MK. Initiation of axillary and floral meristems in Arabidopsis. Dev Biol. 2000;218(2):341-53.
3. Maier AT, Stehling-Sun S, Offenburger SL, Lohmann JU. The bZIP Transcription Factor PERIANTHIA: A Multifunctional Hub for Meristem Control. Front Plant Sci. 2011;2:79.

### SUPERMAN

| | Stages of flower development with future meristem and sepals marked | top view | section | itoh et al 2003 | Prunet et al 2017 | References |
|---|---|---|---|---|---|---|
| Initium stage | | | | | | |
| stage 1 | | | | | | |
| stage 2 | | | | | | |
| stage 3 | | | | | | |
| stage 4 | | | | | | |

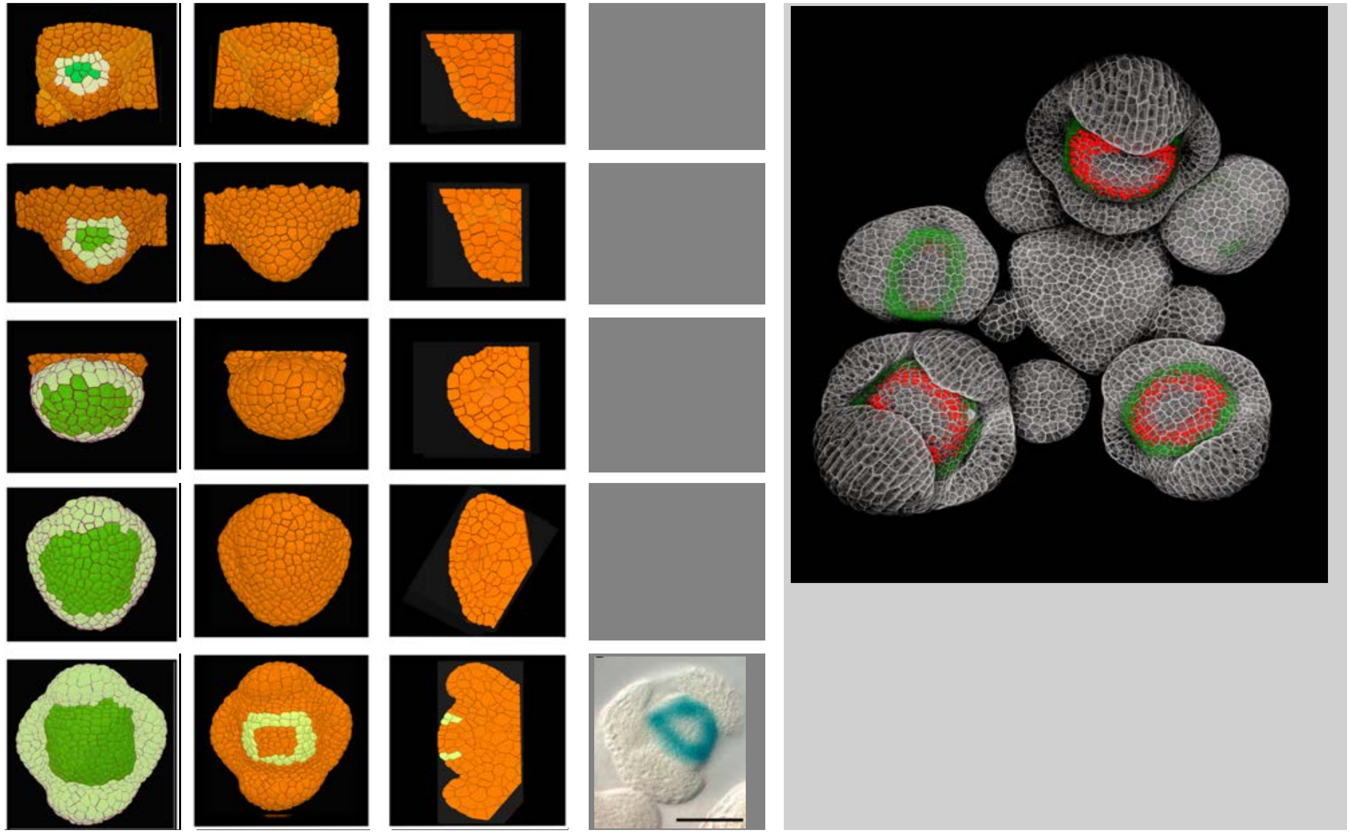

Ito, T., et al. (2003). "Whorl-specific expression of the SUPERMAN gene of Arabidopsis is mediated by cis elements in the transcribed region." Curr Biol 13(17): 1524-1530. Pru N., Jack, T.P., and Meyerowitz, E.M. (2016). Live confocal imaging of Arabidopsis flower buds. Dev. Biol. 419, 114–120.

Remarks: The publication by Ito et al (2003) described an expression pattern somehwere between the third and fourth whorl. The line made by Prunet al shows a partial overlap between AP3-GFP (green) and SUP-VENUS N7 (red).

### Short Vegetative Phase

| | Stages with approx. zones of future meristem (green) and sepals (yellow) marked | SVP top view | SVP section | Hartmann et al 2000 | Liu et al 2013 | Liu et al 2007 |
|---|---|---|---|---|---|---|
| Initium stage | 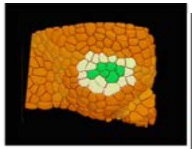 | 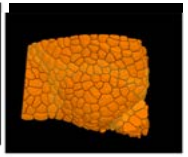 | 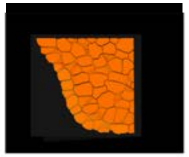 | | | |
| stage 1 | 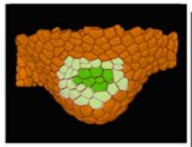 | 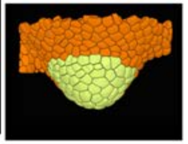 | 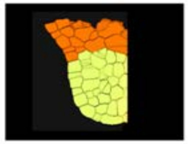 | 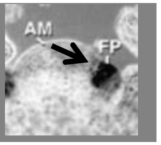  | 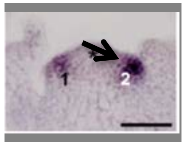  | 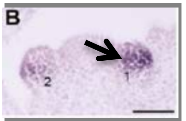  |
| stage 2 | 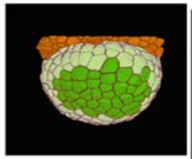 | 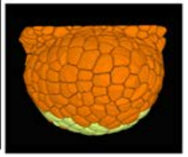 | 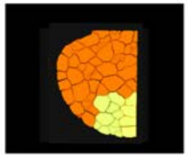 | | | 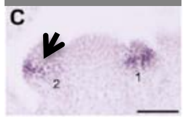  |
| stage 3 | 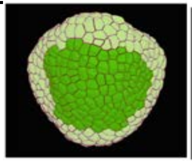 | 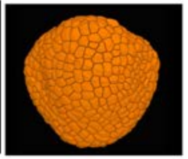 | 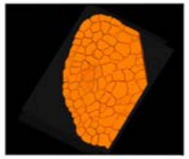 | 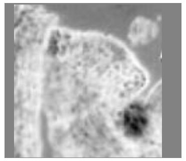 | | |
| stage 4 | 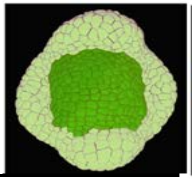 | 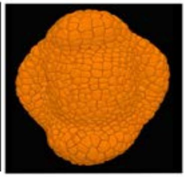 | 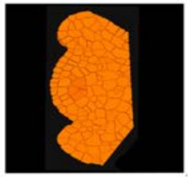 | | | 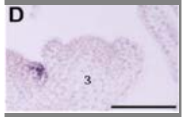  |
| Remarks: | SVP is expressed during the early stages of flower development and then gradually fades, remaining in the bract domain until stage 2. | | | | | |

### WUSCHEL

Stages of flower development with future meristem and sepals marked | top view | section | Maier et al | Rozier et al | Mayer et et al | Lenhard et al

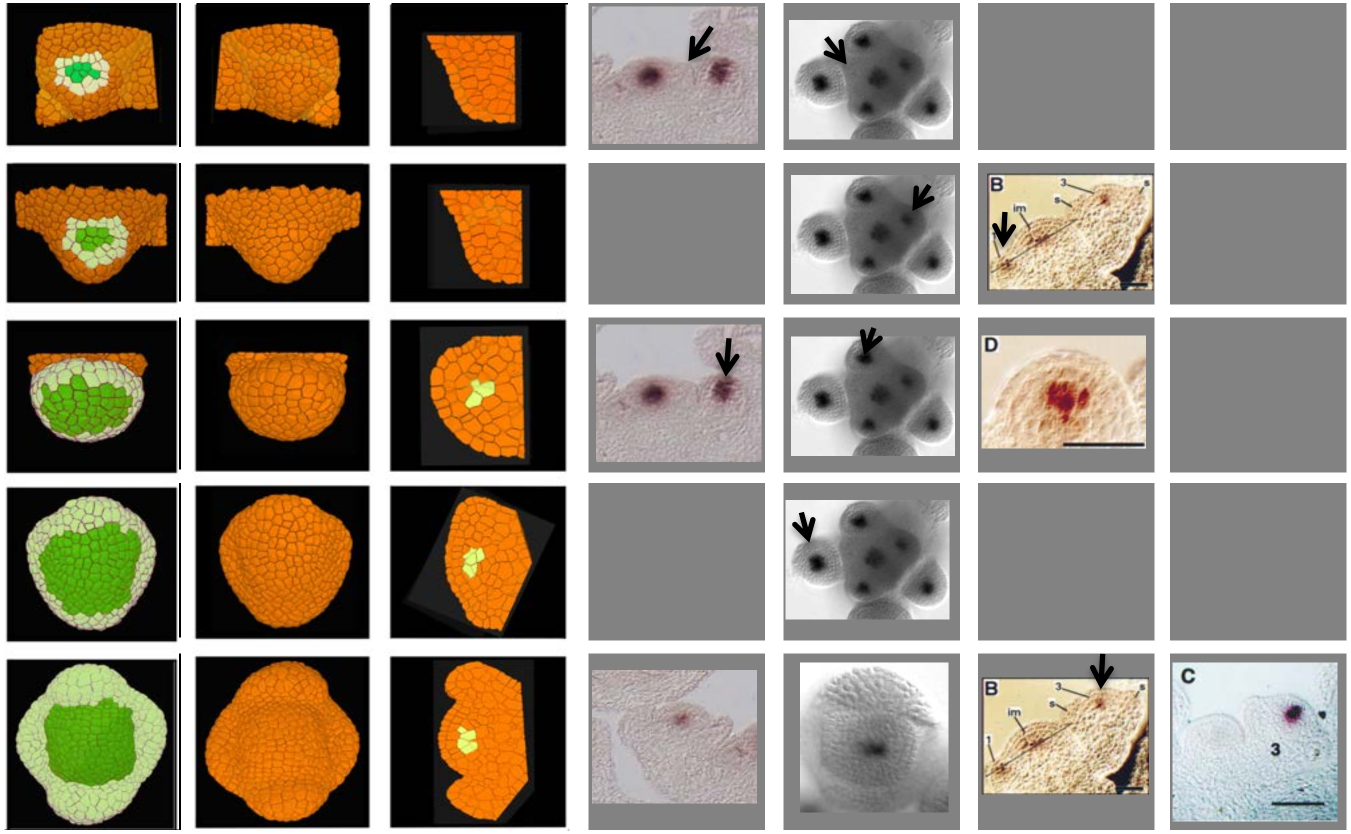

Remarks: In the flower the expression of WUS is limited to the L3. The gene is activated during stage 1 : note relatively weak expression in images from Rozier et al.

**How to use morphonet?**

1. Connect to:  www.morphonet.org

2. Login:

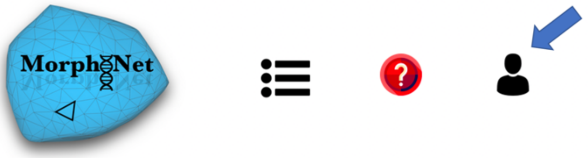

click on this icon

**login:** ReviewerAtlas **password:** review

3. Click on 'Arabidopsis thaliana'

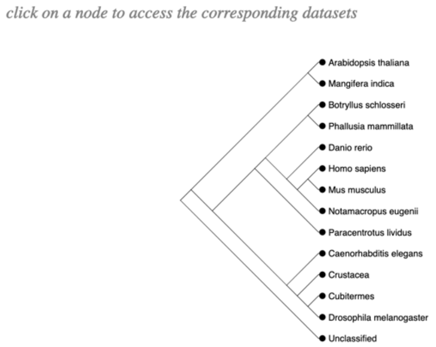

4. Click on one of the following datasets ( )

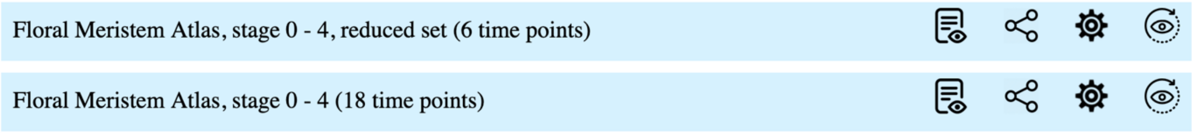

5. For visualization using Mac:

- 'control' key + mouse to rotate the meristem
- 'option' key + mouse to move it up down, right or left
- use scroll bar on Morphonet menu to go from one time point to the other 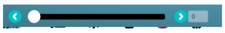
- to visualize the inner cells, go to 'Dataset' menu and use 'crop' function
- **if screen resolution is insufficient not all information might be visible or accessible. In that case type F (capital) to go on full screen.**

6. To visualize cell lineage

- click on 1 cell and then move the scroll bar to another time point

7. To visualize expression patterns

- Go to (i) time point 1-5 in the reduced set or (ii) go to time point 1, 5, 12, 15 or 17 in the complete set.
- In the 'genetic' menu click on one or more of the colored squares next to the gene names
